# A-ring-fused pyrazoles of dihydrotestosterone targeting prostate cancer cells via the downregulation of the androgen receptor

**DOI:** 10.1101/2022.12.17.520850

**Authors:** Miroslav Peřina, Márton A. Kiss, Gergő Mótyán, Eva Szczyrbová, Martin Eliáš, Vladimír Študent, Daniela Kurfurstová, Markéta Kovalová, Lukáš Mada, Jan Bouchal, Éva Frank, Radek Jorda

## Abstract

High expression of the androgen receptor (AR) and the disruption of its regulation are strongly responsible for the development of prostate cancer (PCa). Therapeutically relevant non-steroidal or steroidal antiandrogens are able to block the AR effect by eliminating AR-mediated signalling. Herein we report the synthesis of novel steroidal pyrazoles derived from the natural sex hormone 5α-dihydrotestosterone (DHT). 2-Ethylidene or 2-(hetero)arylidene derivatives of DHT obtained by regioselective Claisen-Schmidt condensation with acetaldehyde or (hetero)aromatic aldehydes in alkaline ethanol were reacted with monosubstituted hydrazines to give A-ring-fused 1,5-disubstituted pyrazoles as main or exclusive products, depending on the reaction conditions applied. Spontaneous or 2,3-dichloro-5,6-dicyanobenzoquinone (DDQ)-induced oxidation of the primarily formed pyrazolines resulted in the desired products in moderate to good yields, while 17-oxidation also occurred by using the Jones reagent as a strong oxidant. Transcriptional activity of the AR in a reporter cell line was examined for all novel compounds, and several previously synthesized similar DHT-based pyrazoles with differently substituted heteroring were also included to obtain information about the structure-activity relationship. Two specific regioisomeric groups of derivatives significantly diminished the transcriptional activity of AR in reporter cell line in 10 μM concentration, and displayed reasonable antiproliferative activity in AR-positive PCa cell lines. Lead compound (**3d**) was found to be a potent AR antagonist (IC_50_ = 1.18 μM), generally suppressed AR signalling in time and dose dependent manner, moreover, it also led to a sharp decrease in wt-AR protein level probably caused by proteasomal degradation. We confirmed the antiproliferative activity selective for AR-positive PCa cell lines (with GI_50_ in low micromolar ranges), cellular, biochemical and in silico binding of **3d** in AR ligand-binding domain. Moreover, compound **3d** was shown to be potent even *ex vivo* in patient-derived tissues, which highlights the therapeutic potential of A-ring-fused pyrazoles.

**Table of content graphic:** 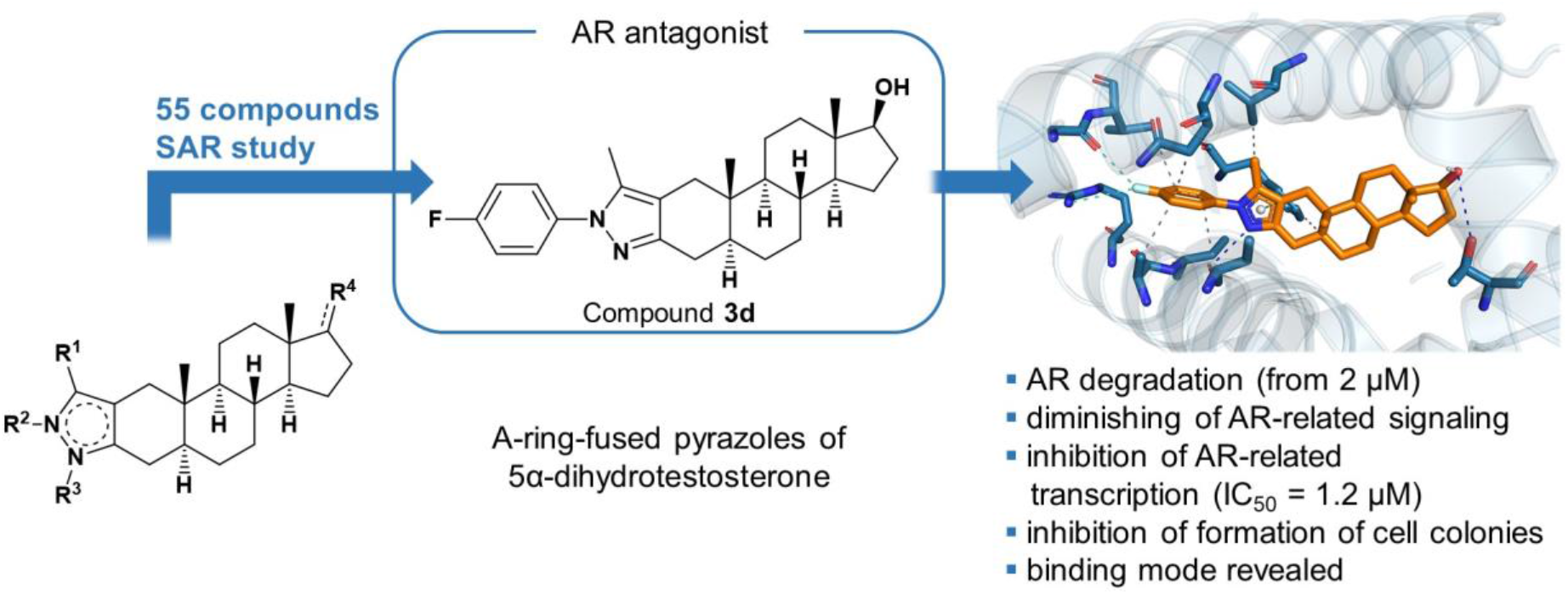

**Highlights:**

- A-ring-fused pyrazoles of 5α-dihydrotestosterone were introduced as AR modulators
- SAR study of 55 differently substituted derivatives was discussed
- Compound **3d** suppressed the expression of AR targets in prostate cell lines
- Compound **3d** caused AR degradation in time-dependent manner
- Ex vivo activity of **3d** was demonstrated in patient-derived explants

## 1. Introduction

The androgen receptor (AR) is a ligand-activated transcription factor that belongs to the superfamily of steroid and thyroid hormone receptors and plays a crucial role in the normal development of male reproductive tissues. Androgen binding induces conformational changes of the AR that influences its interactions with other proteins and DNA, as well as its subcellular localization and transcriptional activity. The AR is further regulated by numerous post-translational modifications that affect its physiological role, especially its transcriptional program [1,2].

High expression and/or relaxation of AR regulation is strongly implicated in prostate cancer (PCa). PCa is the second most frequent cancer in men according to the National Cancer Institute (USA). Primary therapy (androgen deprivation therapy) is based on reduction of the circulating androgens in plasma (by orchiectomy or with luteinizing hormone-releasing hormone agonists), but usually the disease progresses to castration-resistant prostate cancer (CRPC) stage, which is defined by alteration in signalling of AR (high expression, splicing variants) and resistance to standard therapeutics. Therefore, AR has been suggested as an important therapeutic target for PCa, and several anti-AR strategies have been introduced for decades including inhibition of the transcription of the AR gene, inhibition or destabilization of transcript or protein or splicing variants, AR degradation, blocking of AR synthesis, interference with intracellular trafficking of AR, and inhibition of downstream signalling related to AR-V7 activation (**Figure 1**) [3,4].

**Figure 1.**
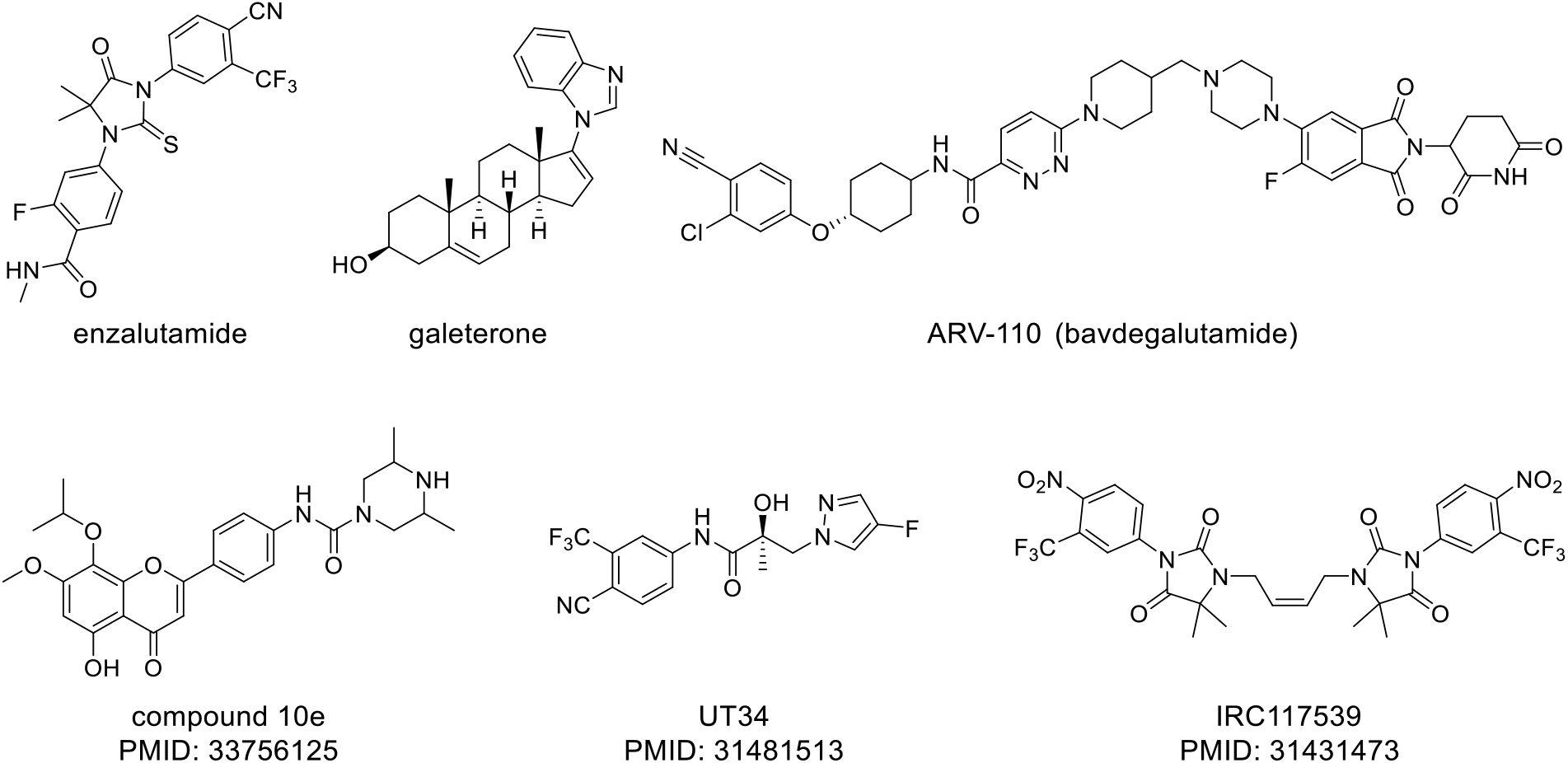
Examples of AR modulators in clinical trials (upper panel) and compounds with the ability to degrade AR (bottom panel).

Several steroidal or non-steroidal agents (e.g. abiraterone, enzalutamide, apalutamide, darolutamide) with diverse modes of action (CYP17 hydroxylase/17,20 lyase inhibitors (CYP17), AR antagonists) have been approved for CRPC, and data have demonstrated an overall survival benefit also in the castration-sensitive stage. On the other hand, several AR-related mechanisms of resistance have been described, and novel strategies to overcome them are an important unmet need. Since recent studies have revealed that several mutations in the AR converts the action of enzalutamide and apalutamide from antagonist to agonist [5,6], it appears suitable to target AR also indirectly, e.g. by AR-destabilizing agents. Decrease in AR protein stability accompanied by increased degradation, which is connected with its short half-life (highlighted in the presence of no androgens) and by the presence of PEST sequence responsible for rapid degradation in proteasome. Several compounds with AR degradation activity have been introduced including hybrid molecules such as PROTACs (proteolysis-targeting chimeras based on proteasome-mediated degradation of protein of interest) [7–10] or SNIPERs (specific and nongenetic inhibitor of apoptosis protein-dependent protein erasers [11]. Interestingly, biopharmaceutical company Arvinas developed several AR degraders, and currently ARV-110 (bavdegalutamide) is the first degrader in Phase 2 clinical trial for the treatment of patients with metastatic CRPC [12,13].

Sterane-based D-ring-modified pyrazoles [14,15], structurally related to abiraterone, are known to be potentially effective antiandrogen agents for the treatment of PCa by antagonizing AR, acting as CYP17 enzyme inhibitors and/or having direct cytotoxicity. However, steroids containing a pyrazole moiety in the A-ring were less investigated. We previously described some A-ring-fused arylpyrazoles of DHT and demonstrated their anticancer activity against multiple cancer cell lines including PCa cells [16,17]. Cyclocondensation reactions of 2-hydroxymethylene-DHT led to a mixture of separable pyrazole regioisomers (**1** and **2** series) in varying ratio depending on the applied medium and on the electronic character of the substituent of the phenylhydrazine applied [16]. Similar derivatives containing a 1,5-disubstituted pyrazole moiety (**3** and **4** series) were also obtained (**Figure 2**) [17].

**Figure 2.**
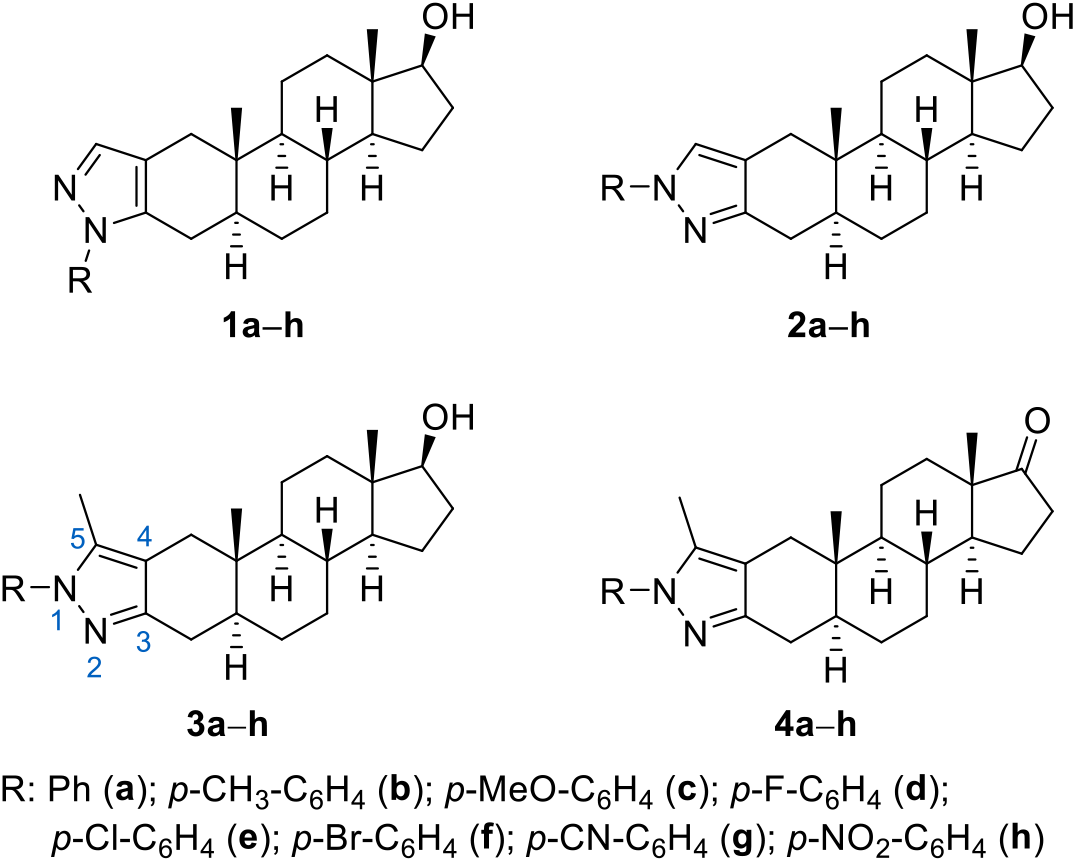
A-ring-fused pyrazoles of DHT previously synthetized by us [16,17].

In the current work, additional 1,5-disubstituted pyrazoles structurally similar to **3** and **4** have been prepared, and a total of 55 compounds including the previously and newly synthesized ones were screened for their ability to affect the transcriptional activity of AR in a reporter cell line. The structure of all novel compounds was determined by 1D and 2D NMR spectroscopy as well as electrospray ionization mass spectrometry (ESI-MS). All compounds were analysed for their agonist and antagonist activity towards the AR, for their ability to decrease AR protein level and to block the formation of LAPC-4 colonies. Based on the obtained results from screening, SAR for compounds was described, and lead compound **3d** was further studied in detail. Lead compound exerted reasonable antiproliferative activity towards AR-positive PCa cell lines and inhibited the colony formation. It was demonstrated that **3d** dose-dependently diminishes AR transcriptional activity and mRNA expression of PSA (key downstream target). Moreover, lead compound markedly decreased the AR protein level and totally turned off downstream signalling upon longer treatment. Molecular docking proved the possibility of binding **3d** into the wt-AR ligand binding domain, but also into frequent Thr877Ala mutant with extensive interactions.

## 2. Results and Discussion

### 2.1. Synthesis and characterization of the target compounds

In order to synthesize compounds structurally similar to those from series **3**, arylidene derivatives (**5a–j**) of DHT were first synthesized by Claisen-Schmidt condensation as α,β-enone precursors suitable for heterocyclization (Supporting Information, **Scheme S1**). The majority of the compounds were obtained by the method described previously [18], however, two additional molecules (**5f** and **5i**) were also prepared on the basis of preliminary flexible docking into AR and solubility considerations. As a next step, the cyclization of the prepared enones (**5**) with methylhydrazine as binucleophilic reagent was planned to be carried out. Preliminary experiment was performed with **5a** in EtOH using MW irradiation. After an irradiation time of 3 min, complete conversion was observed by thin-layer chromatography (TLC), and four new spots were detected on the silica plate. The products were separated by column chromatography, and their structures were determined by NMR spectroscopy. Two of the four compounds proved to be pyrazole regioisomers **8a** and **9a** formed by heterocyclization and subsequent autooxidation under the reaction conditions, while the other two products were the diastereomeric pairs of the pyrazoline precursor **6a** of the major heteroaromatic product **8a** (**Scheme 1**). The order of elution in descending polarity was as follows: **9a** (1,3-pyrazole) > **8a** (1,5-pyrazole) > **6a** (inseparable mixture of two isomers). These results thus showed that a considerable amount of the pyrazolines formed primarily by the ring-closure underwent oxidation under the reaction conditions. The partially saturated precursor **7a** of the 1,3-disubstituted pyrazole **9a** could not be detected.

**Scheme 1.**
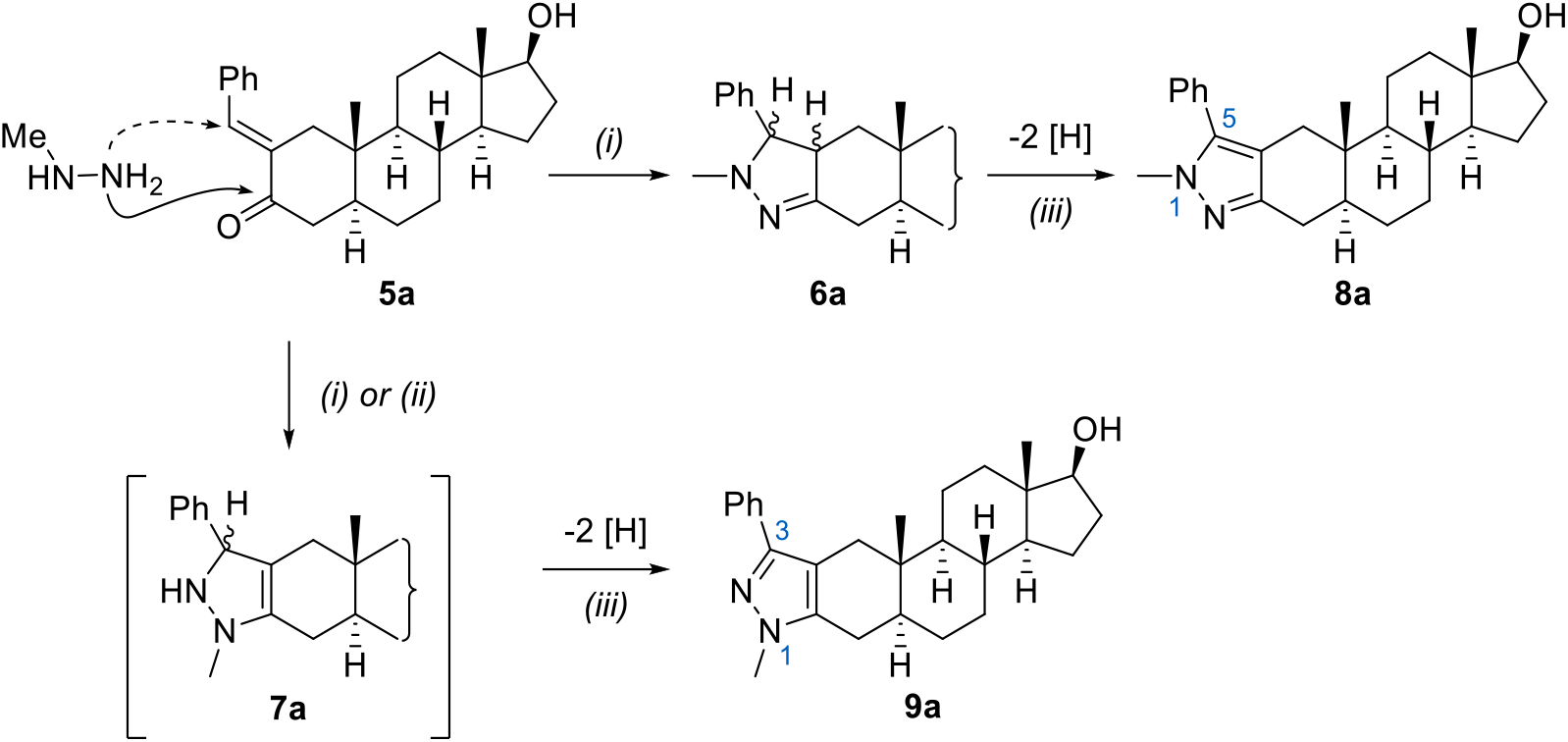
Synthesis of ring A-fused disubstituted pyrazole regioisomers of DHT. *Reagents and conditions: (i)methylhydrazine (2.0 equiv.), EtOH, MW, 100 °C, 3 min; (ii) methylhydrazine sulfate (2.0 equiv.), EtOH, reflux, 24 h; (iii) DDQ, dioxane, 120 °C, 5 min (**8a**: 57%, **9a**: 28%) or spontaneous oxidation (**8a**:80%)*

Since the reaction described above led to the formation of four difficult-to-separate products, an additional oxidation step was introduced before purification to complete the spontaneous oxidation observed partially, and thus reduce the number of products to two. 2,3-Dichloro-5,6-dicyanobenzoquinone (DDQ), which has been successfully used previously for the oxidation of the heteroring without affecting the C17-OH bond [18] was suitable for this purpose. The reaction was first carried out in 1,4-dioxane with conventional heating, which required a relatively large amount of hot solvent to dissolve the starting materials. Although the oxidation was completed within 2 h, the work-up procedure and extraction of the products proved to be difficult. However, when the reaction was carried out in a MW reactor, the higher reaction temperature (120 °C) in a closed vessel allowed the starting materials to dissolve in significantly less solvent (5 mL), and a much shorter time (5 min) was required for the DDQ-induced oxidation. The use of a smaller amount of solvent facilitated the work-up, as a filterable precipitate was observed when the reaction mixture was poured onto water. As expected, the oxidation resulted in a mixture of the two possible pyrazole regioisomers (**8a** and **9a**) in a ratio of about 2:1, which were easily separable by column chromatography. However, the heterocyclization of **5a** with methylhydrazine sulfate under conventional heating in EtOH for 24 h led to the exclusive formation of the 1,5-pyrazole (**8a**) without the traces of its pyrazoline precursor (**6a**) or the other heteroaromatic regioisomer (**9a**) (**Scheme 1**). Since the formation of **9a** by initial 1,4-addition of methylhydrazine to **5a** is under kinetic control, short reaction time under MW heating can lead to the 1,3-disubstituted product in a minor extent by rapid dehydrogenation of the cyclized pyrazoline precursor **7a**. Contrarily, longer reaction time at reflux temperature favoured the exclusive formation of the thermodynamically more stable 1,5-regioisomer (**8a**) by nucleophilic attack of the reagent on the carbonyl-C followed by cyclization and spontaneous oxidation [19].

The structure of the regioisomeric pyrazoles **8a** and **9a** was confirmed by 1D NMR (^1^H and ^13^C NMR, Supporting Information) and 2D NMR measurements (HSQC, HMBC, NOESY). A significant difference between the two spectra can be observed in the range of 2.00–3.00 ppm and above 7.00 ppm in the aromatic region, where the peaks of the protons located at or near to the reaction centre appear. The characteristic signals of the hydrogens at C-1 and C-4 are found at a chemical shift above 2.00 ppm due to the proximity of the heteroaromatic ring. For **8a** and **9a**, the reversed order of the signals indicates a structural difference between the compounds. In order to identify the 1,3- and the 1,5-pyrazoles, their 2D-NMR spectra were also recorded, and after determining the characteristic correlations, we also examined the NOESY spectra of the products, which provided information on the protons with the same spatial arrangement (**Figures S1** and **S2**).

As mentioned above, the heterocyclization led to the regioselective formation of the 1,5-regioisomer (**8a**) in good yield under conventional heating without the need for an additional oxidation step (**Scheme 1**, **Table 1**, entry 1). As a continuation, further arylidene derivatives (**5b–h**) were subjected to ring-closure, and the corresponding 1,5-regioisomers (**8b–h**) were obtained in moderate to good yields (**Table 1**, entries 2–8). For compounds bearing a halogen substituent on the benzene ring, the oxidation was completed using DDQ, while spontaneous oxidation occurred in other cases. Treating the crude products with the Jones reagent in acetone, besides the pyrazoline ring, the 17β-hydroxyl group was also oxidized leading to the corresponding 17-ones (**10b–h**) (**Table 1**, entries 1–8). Furthermore, the 1,5-dimethyl-substitued pyrazole (**8l**) and its 17-one derivative (**10l**) were also prepared from the previously synthesized ethylidene derivative **5j** [17]. Based on promising preliminary flexible docking studies for diaryl-substituted pyrazoles, oxidative heterocyclization of **5a**, **5e** and **5i** was carried out with phenylhydrazine hydrochloride under MW condition (**Table 1**, entries 9–11), leading to compounds **8i–k** in high yields. Their D-ring-oxidized analogues **10i–k** were also prepared for pharmacological comparison.

**Table 1.**
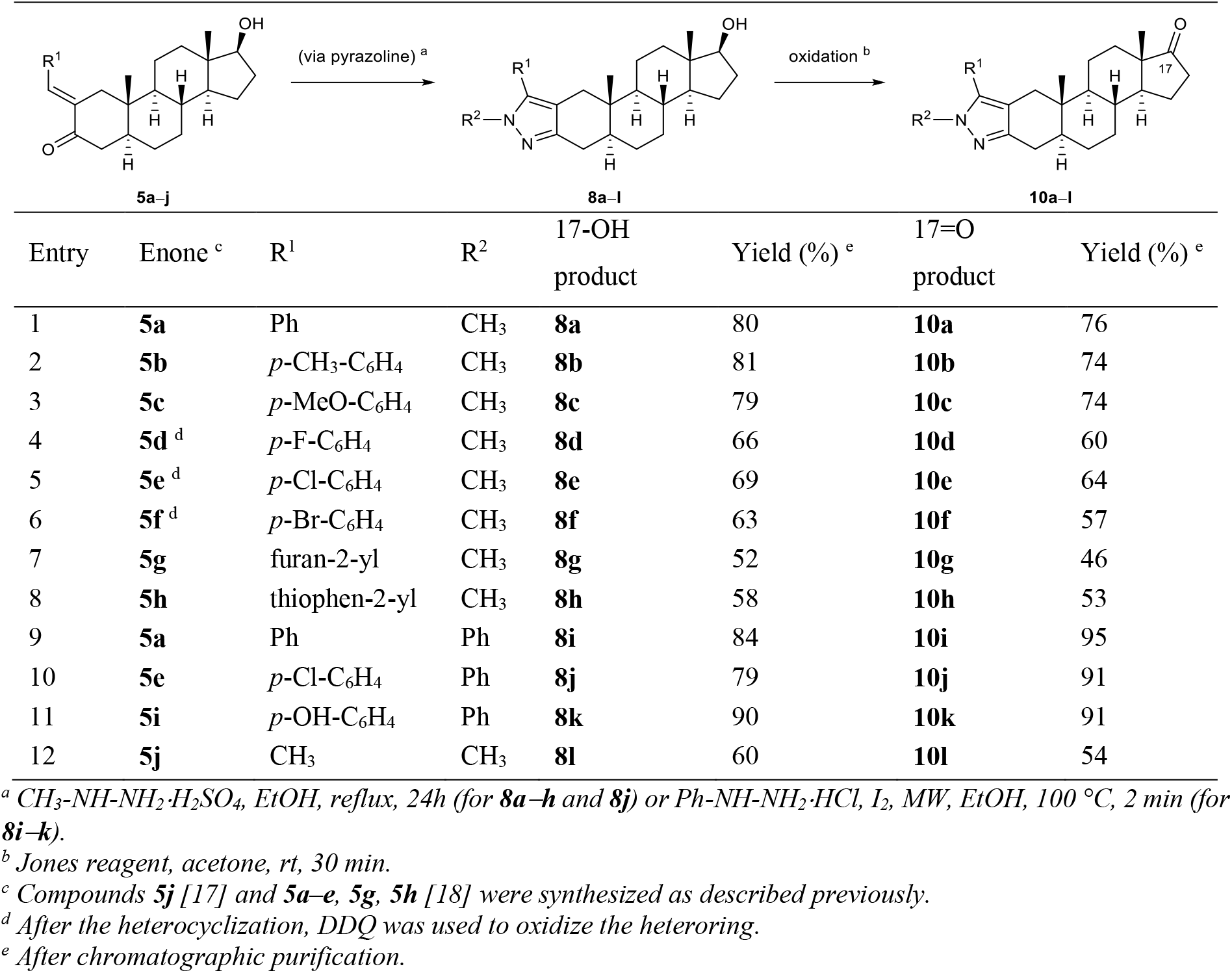
Synthesis of DHT-derived A-ring-fused 1’,5’-pyrazole derivatives

| Entry | Enone <sup>c</sup> | R<sup>1</sup> | R<sup>2</sup> | 17-OH product | Yield (%) <sup>e</sup> | 17=O product | Yield (%) <sup>e</sup> |
| --- | --- | --- | --- | --- | --- | --- | --- |
| 1 | **5a** | Ph | CH<sub>3</sub> | **8a** | 80 | **10a** | 76 |
| 2 | **5b** | *p*-CH<sub>3</sub>-C<sub>6</sub>H<sub>4</sub> | CH<sub>3</sub> | **8b** | 81 | **10b** | 74 |
| 3 | **5c** | *p*-MeO-C<sub>6</sub>H<sub>4</sub> | CH<sub>3</sub> | **8c** | 79 | **10c** | 74 |
| 4 | **5d** <sup>d</sup> | *p*-F-C<sub>6</sub>H<sub>4</sub> | CH<sub>3</sub> | **8d** | 66 | **10d** | 60 |
| 5 | **5e** <sup>d</sup> | *p*-Cl-C<sub>6</sub>H<sub>4</sub> | CH<sub>3</sub> | **8e** | 69 | **10e** | 64 |
| 6 | **5f** <sup>d</sup> | *p*-Br-C<sub>6</sub>H<sub>4</sub> | CH<sub>3</sub> | **8f** | 63 | **10f** | 57 |
| 7 | **5g** | furan-2-yl | CH<sub>3</sub> | **8g** | 52 | **10g** | 46 |
| 8 | **5h** | thiophen-2-yl | CH<sub>3</sub> | **8h** | 58 | **10h** | 53 |
| 9 | **5a** | Ph | Ph | **8i** | 84 | **10i** | 95 |
| 10 | **5e** | *p*-Cl-C<sub>6</sub>H<sub>4</sub> | Ph | **8j** | 79 | **10j** | 91 |
| 11 | **5i** | *p*-OH-C<sub>6</sub>H<sub>4</sub> | Ph | **8k** | 90 | **10k** | 91 |
| 12 | **5j** | CH<sub>3</sub> | CH<sub>3</sub> | **8l** | 60 | **10l** | 54 |

<sup>a</sup> CH<sub>3</sub>-NH-NH<sub>2</sub>·H<sub>2</sub>SO<sub>4</sub>, EtOH, reflux, 24h (for **8a–h** and **8j**) or Ph-NH-NH<sub>2</sub>·HCl, I<sub>2</sub>, MW, EtOH, 100 °C, 2 min (for **8i–k**).
<sup>b</sup> Jones reagent, acetone, rt, 30 min.
<sup>c</sup> Compounds **5j** [17] and **5a–e**, **5g**, **5h** [18] were synthesized as described previously.
<sup>d</sup> After the heterocyclization, DDQ was used to oxidize the heteroring.
<sup>e</sup> After chromatographic purification.

### 2.2. Targeting AR and AR-related processes

We recently described the synthesis of DHT derivatives modified in the A-ring with (hetero)arylidene, pyrazolo[1,5-*a*]pyrimidine and triazolo[1,5-*a*]pyrimidine moieties and their targeting of the AR in PCa cell lines [18].

Currently we introduced novel DHT derivatives, namely A-ring-fused 1,5-disubstituted pyrazoles (**8**) and their C17-oxidized derivatives (**10**). These series were complemented (for better understanding of structure-activity relationship) with previously published A-ring-fused monosubstituted pyrazoles (**1** and **2**) and 1,5-disubstituted pyrazole derivatives of DHT (**3** and **4**) that demonstrated anticancer activity against multiple cancer cell lines including PCa cells, but were not pharmacologically investigated in relation to AR [16,17]. All compounds were investigated for their ability to affect i) the transcriptional activity of AR using an AR-dependent reporter cell line [20], ii) the formation of cell colonies, and iii) the expression of AR and its well-known AR-regulated genes (Nkx3.1 and PSA) via immunoblotting. All data are presented in **Tables 2–4**, **Table S1**, **Figure 3** and **Figures S3, S4, S5**.

**Figure 3.**
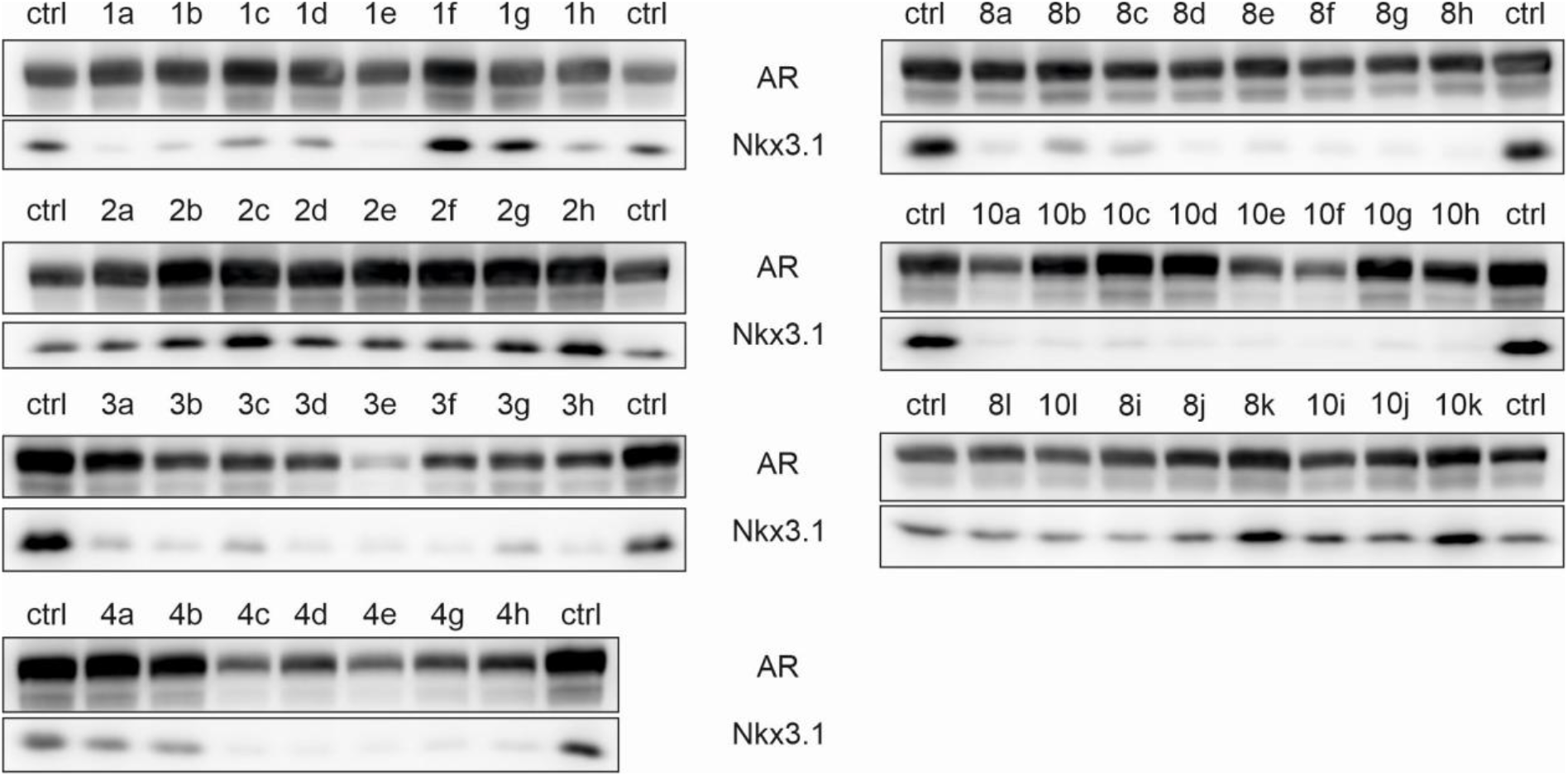
The LAPC-4 cells were treated with the studied steroids (10 μM, 24 h) in FBS containing medium and lysates were then blotted for detection of appropriate proteins. Representative results are shown; the expression of PSA and α-tubulin (loading control) are included in the Supporting Information, **Figure S3**.

**Table 2.**
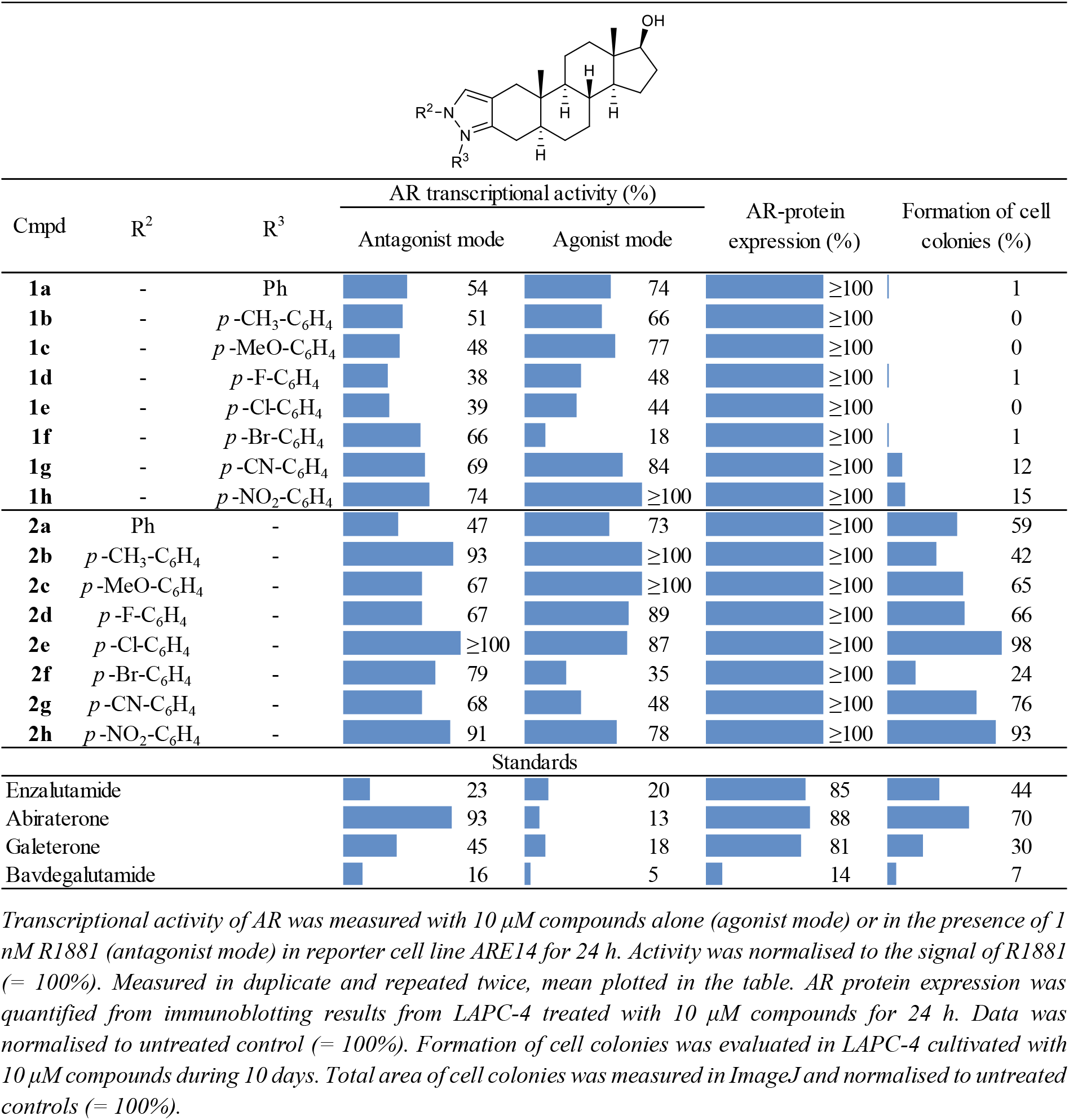
Investigated activities for N-monosubstituted pyrazole derivatives of DHT and selected standards.

**Table 3.**
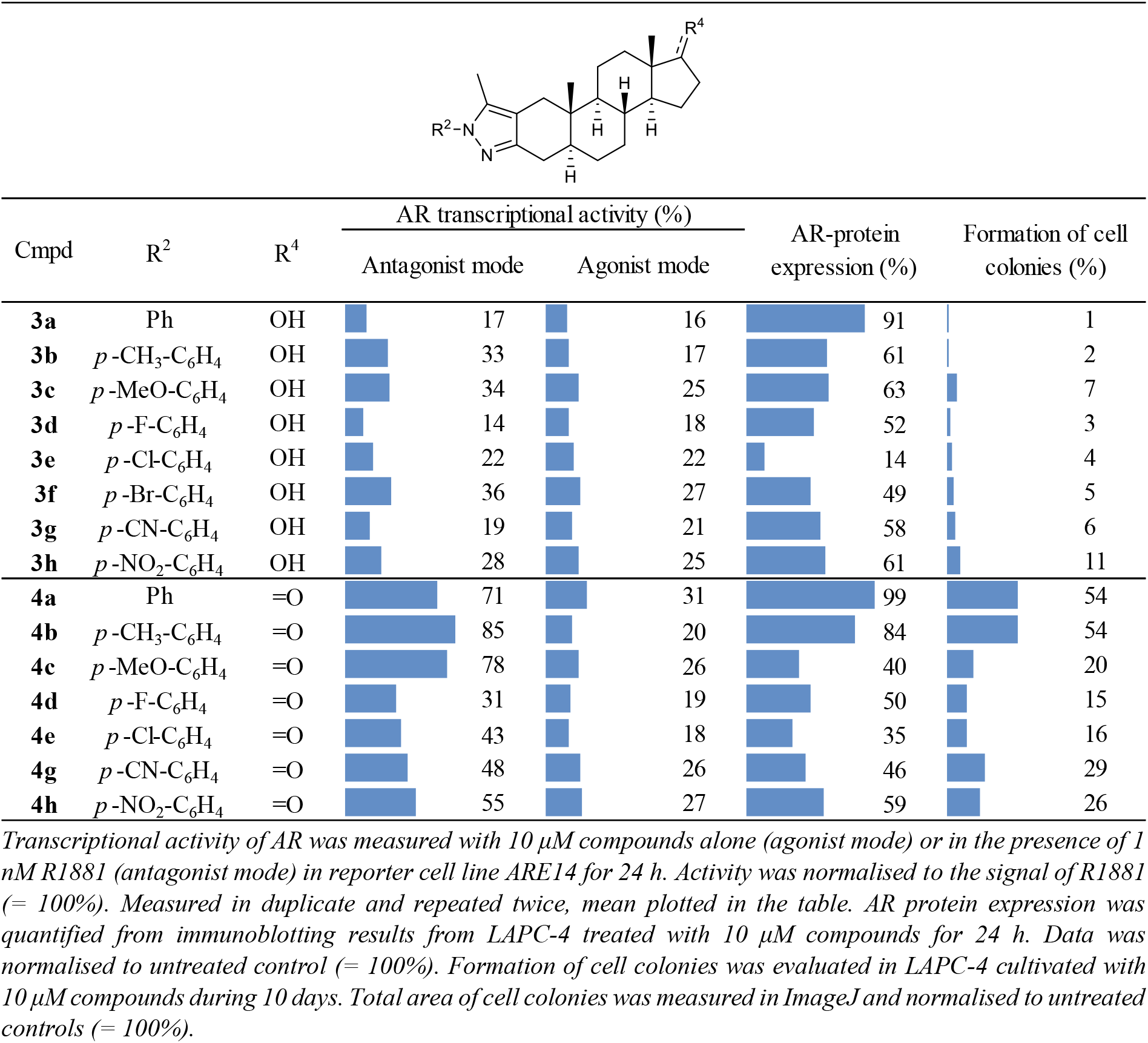
Investigated activities for 1-aryl-5-methyl pyrazole derivatives of DHT.

**Table 4.**
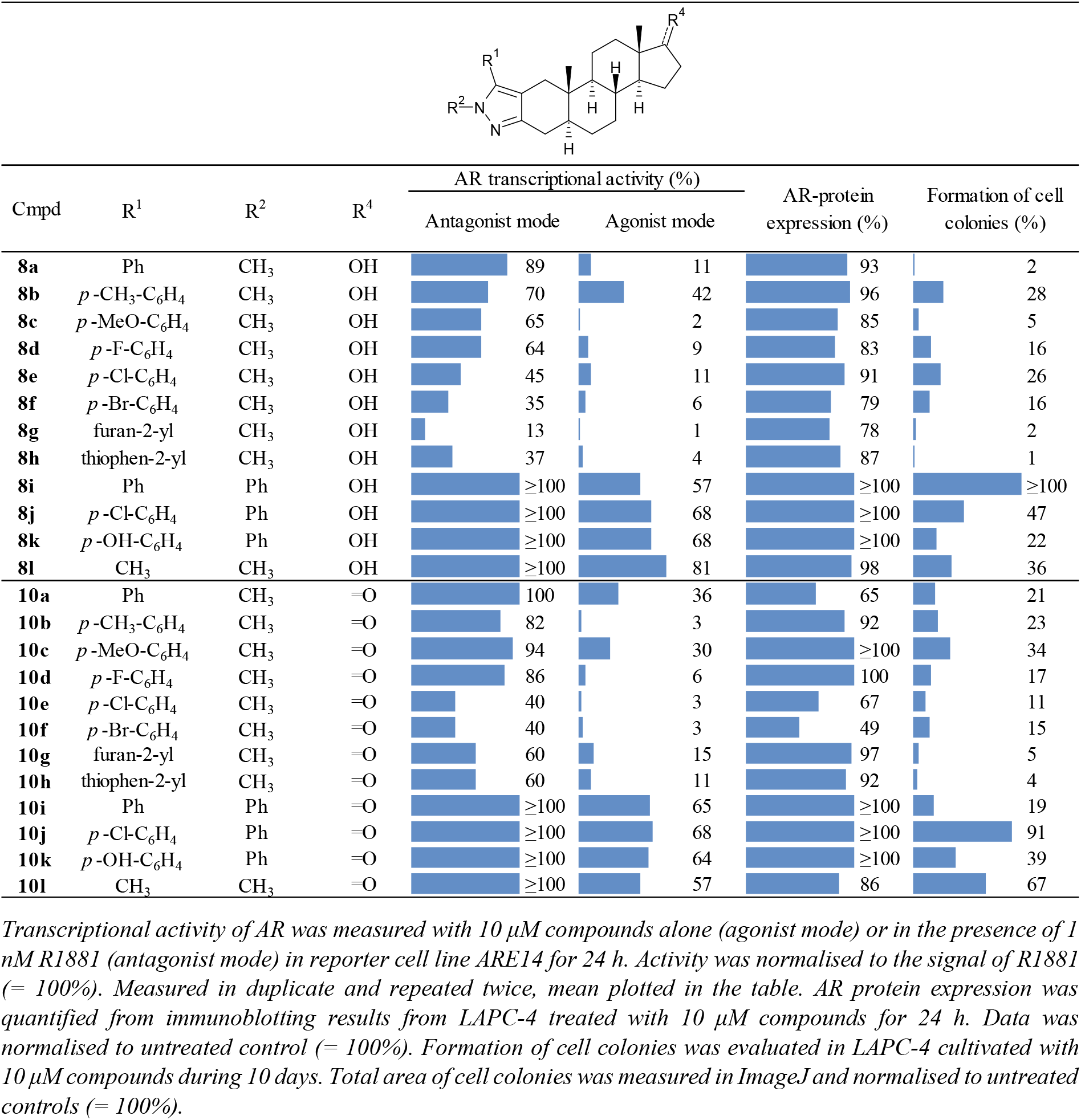
Investigated activities for 1,5-disubstituted pyrazole derivatives of DHT.

The ability of compounds to diminish the R1881-stimulated transcriptional activity of AR (antagonist mode) or to transactivate AR itself (agonist mode) was examined using an AR-dependent reporter cell line [20]. Compounds from series **1** and **2** displayed only moderate effect on R1881-stimulated AR-transcriptional activity (**Table 2**). Most of the compounds were not able to reach inhibition values ≤ 50% of control cells stimulated with 1 nM R1881 in 10 μM concentration. Only compounds **1d** and **1e** (2’-*p*-fluorophenyl and 2’-*p*-chlorophenyl-substituted pyrazole derivatives, respectively) showed inhibition value around 39%. Unfortunately, most compounds (except of regioisomers **1f** and **2f**) undesirably activated AR in agonist mode (**Table S1**).

Regioisomers of **3** and **8** that bear C-17 hydroxy group and combine methyl and aryl substitution at N1 or C5 position of pyrazole (**3a–3h**, **8a–8h**) showed to act as strong antagonists. In total, 12 from 16 compounds were able to decrease the AR-transcriptional activity below 50% of R1881-stimulated control (**Table 3, 4**). Compounds from series **3** were generally more potent antagonists than their regioisomers from series **8**. Importantly, none of compounds displayed agonist activities except of **8b**. The most potent steroids which were able to decrease the AR-transcriptional activity below 50% already in 2 μM concentration were **3a** bearing 5-methyl-1-phenyl pyrazole moiety, **3d** with 1-fluorophenyl-5-methyl pyrazole moiety and **3g** with 1-cyaonophenyl-5-methyl pyrazole group (**Table S1**).

Antagonist activities were observed also for series **4** and **10** bearing C-17 keto group (counterparts of **3** and **8,** combining methyl and aryl substitution at N1 or C5 position of the pyrazole ring, respectively). All compounds (**4a–4h**, **10a–10h)** acted predominantly as AR antagonists but were less potent than compounds from series **3** and **8**. Only 5 compounds were capable to decrease the AR-transcriptional activity below 50% of R1881-stimulated control (**Table 3, 4**).

Previous results showed that the combination of a small substituent (Me group) with a bulky one (Ph or modified *para*-substituted Ph moiety) in the 1,5 positions of the pyrazole ring is fundamental to reach strong antagonist activity. This scenario was obvious from the results with derivatives **8l**, **10l** (1,5-dimethyl-substituted) and **8i**, **10i** (1,5-diaryl-substituted) that displayed only agonist activities without any inhibition effect to R1881-stimulated AR transactivation.

Further, we evaluated the protein expression of AR and its transcriptional targets, PSA and Nkx3.1 (**Figures 3, S3**) in LAPC-4 cell treated with 10 μM concentration of compounds for 24 h. Results from immunoblot analyses showed that the most significant downregulation of AR-downstream signalling was induced by active compounds from series **3** and **4** (1-aryl-5-methyl substituted pyrazoles) as well as **8** and **10** (5-aryl-1-methyl-substituted pyrazoles). Oppositely, no decrease in analysed proteins was observed upon treatment with compounds from series **2**, **8i**–**8l** and **10i**–**10l** that clearly corresponded to the results from AR-transactivation assay. In addition, only three compounds from the monosubstituted pyrazoles, namely, **1a**, **1b**, and **1e** displayed moderate downregulation of the investigated proteins.

More importantly, some compounds from series **3**, **4** and **10** significantly reduced AR protein level (**Figure 3**). This process was shown to be important for the prevention of AR re-activation by alternative signalling pathways and by androgens, and provide therapeutic option for CRPC [8,9]. Decrease in AR protein stability, connected with increased degradation was previously published for enzalutamide, bicalutamide, apalutamide and darolutamide [21], but abiraterone and galeterone contributed to enhanced AR protein degradation mainly in cells expressing mutated AR-T878A [22,23].

### 2.2. Antiproliferative properties of steroid derivatives in different PCa cell lines

Antiproliferative properties of the novel derivatives were screened on the panel of three PCa cell lines, namely LAPC-4 (expressing wild type AR), 22Rv1 (expressing AR-H875Y and splicing variant V-7) and DU145 (no AR expression). Data confirmed the targeting of AR because DU145 has stayed as the most resistant cell lines (**Table 5**). In general, effects corresponded with previous assays showing that compounds from series **3**, **4**, **8**, and **10** belong to the most active ones, while compounds from series **1** and **2** did not exert any effect on viability of PCa cells, consistently with the weak agonist activity of these derivatives. The most potent derivatives in 22Rv1 cells were compounds **3a**, **3d**, **8d–g** (< 30% viable cells in 20 μM concentration), while LAPC4 cells were the most sensitive to compounds **3d**, **4e** and **10a–h** (< 40% viable cells in 20 μM concentration). Cellular activities of compounds **8i–l**, **10i–l** displaying no AR-antagonist properties were probably related to another mechanism of action.

**Table 5.** Viability of PCa cells upon treatment with 20 μM compounds for 72 h ^a^.

| Cmpd | Viability % |  |  |
| --- | --- | --- | --- |
|  | LAPC-4 | 22Rv1 | DU145 |
| 1a | 60 | 36 | 97 |
| 1b | 70 | 97 | 96 |
| 1c | 43 | 75 | 97 |
| 1d | 44 | 57 | 55 |
| 1e | 49 | 80 | 91 |
| 1f | 58 | 62 | 100 |
| 1g | 65 | 65 | 88 |
| 1h | 40 | 97 | 99 |
| 2a | 100 | $\geq 100$ | $\geq 100$ |
| 2b | $\geq 100$ | $\geq 100$ | 100 |
| 2c | 99 | $\geq 100$ | $\geq 100$ |
| 2d | 95 | $\geq 100$ | $\geq 100$ |
| 2e | 82 | $\geq 100$ | $\geq 100$ |
| 2f | 79 | 83 | $\geq 100$ |
| 2g | 93 | 94 | 97 |
| 2h | 96 | $\geq 100$ | 99 |
| 3a | 65 | 24 | 97 |
| 3b | 88 | 37 | 93 |
| 3c | 82 | 54 | 29 |
| 3d | 38 | 26 | $\geq 100$ |
| 3e | 47 | 52 | 107 |
| 3f | 52 | 50 | 89 |
| 3g | 73 | 43 | 50 |
| 3h | 74 | 58 | 85 |
| 4a | 87 | 90 | 100 |
| 4b | 87 | 97 | $\geq 100$ |
| 4c | 57 | 77 | $\geq 100$ |
| 4d | 42 | 75 | $\geq 100$ |
| 4e | 38 | 75 | $\geq 100$ |
| 4g | 78 | 87 | $\geq 100$ |
| 4h | 48 | 57 | $\geq 100$ |
|  | LAPC-4 | 22Rv1 | DU145 |
| 8a | 51 | 48 | 100 |
| 8b | 52 | 36 | 98 |
| 8c | 51 | 33 | $\geq 100$ |
| 8d | 44 | 26 | 89 |
| 8e | 41 | 29 | 90 |
| 8f | 42 | 21 | 94 |
| 8g | 50 | 18 | 100 |
| 8h | 46 | 44 | 100 |
| 8i | 72 | 47 | 12 |
| 8j | 30 | 4 | 5 |
| 8k | 63 | 37 | 57 |
| 8l | 52 | 52 | 98 |
| 10a | 36 | 68 | $\geq 100$ |
| 10b | 34 | 73 | $\geq 100$ |
| 10c | 35 | 58 | 97 |
| 10d | 29 | 57 | 90 |
| 10e | 21 | 61 | 98 |
| 10f | 20 | 56 | 96 |
| 10g | 39 | 43 | 100 |
| 10h | 31 | 68 | 100 |
| 10i | 61 | 19 | 19 |
| 10j | 83 | 93 | 98 |
| 10k | 67 | 57 | 94 |
| 10l | 53 | 103 | $\geq 100$ |
| Enzalutamide | 81 | $\geq 100$ | 96 |
| Abiraterone | $\geq 100$ | $\geq 100$ | $\geq 100$ |
| Galeterone | 76 | 95 | $\geq 100$ |
| Bavdegalutamide | 5 | 96 | 85 |

<sup>a</sup> Measured in duplicate and repeated twice.

In the consequence of relatively low antiproliferative potential of novel compounds, we evaluated their effect on the formation of colonies from LAPC-4 cells during the 10 days period. As shown in **Tables 2–4**, **Figures S4 and S5**, all compounds from series **3**, **4**, **8** and **10** were potent to block the formation of cell colonies in 5 μM concentration. Consistently with previous results, compounds from series **2** and **8i**–**l**, **10i**–**l** displayed only weak effect comparable to enzalutamide. On the other hand, many compounds from series **1** effectively inhibited colonies growth, but our abovementioned results indicated the targeting of non-AR related processes in cells.

### 2.3. Further profiling of candidate compounds

As a next step, we selected 10 hits (compounds **3a**, **3d**, **3g**, **3f**, **4c**, **4e**, **4g**, **8g**, **8h**, and **10f**) from previous assays (potent AR antagonists, AR degraders and inhibitors of formation of LAPC-4 colonies) for further profiling to compare their effect side by side. We excluded all compounds with poor solubility (e.g. **3e**) and those showing agonist or dualist mode of action in reporter assay.

Results from LNCaP cells (AR-Thr877Ala model) treated with 1nM R1881 along with our hits (**Figure 4A**) showed that compounds **3a**, **3d**, **3f** belong to the most potent derivatives, as clearly documented mainly by suppression of R1881-stimulated phosphorylation of AR at serine 81, an important marker of AR status in cells [24,25]. Interestingly, the expression of AR was only slightly decreased. The inhibitory effect of the compounds on AR-regulated signalling was further supported by the observation of the Nkx3.1 downregulation (most significantly in the presence of **3a**, **3d**, **4c** and **8g**) contrary to PSA expression.

**Figure 4.**
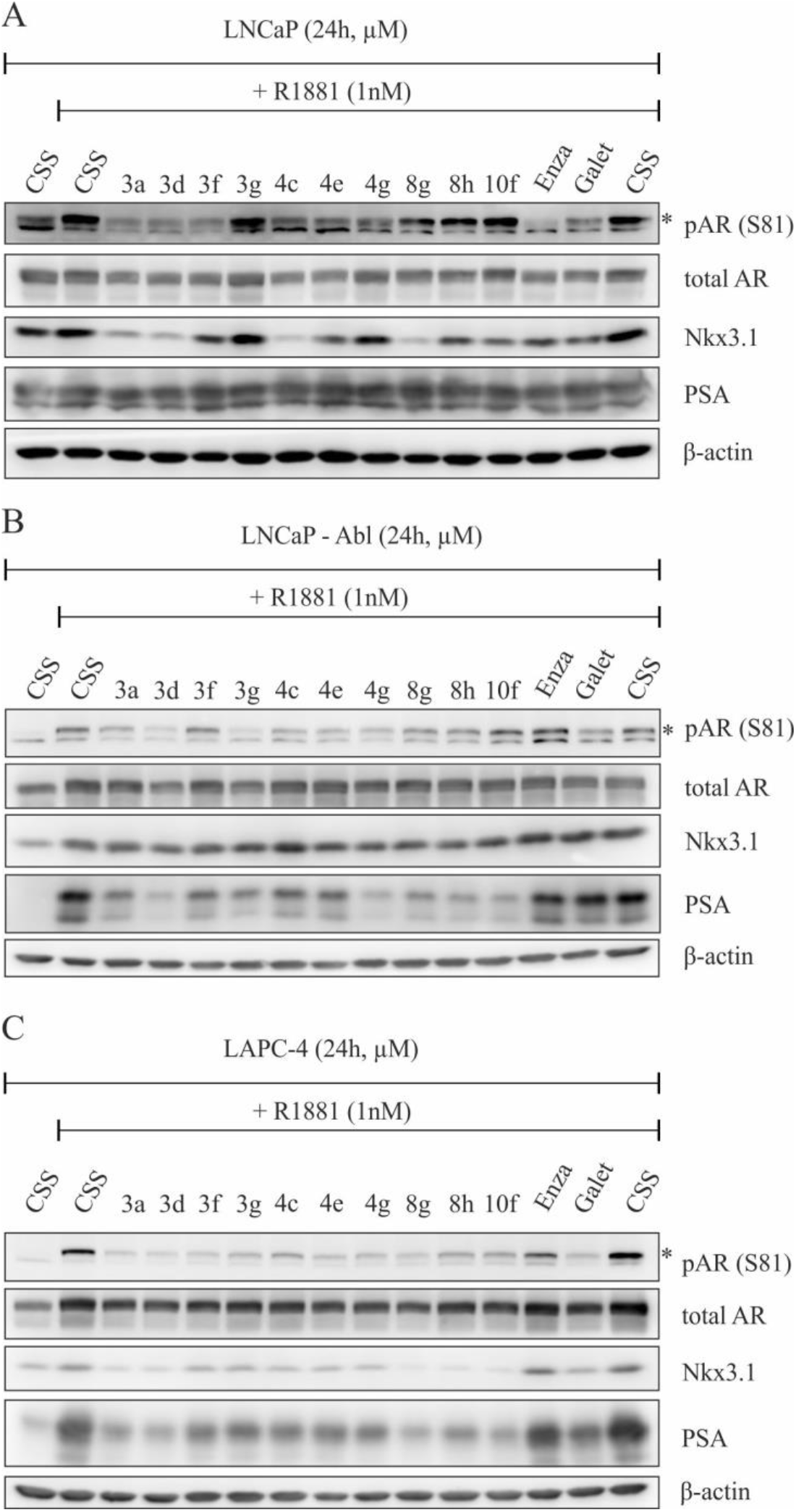
(A) LNCaP, (B) LNCaP-Abl or (C) LAPC-4 cells were cultivated in CSS containing medium, and then stimulated with 1 nM R1881 in the presence of studied steroids (10 μM) for 24 h. Lysates were then blotted for detection of appropriate proteins. Stars indicate bands corresponding with phosphorylated AR at S81. β-actin served as protein loading control.

The same experiment was performed also in LNCaP – Abl cells that were established after long-term cultivation in androgen-depleted medium resulted in AR hypersensitivity [5]. As shown in **Figure 4B**, most of the compounds are able to block AR activation as documented by monitoring its phosphorylation at S81 and PSA expression. The most profound changes were observed in the cells treated with compounds **3d** and **4g**. Furthermore, sharp decreases in both Nkx3.1 and PSA protein levels were observed upon treatment of LAPC-4 cells and **3a**, **3d**, **8g**, **8h** and **10f** belonged to the most potent compounds (**Figure 4C**). Also, based on the significant decreases in phosphorylation at S81, LAPC-4 cells (AR-wt model) were found to be the most sensitive cell line within our PCa panel.

### 2.4. Targeting the AR with compound 3d *in vitro*

We selected compound **3d** for further biological evaluation, mainly due its pronounced effects (PSA and Nkx3.1 decline, diminishing of S81 phosphorylation) in tested PCa cell lines. Preliminary data were obtained from several PCa cell lines treated with 10 μM concentration in two different conditions, without stimulation by R1881 in FBS containing media or with stimulation by R1881 in CSS containing media. We therefore decided to evaluate the changes in AR and downstream targets precisely in different concentrations and time of treatment in LAPC-4 cells. In the experiment where stimulation of AR signalling is evoked by synthetic androgen R1881, we observed dose-dependent suppression of AR signalling up to 10 μM concentration of **3d** that was comparable to galeterone’s effect in LAPC-4 (**Figure 5A**), as well as in 22Rv1 and LNCaP (**Figure S6**). Long-term treatment of LAPC-4 showed that the AR expression significantly decreased in tested concentrations after 48 and 72 h (**Figure 5B**), as also observed for PROTAC-based AR antagonist ARV110 (bavdegalutamide). Moreover, the expression of PSA and Nkx3.1 was completely switched off upon 72-h treatment with **3d**. Importantly, the long-term treatment with **3d** caused only proliferation inhibition (still > 80 % cells alive after 24h and 48 h treatment; decrease in Myc expression) without massive induction of apoptosis as documented by monitoring of PARP-1 and procaspases protein expression including their fragmentation (**Figure S7**). We therefore confirmed that the decrease in AR protein level is not directly evoked by cell death.

**Figure 5.**
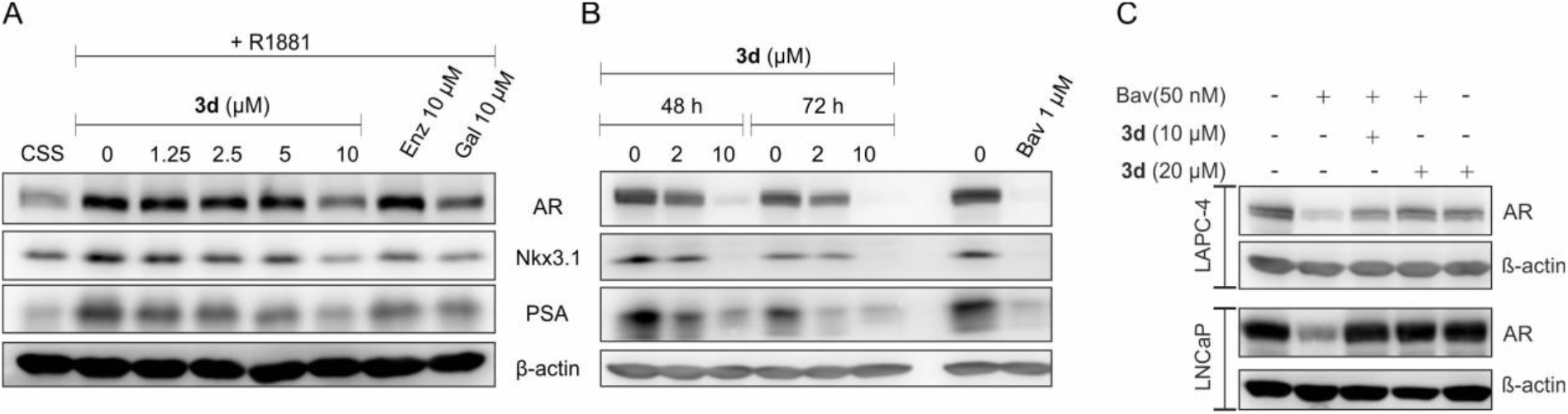
Effect of **3d** on expression of AR and downstream targets PSA and Nkx3.1 in LAPC-4 cells. (A) Cells were cultivated in charcoal-stripped serum medium for 24 h, and then stimulated with 1 nM R1881 alone or with different doses of **3d** for additional 24 h. (B) Cells were cultivated in standard media and treated with **3d** for indicated time. (C) LAPC-4 and LNCaP cells were pre-treated with **3d** for 1 h, and then bavdegalutamide was added for the next 4h. β-actin level served as protein loading control. Enz, enzalutamide; Gal, galeterone; Bav, bavdegalutamide

In rescue experiment, we verified the ability of **3d** to bind to AR cavity. LAPC-4 and LNCaP cells were pre-treated with **3d** for 1 h to saturate the AR LBD domain, and then bavdegalutamide (an effective AR degrader) was added for 4 h. As showed in **Figure 5C**, the degradation of AR in the presence of bavdegalutamide was blocked by different doses of **3d** which confirmed targeting the AR.

The binding of the candidate compound **3d** into AR-LBD was evaluated using flexible molecular docking into the LBD of AR (PDB:2PIV) crystallised with natural agonist DHT. Key interaction residues in both extremities of the LBD cavity such us Gln711, Arg752, Thr877, and Asn705 were set as flexible. The flexible docking of compound **3d** revealed extensive binding in AR-LBD with similar positions of interacting residues as in published antagonist model [26]. First two poses of **3d** were characterised by high binding energy (ΔG_Vina_ = −11.8 kcal/mol and −11.6 kcal/mol, respectively), and similar orientation with nearly identical interactions in the 1’-(4”-fluorophenyl)-5’-methylpyrazolo part of **3d**, where Arg752 and Gly583 create a halogen bond with fluorine on the phenyl ring, which is probably stabilized by hydrophobic interactions with side chains of Gln711, Met749, Val684 and Ala748 (**Figure 6A, B**). The methyl group (on the pyrazole ring of **3d**) forms a hydrophobic interaction with Leu707, while the fused pyrazole ring interacts with Phe764 and might also form a hydrogen bond with Met745.

**Figure 6.**
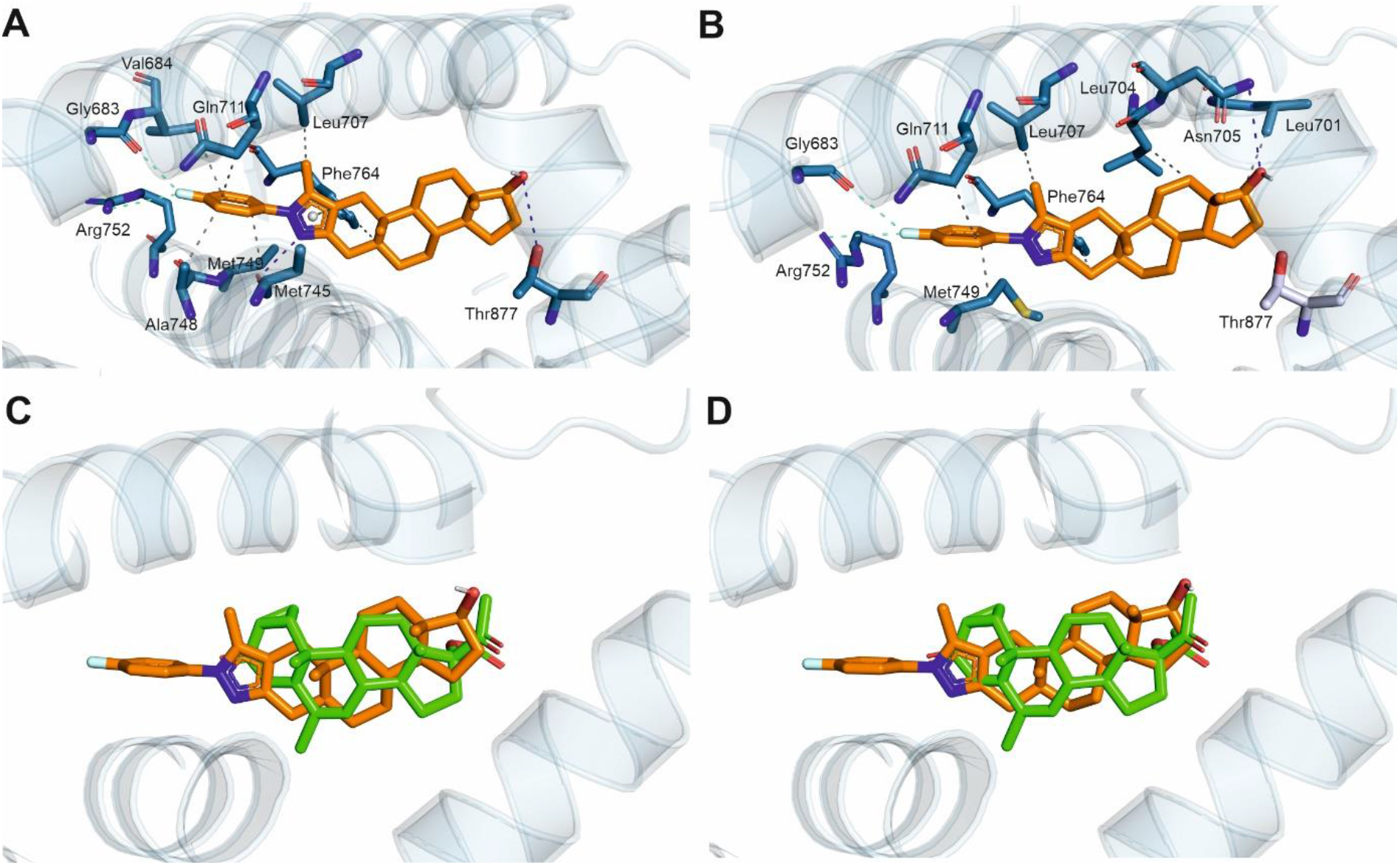
Binding poses of **3d** in the ligand binding domain of AR (PDB:2PIV) performed by flexible docking. The first pose (A) shows interactions including Thr877, while the second pose (B) shows binding independent of Thr877. Respective alignments with cyproterone is shown as well (C, D). Sticks represent interacting amino acid residues. Nitrogen atoms are shown in blue, oxygen atoms in red, fluorine atom in cyan. Hydrogen bonds are shown with blue dash lines, halogen bonds with cyan dash lines and hydrophobic interactions are shown with grey dash lines.

The key difference was observed in the interaction of the 17β-OH on the D-ring of **3d**. In the first pose the hydroxyl group forms a conserved hydrogen bond with Thr877 (known from the binding of DHT) (**Figure 6A**), while in the second pose the steroid core is positioned in a slightly different angle towards Asn705, where it forms a hydrogen bond with the 17β-OH group. Further stabilization of the steroidal ring of **3d** is mediated by hydrophobic interactions with neighbouring leucines 701 and 704. Most importantly, the second pose shows a binding pose independent of Thr877 which is mutated in LNCaP cell line by point mutation T877A. Overall, lead compound should bind to the same region with an orientation similar to the known antagonist cyproterone (PDB:2OZ7) [26,27] (**Figure 6C, D**). The interaction of our lead compound independent of Thr877 should explain its potency even in PCa bearing mutated AR-LBD.

Next, the binding of **3d** in human recombinant AR-LBD was examined by the micro-scale thermophoresis using the Protein Labeling Kit RED-NHS (Nanotemper) The binding of the **3d** (tested in 25μM – 0.25 μM range) induced significant changes in the protein mobility marked by the change of the fluoresence signal (**Figure S8)** and confirmed thus binding of **3d** into the AR-LBD.

Further, we evaluated the antiproliferative effect of **3d** in different cell lines in dose dependent manner using colony formation assay and resazurine-based viability assay. We confirmed the increased sensitivity to AR-positive PCa cell lines with GI_50_ values in low micromole ranges and to wt-AR expressing LAPC-4 cells, being the most sensitive (GI_50_ = 7.9 ± 1.6 μM) (see **Figure S9**). Moreover, cell cycle analysis after 48 h of treatment showed a reduced number of cells in S-phase of the cell cycle, which confirmed to us the negative impact of **3d** on proliferation of AR-positive LAPC-4 and 22Rv1 cells, while having no effect on AR-negative DU145 cells (**Figure S10**).

Finally, we examined the effect of different doses of **3d** on R1881-stimulated transcriptional activity in reporter cell line, and it was found that IC_50_ value of **3d** (1.18 μM) shows higher potency compared to galeterone (7.59 μM) and enzalutamide (3.32 μM) assayed as controls (**Figure 7A**). We further evaluated the functional consequences of the diminishing of the AR transactivation, and performed qPCR analysis of mRNA expression of AR and PSA, respectively. As showed in **Figure 7B**, **3d** inhibited PSA mRNA expression in 22Rv1 cells more potently than enzalutamide and galeterone. We also observed comparable results in LAPC-4 cells under similar experimental conditions, where the expression level of AR transcript decreased moderately, as well (**Figure S11**).

**Figure 7.**
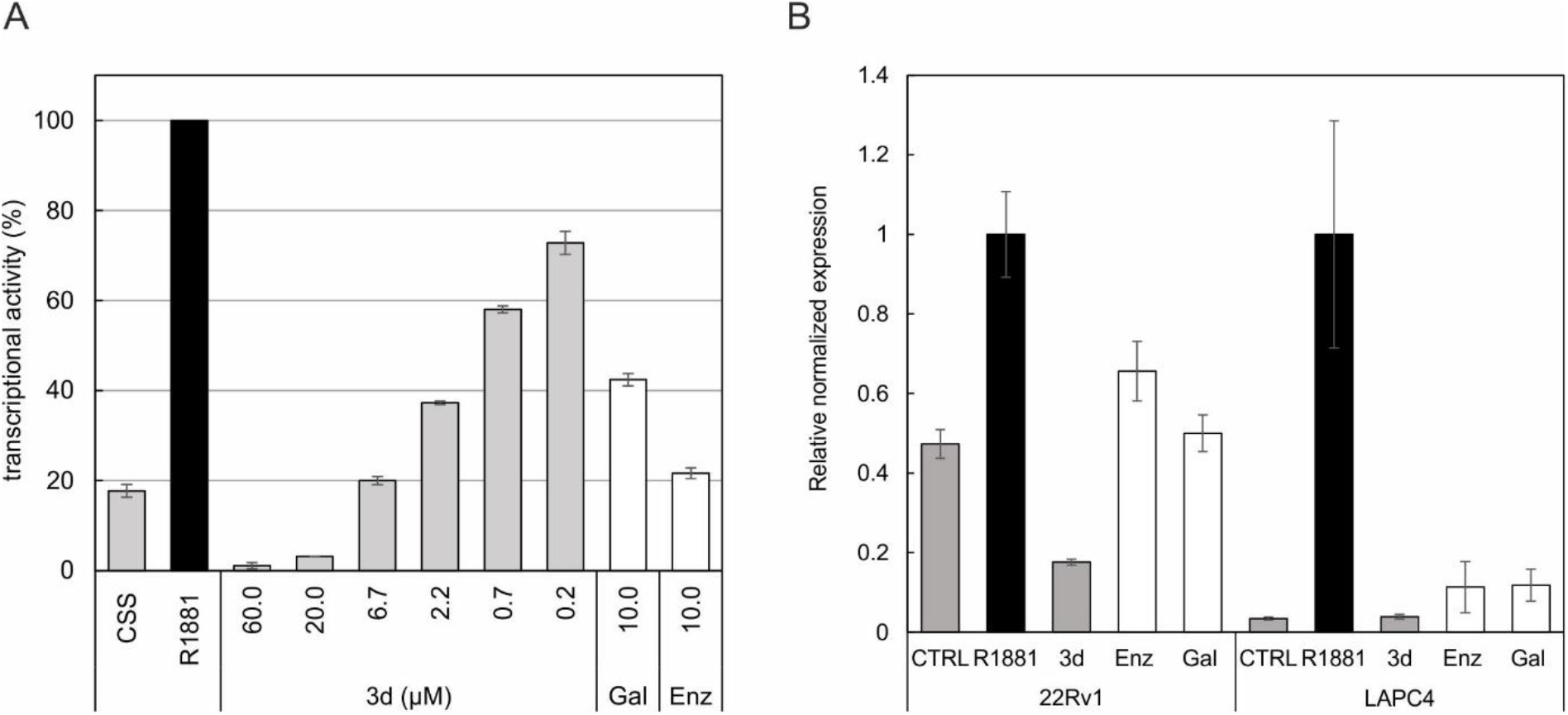
(A) The effect of **3d** on the AR-mediated transcription in the 22Rv1-ARE14 reporter cell line. Cells were stimulated with 1 nM R1881 (R) alone (black column) or with different doses of **3d** (grey columns) for 24 h in charcoal-stripped serum medium (CSS), and then, the luciferase activity in the cell lysate was measured (B) The effect of compound **3d** and standard AR antagonists on relative normalized mRNA expression of AR-downstream gene KLK3 (PSA). Cells were cultivated in CSS medium overnight, then treated with compounds in 10 μM concentration in presence of 1 nM R1881 for 24 h. Enz, enzalutamide; Gal, galeterone.

### 2.5. Targeting the AR with compound 3d *ex vivo*

Selected compounds **3d** and **10f** were preliminary tested in a short-term *ex vivo* culture of patient-derived samples. An experienced pathologist provided non-diagnostic tissues (approximately 0.5 cubic centimeter) from five patients undergoing robotic prostatectomy. The tissues were cut with vibratome, and slices were treated for three days with our candidates (10 μM) along with enzalutamide and bavdegalutamide (1 μM). Immunohistochemistry results of proliferation marker Ki67 and AR are provided in **Figure 8** and **Figure S12**. Samples from one patient were excluded due to missing cancer cells in some tissue slices. Compound **3d** caused a mild decrease of the level of Ki67 (marker of proliferation) and heterogenous effect on the AR level, which was slightly decreased in 3 from 4 patients. Tissue slices obtained from patient 1 display the most homogenous morphology and downregulation of AR is clearly visible after treatment with compound **3d**, enzalutamide and bavdegalutamide (see **Figure S12**). Tissue slices from other patients displayed heterogeneous tissue morphology which hampers direct comparison of the treatments. However, enzalutamide also demonstrated a limited effect only, while bavdegalutamide was shown to be the most potent. Our candidates will be further tested on other patient-derived tissues, as well as organoids which may clarify their therapeutic potential.

**Figure 8.**
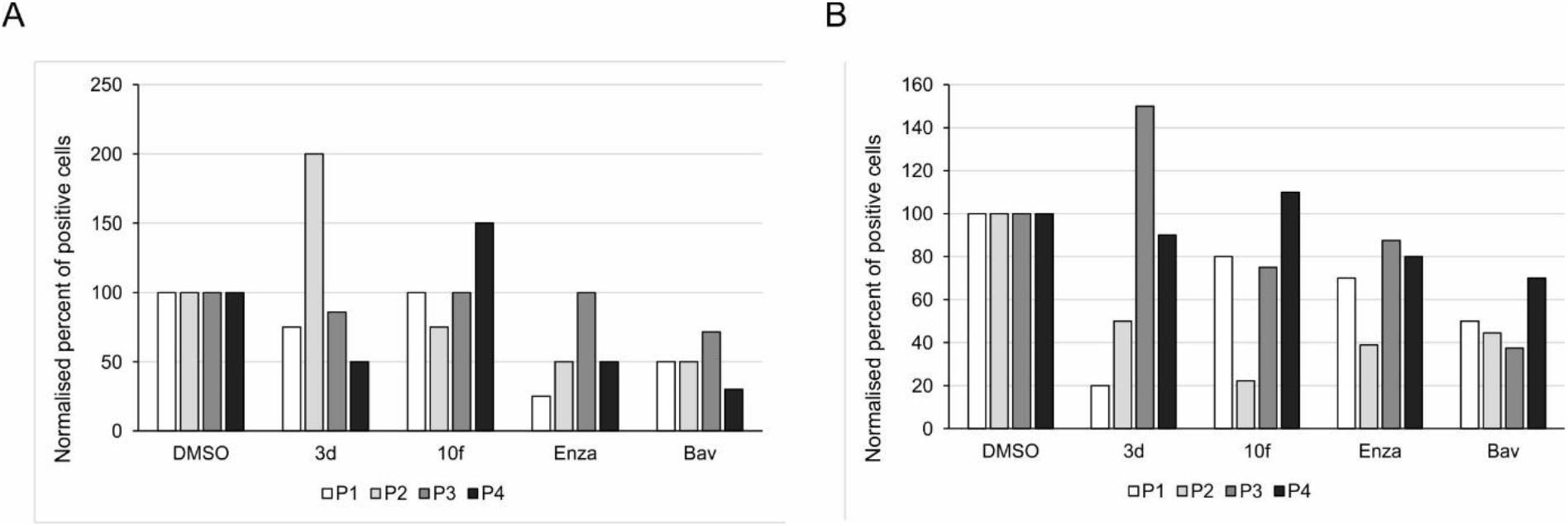
AR (A) and Ki67 (B) levels from the highest-intensity positive cells assessed by immunostaining of tissue from 4 patients in the explants after 3 days of *ex vivo* culture in the presence of indicated compounds. Candidate compounds **3d** and **10f** were applied in 10 μM concentrations, while standards enzalutamide (Enz) and bavdegalutamide (Bav) in 1 μM concentrations. All immunohistochemistry images were evaluated using histoscore method and normalized to DMSO-treated samples.

## 3. Conclusion

In summary, our existing compound library of DHT-derived A-ring-fused pyrazoles has been extended to novel derivatives synthesized from α,β-enones with monosubstituted hydrazines. The heterocyclization and subsequent oxidation led to the regioselective formation of the thermodynamically favoured 1,5-disubstituted heterocyclic compounds under conventional heating. A total of 55 differently functionalized derivatives were subjected to the evaluation of their impact on the AR signalling in several PCa models. Lead compound **3d** displayed significant potency to disrupt AR signalling in both castration-sensitive and resistant PCa cell lines, with selective antiproliferative potency towards AR-positive cell lines. Treatment with 10 μM of compound **3d** induced massive reduction in AR protein level, slight decrease in AR transcript and total blockage in AR downstream targets (PSA, Nkx3.1), which was associated with the decrease in S-phase cells and proliferation blockage. Moreover, *ex vivo* activity was shown on PCa patient’s biopsies with overall encouraging potential as a PCa anticancer agent.

## 4. Experimental

### 4.1. General

Chemicals, reagents and solvents were purchased from commercial suppliers (Sigma-Aldrich, TCI and Alfa Aesar) and used without further purification. Melting points (Mp) were determined on an SRS Optimelt digital apparatus and are uncorrected. For MW-assisted syntheses, a CEM Discover SP laboratory MW reactor was used with a max. power of 200 W (running a dynamic control program). Elementary analysis data were obtained with a PerkinElmer CHN analyzer model 2400. The transformations were monitored by TLC using 0.25 mm thick Kieselgel-G plates (Si 254 F, Merck). The compound spots were detected by spraying with 5% phosphomolybdic acid in 50% aqueous phosphoric acid. Flash chromatographic purifications were carried out on silica gel 60, 40–63 μm (Merck). NMR spectra were recorded with a Bruker DRX 500 instrument at room temperature in CDCl_3_ and DMSO-*d*_6_ using residual solvent signal as an internal reference. Chemical shifts are reported in ppm (*δ* scale) and coupling constants (*J*) are given in Hz. Multiplicities of the ^1^H signals are indicated as a singlet (s), a broad singlet (bs), a doublet (d), a double doublet (dd), a triplet (t), or a multiplet (m). ^13^C NMR spectra are ^1^H-decoupled and the J-MOD pulse sequence was used for multiplicity editing. In this spin-echo type experiment, the signal intensity is modulated by the different coupling constants *J* of carbons depending on the number of attached protons.

Both protonated and unprotonated carbons can be detected (CH_3_ and CH carbons appear as positive signals, while CH_2_ and C carbons as negative signals). The purified derivatives were dissolved in high purity acetonitrile and introduced with an Agilent 1290 Infinity II liquid chromatography pump to an Agilent 6470 tandem mass spectrometer equipped an electrospray ionization chamber. Flow rate was 0.5 mL·min^-1^, and contained 0.1% formic acid or 0.1% ammonium hydroxide to help facilitate ionization. The instrument operated in MS1 scan mode with 135 V fragmentor voltage, and the spectra were recorded from 200 to 600 m/z, which were corrected with the background.

### 4.2. Chemistry

#### 4.2.1. Synthesis of the A-ring-modified a,β-enones

Compounds **5a–e, 5g** and **5h** were synthesized as described previously [18].

##### 4.2.1.1. 17β-Hydroxy-2-(4-bromo)benzylidene-5α-androstan-3-one (**5f**)

According to the general method described previously [18], 4-bromobenzaldehyde (666 mg) was used for the reaction. Reaction time: 3 h. White solid. Yield: 1.18 g (86%); Mp 205–207 °C; ^1^H NMR (CDCl_3_, 500 MHz): *δ*_H_ 0.73 (s, 3H, 18-H_3_), 0.80 (s, 3H, 19-H_3_), 0.85-1.02 (overlapping m, 3H), 1.11 (m, 1H), 1.22-1.65 (overlapping m, 9H), 1.78 (m, 3H), 2.07 (m, 1H), 2.14 (d, 1H, *J* = 15.7 Hz, one of 1-H_2_), 2.23 (dd, 1H, *J* = 18.6 Hz, 13.3 Hz, one of 4-H_2_), 2.45 (dd, 1H, *J* = 18.6 Hz, *J* = 5.2 Hz, the other of 4-H_2_), 3.03 (d, 1H, *J* = 15.7 Hz, the other of 1-H_2_), 3.65 (t, 1H, *J* = 8.6 Hz, 17-H), 7.23 (d, 2H, *J* = 8.4 Hz, 3’-H and 5’-H), 7.45 (s, 1H, 2a-H), 7.52 (d, 2H, *J* = 8.4 Hz, 2’-H and 6’-H); ^13^C NMR (CDCl_3_, 125 MHz): *δC* 11.2 (C-18), 12.0 (C-19), 21.2 (CH_2_), 23.5 (CH_2_), 28.7 (CH_2_), 30.7 (CH_2_), 31.2 (CH_2_), 35.6 (CH), 36.3 (C-10), 36.8 (CH_2_), 42.0 (CH_2_), 42.7 (CH), 42.9 (CH_2_), 43.0 (C-13), 51.1 (CH), 53.8 (CH), 82.0 (C-17), 122.9 (C-4’), 131.8 (2C, C-2’ and C-6’), 131.8 (2C, C-3’ and C-5’), 134.7 (C-1’), 136.0 (C-2a), 136.1 (C-2), 201.4 (C-3); ESI-MS 457 [M+H]^+^; Anal. Calcd. for C_26_H_33_BrO_2_ C 68.27; H 7.27. Found C 68.29; H 7.28.

##### 4.2.1.2. 17β-Hydroxy-2-(4-hydroxy)benzylidene-5α-androstan-3-one (**5i**)

According to the general method described previously [18], 4-(methoxymethoxy)benzaldehyde (598 mg) was used for the reaction. The reaction mixture was heated under reflux temperature for 16 h. MOM-protected-**5i** was obtained as a yellow crystal. Yield: 682 mg (52%). This reaction was repeated to obtain a sufficient amount of protected derivative, then 877 mg (2 mmol) was dissolved in hot MeOH followed by the addition of 6 M HCl solution (0.2 mL) dropwise. The reaction mixture was stirred for 30 min, and then cooled to room temperature. The mixture was poured into water and extracted with EtOAc (3 *×* 10 mL). The combined organic layer was washed with water (2 × 20 mL) and brine (20 mL), dried over anhydrous Na_2_SO_4_ and the solvent was evaporated *in vacuo*. The crude product was purified by column chromatography (silica gel, EtOAc/CH_2_Cl_2_=10:90 to EtOAc/CH_2_Cl_2_=50:50 using gradient elution). Yield: 554 mg (70%); Mp > 250 °C (decomp.); ^1^H NMR (DMSO-*d*_6_, 500 MHz): *δ*_H_ 0.62 (s, 3H, 18-H_3_), 0.71 (s, 3H, 19-H_3_), 0.89 (overlapping m, 3H), 1.02 (m, 1H), 1.11-1.42 (overlapping m, 7H), 1.50 (m, 1H), 1.58-1.65 (overlapping m, 2H), 1.77 (m, 1H), 1.84 (m, 1H), 2.11 (dd, 1H, *J* = 18.7, 13.1 Hz, one of 4-H_2_), 2.24 (d, 1H, *J* = 15.9 Hz, one of 1-H_2_), 2.30 (dd, 1H, *J* = 18.7, 5.4 Hz, the other of 4-H_2_), 2.94 (d, 1H, *J* = 16.0 Hz, the other of 1-H_2_), 3.45 (t, 1H, *J* = 8.1 Hz, 17-H), 4.42 (bs, 1H, 17-OH), 6.83 (d, 2H, *J* = 8.6 Hz, 3’-H and 5’-H), 7.34 (d, 2H, *J* = 8.6 Hz, 2’-H and 6’-H), 7.35 (s, 1H, 2a-H), 9.92 (s, 1H, 4’-OH); ^13^C NMR (DMSO-*d*_6_, 125 MHz): *δ*_C_ 11.1 (C-18), 11.7 (C-19), 20.5 (CH_2_), 23.1 (CH_2_), 28.0 (CH_2_), 29.9 (CH_2_), 30.7 (CH_2_), 35.0 (CH), 35.3 (C-10), 36.6 (CH_2_), 41.3 (CH), 41.5 (CH_2_), 42.1 (CH_2_), 42.4 (C-13), 50.4 (CH), 53.0 (CH), 80.0 (C-17), 115.5 (2C, C-3’ and C-5’), 126.0 (C-1’), 132.1 (C-2), 132.4 (2C, C-2’ and C-6’), 136.5 (C-2a), 158.3 (C-4’) 199.3 (C-3); ESI-MS 395 [M+H]^+^; Anal. Calcd. for C_26_H_34_O_3_ C 79.15; H 8.69. Found C 79.08; H 8.68.

#### 4.2.2. General procedures for the synthesis of DHT-derived A-ring-fused pyrazoles

##### Method A

1 mmol arylidene (**5a**) and methylhydrazine (105 μL, 2 equiv.) were dissolved in EtOH (4 mL), and the mixture was irradiated in a closed vessel at 100 °C for 3 min. After completion of the reaction, the mixture was poured into ice water (10 mL), NH4Cl was added and the white precipitate formed was filtered *in vacuo*, washed with water and dried. The solid thus obtained containing both **6a** and **7a** was oxidized: the crude product was dissolved in 5 mL 1,4-dioxane, and DDQ (250 mg, 1.1 equiv.) was added. The reaction mixture was irradiated in a closed vessel at 120 °C for 5 min. After completion of the reaction, the mixture was poured into ice water, NH4Cl was added, and the resulting precipitate was filtered and dried. The crude product containing **8a** and **9a** was purified by column chromatography with EtOAc/CH_2_Cl_2_ = 10:90.

##### Method B

Arylidene (**5a**–**f**, **5i**), heteroarylidene (**5g**, **5h**) or ethylidene derivative (**5j**) [17] (1.0 mmol) and methylhydrazine sulfate (288 mg, 2 equiv.) were dissolved in absolute EtOH (15 mL) and the mixture was stirred at 78 °C for 24 h, during which in addition to the complete conversion of the starting material, the spontaneous oxidation of the heteroring also occurred in most of the cases. During work-up, the mixture was cooled to room temperature, poured into water and extracted with CH_2_Cl_2_ (3 *×* 10 mL). The combined organic layer was washed with water (2 × 20 mL) and brine (20 mL), dried over anhydrous Na_2_SO_4_ and the solvent was evaporated *in vacuo*. The crude product was purified by column chromatography except in the reaction of **5d–5f** where oxidation of the heteroring with DDQ was needed. Thus, in this latter case, the residue was dissolved in 1,4-dioxane (10 mL) and DDQ (250 mg, 1.1 equiv.) was added. The mixture was irradiated in a closed vessel at 120 °C for 5 min, then poured into ice-cold water. NH4Cl was added and the resulting precipitate was filtered and dried. The crude product was purified by column chromatography.

##### 4.2.2.1. 17β-Hydroxy-1’-methyl-5’-phenylpyrazolo[3’,4’:3,2]-5α-androstane (**8a**)

According to Section *4.2.2., Method A* or *B,* 379 mg of **5a** was used. Eluent: EtOAc/CH_2_Cl_2_ = 10:90. Off white solid. Yield: 232 mg (57%, *Method A),* or 323 mg (80%, *Method B);* Mp 157–160 °C; ^1^H NMR (CDCl_3_, 500 MHz): *δ*H 0.74 (s, 3H, 18-H_3_), 0.76 (s, 3H, 19-H_3_), 0.81-1.00 (overlapping m, 3H, 9α-H, 7α-H and 14α-H), 1.05 (m, 1H, 12α-H), 1.23-1.47 (overlapping m, 5H, 15β-H, 11β-H, 6β-H, 8β-H and 16β-H), 1.55-1.65 (overlapping m, 4H, 11α-H, 5α-H, 15α-H and 6α-H), 1.73 (m, 1H, 7β-H), 1.79 (m, 1H, 12β-H), 2.06 (m, 1H, 16α-H), 2.13 (d, 1H, *J* = 15.3 Hz, 1α-H), 2.33 (dd, 1H, *J* = 16.4, 12.2 Hz, 4β-H), 2.47 (d, 1H, *J* = 15.3 Hz, 1β-H), 2.67 (dd, 1H, *J* = 16.4, 5.1 Hz, 4α-H), 3.63 (t, 1H, *J* = 8.5 Hz, 17α-H), 3.77 (s, 3H, N-CH_3_), 7.31 (d-like m, 2H, 3”-H and 5”-H), 7.38 (t-like m, 1H, 4”-H), 7.46 (t-like m, 2H, 2”-H and 6”-H); ^13^C NMR (CDCl_3_, 125 MHz): *δ*C 11.2 (C-18), 11.8 (C-19), 21.0 (C-11), 23.6 (C-15), 27.9 (C-4), 29.5 (C-6), 30.8 (C-16), 31.6 (C-7), 35.7 (C-1), 36.1 (C-8), 36.6 (C-10), 37.0 (C-12), 37.2 (N-CH_3_), 42.8 (C-5), 43.0 (C-13), 51.2 (C-14), 54.3 (C-9), 82.1 (C-17), 114.3 (C-2), 128.1 (C-4”), 128.8 (2C) and 129.3 (2C): C-2”, C-6”, C-3”, C-5”, 131.1 (C-1”), 140.2 (C-5’) 147.3 (C-3); ESI-MS 405 [M+H]^+^; Anal. Calcd. for C_27_H_36_N_2_O C 80.15; H 8.97. Found C 80.21; H 8.98.

##### 4.2.2.2. 17β-Hydroxy-1’-methyl-3’-phenylpyrazolo[4’,3’:2,3]-5α-androstane (**9a**)

According to Section *4.2.2., Method A,* 379 mg of **5a** was used. Eluent: EtOAc/CH_2_Cl_2_ = 10:90.. Off white solid. Yield: 114 mg (28%); Mp 232–236 °C; ^1^H NMR (CDCl_3_, 500 MHz): *δ*H 0.73 (s, 3H, 18-H_3_), 0.76 (s, 3H, 19-H_3_), 0.86-1.03 (overlapping m, 3H, 9α-H, 7α-H and 14α-H), 1.13 (m, 1H, 12α-H), 1.28 (m, 1H, 15β-H), 1.35-1.49 (overlapping m, 4H, 6β-H, 8β-H, 11β-H and 16β-H), 1.60-1.77 (overlapping m, 5H, 15α-H, 5α-H, 6α-H, 11α-H, 7β-H), 1.86 (m, 1H, 12β-H), 2.07 (m, 1H, 16α-H), 2.21 (dd, 1H, *J* = 16.2, 12.4 Hz, 4β-H), 2.32 (d, 1H, *J* = 15.1 Hz, 1α-H), 2.53 (dd, 1H, *J* = 16.2, 4.9 Hz, 4α-H), 2.76 (d, 1H, *J* = 15.1 Hz, 1β-H), 3.66 (t, 1H, *J* = 8.5 Hz, 17α-H), 3.79 (s, 3H, N-CH_3_), 7.29 (t-like m, 1H, 4”-H), 7.40 (t-like m, 2H, 3”-H and 5”-H), 7.70 (d, 2H, *J* = 7.3 Hz, 2”-H and 6”-H); ^13^C NMR (CDCl_3_, 125 MHz): *δ*_C_ 11.2 (C-18), 11.7 (C-19), 21.0 (C-11), 23.6 (C-15), 26.3 (C-4), 29.2 (C-6), 30.7 (C-16), 31.4 (C-7), 35.7 (N-CH_3_), 35.9 (C-8), 36.6 (C-10), 36.8 (C-1), 36.9 (C-12), 41.8 (C-5), 43.0 (C-13), 51.1 (C-14), 54.1 (C-9), 82.0 (C-17), 113.0 (C-2), 126.8 (2C, C-2” and C-6”), 127.1 (C-4”), 128.6 (2C, C-3” and C-5”), 134.4 (C-1”), 138.8 (C-5’) 147.6 (C-3’); ESI-MS 405 [M+H]^+^; Anal. Calcd. for C_27_H_36_N_2_O C 80.15; H 8.97. Found C 80.08; H 8.95.

##### 4.2.2.3. 17β-Hydroxy-1’-methyl-5’-(4”-tolyl)-pyrazolo[3’,4’:3,2]-5α-androstane (**8b**)

According to Section *4.2.2., Method B,* 393 mg of **5b** was used. Eluent: EtOAc/CH_2_Cl_2_ = 5:95.. Off white solid. Yield: 340 mg (81%); Mp 124–126 °C; ^1^H NMR (CDCl_3_, 500 MHz): *δ*_H_ 0.74 (s, 3H, 18-H_3_), 0.75 (s, 3H, 19-H_3_), 0.80-0.99 (overlapping m, 3H), 1.06 (m, 1H), 1.23-1.47 (overlapping m, 5H), 1.54-1.64 (overlapping m, 4H), 1.73 (m, 1H), 1.78 (m, 1H), 2.06 (m, 1H), 2.11 (d, 1H, *J* = 15.2 Hz, 1α-H), 2.31 (dd, 1H, *J* = 16.5, 12.1 Hz, 4β-H), 2.41 (s, 4”-CHs), 2.46 (d, 1H, *J* = 15.2 Hz, 1β-H), 2.66 (dd, 1H, *J* = 16.5, 5.0 Hz, 4α-H), 3.63 (t, 1H, *J* = 8.5 Hz, 17α-H), 3.77 (s, 3H, N-CH_3_), 7.21 (d, 2H, *J* = 8.0 Hz), 7.26 (d, 2H, *J* = 8.0 Hz); ^13^C NMR (CDCl_3_, 125 MHz): *δ*_C_ 11.2 (C-18), 11.7 (C-19), 21.0 (C-11), 21.4 (4”-CH_3_), 23.6 (C-15), 27.8 (C-4), 29.4 (C-6), 30.7 (C-16), 31.5 (C-7), 35.6 (C-1), 36.0 (C-8), 36.5 (C-10), 36.9 (C-12), 37.1 (N-CH_3_), 42.7 (C-5), 43.0 (C-13), 51.2 (C-14), 54.3 (C-9), 82.1 (C-17), 114.1 (C-2), 127.9 (C-4”), 129.2 (2C, C-2” and C-6”), 129.5 (2C, C-3” and C-5”), 138.1 (C-1”), 140.2 (C-5’), 147.2 (C-3); ESI-MS 419 [M+H]^+^; Anal. Calcd. for C_28_H_38_N_2_O C 80.34; H 9.15. Found C 80.11; H 9.16.

##### 4.2.2.4. 17β-Hydroxy-5’-(4”-methoxyphenyl)-1’-methylpyrazolo[3’,4’:3,2]-5α-androstane (**8c**)

According to Section *4.2.2., Method B,* 409 mg of **5c** was used. Eluent: EtOAc/CH_2_Cl_2_ = 5:95. White solid. Yield: 343 mg (79%); Mp 125–128 °C; ^1^H NMR (CDCl_3_, 500 MHz): *δ*_H_ 0.74 (s, 3H, 18-H_3_), 0.75 (s, 3H, 19-H_3_), 0.80-0.99 (overlapping m, 3H), 1.06 (m, 1H), 1.23-1.47 (overlapping m, 5H), 1.55-1.65 (overlapping m, 4H), 1.73 (m, 1H), 1.79 (m, 1H), 2.05 (m, 1H), 2.10 (d, 1H, *J* = 15.2 Hz, 1α-H), 2.31 (dd, 1H, *J* = 16.4, 12.1 Hz, 4β-H), 2.45 (d, 1H, *J* = 15.2 Hz, 1β-H), 2.65 (dd, 1H, *J* = 16.4, 5.0 Hz, 4α-H), 3.63 (t, 1H, *J* = 8.5 Hz, 17α-H), 3.76 (s, 3H, N-CH_3_), 3.86 (s, 3H, 4”-OCH_3_), 6.99 (d, 2H, *J* = 8.7 Hz, 3”-H and 5”-H), 7.24 (d, 2H, *J* = 8.7 Hz, 2”-H and 6”-H); ^13^C NMR (CDCl_3_, 125 MHz): *δ*C 11.2 (C-18), 11.7 (C-19), 21.0 (C-11), 23.6 (C-15), 27.8 (C-4), 29.5 (C-6), 30.7 (C-16), 31.5 (C-7), 35.6 (C-1), 36.0 (C-8), 36.5 (C-10), 36.9 (C-12), 37.1 (N-CH_3_), 42.7 (C-5), 43.0 (C-13), 51.2 (C-14), 54.3 (C-9), 55.5 (4”-OCH_3_), 82.1 (C-17), 114.0 (C-2), 114.3 (2C, C-3” and C-5”), 123.2 (C-1”), 130.5 (2C, C-2” and C-6”), 140.0 (C-5’), 147.2 (C-3), 159.5 (C-4”); ESI-MS 435 [M+H]^+^; Anal. Calcd. for C_28_H_38_N_2_O_2_ C 77.38; H 8.81. Found C 77.36; H 8.83.

##### 4.2.2.5. 17β-Hydroxy-5’-(4”-fluorophenyl)-Γ-methylpyrazolo[3’,4’:3,2]-5α-androstane (**8d**)

According to Section *4.2.2.,Method B,* 397 mg of **5d** was used. Eluent: EtOAc/CH_2_Cl_2_ = 10:90. White solid. Yield: 278 mg (66%); Mp 185–187 °C; ^1^H NMR (CDCl_3_, 500 MHz): *δ*_H_ 0.74 (s, 3H, 18-H_3_), 0.75 (s, 3H, 19-H_3_), 0.80-0.99 (overlapping m, 3H), 1.06 (m, 1H), 1.23-1.47 (overlapping m, 5H), 1.54-1.64 (overlapping m, 4H), 1.72 (m, 1H), 1.80 (m, 1H), 2.05 (m, 1H), 2.10 (d, 1H, *J* = 15.2 Hz, 1α-H), 2.30 (dd, 1H, *J* = 16.5, 11.9 Hz, 4β-H), 2.41 (d, 1H, *J* = 15.2 Hz, 1β-H), 2.65 (dd, 1H, *J* = 16.5, 5.1 Hz, 4α-H), 3.63 (t, 1H, *J* = 7.6 Hz, 17α-H), 3.75 (s, 3H, N-CH_3_), 7.15 (t-like m, 2H), 7.28 (overlapping m, 2H); ^13^C NMR (CDCl_3_, 125 MHz): *δ*C 11.2 (C-18), 11.7 (C-19), 21.0 (C-11), 23.6 (C-15), 27.8 (C-4), 29.4 (C-6), 30.7 (C-16), 31.5 (C-7), 35.5 (C-1), 36.0 (C-8), 36.5 (C-10), 36.9 (C-12), 37.1 (N-CH_3_), 42.7 (C-5), 43.0 (C-13), 51.1 (C-14), 54.2 (C-9), 82.1 (C-17), 114.4 (C-2), 115.9 (d, 2C, *J* = 21.7 Hz, C-3” and C-5”), 127.0 (d, *J* = 3.5 Hz, C-1”), 131.1 (d, 2C, *J* = 8.2 Hz, C-2” and C-6”), 139.2 (C-5’), 147.3 (C-3), 161.6 (d, *J* = 248.4 Hz, C-4”); ESI-MS 423 [M+H]^+^; Anal. Calcd. for C_27_H_35_FN_2_O C 76.74; H 8.35. Found C 76.86; H 8.32.

##### 4.2.2.6. 17β-Hydroxy-5’-(4”-chlorophenyl)-1’-methylpyrazolo[3’,4’:3,2]-5α-androstane (**8e**)

According to Section *4.2.2.,Method B,* 413 mg of **5e** was used. Eluent: EtOAc/CH_2_Cl_2_ = 10:90. White solid. Yield: 302 mg (69%); Mp 123–125 °C; ^1^H NMR (CDCl_3_, 500 MHz): *δ*_H_ 0.74 (s, 3H, 18-H_3_), 0.75 (s, 3H, 19-H_3_), 0.80-1.00 (overlapping m, 3H), 1.07 (m, 1H), 1.23-1.47 (overlapping m, 5H), 1.54-1.66 (overlapping m, 4H), 1.74 (m, 1H), 1.80 (m, 1H), 2.05 (m, 1H), 2.11 (d, 1H, *J* = 15.2 Hz, 1α-H), 2.32 (dd, 1H, *J* = 16.5, 11.9 Hz, 4β-H), 2.43 (d, 1H, *J* = 15.2 Hz, 1β-H), 2.66 (dd, 1H, *J* = 16.5, 5.1 Hz, 4α-H), 3.63 (t, 1H, *J* = 8.5 Hz, 17α-H), 3.76 (s, 3H, N-CH_3_), 7.24 (d, 2H, *J* = 8.5 Hz, 3”-H and 5”-H), 7.44 (d, 2H, *J* = 8.5 Hz, 2”-H and 6”-H); ^13^C NMR (CDCl_3_, 125 MHz): *δ*C 11.2 (C-18), 11.8 (C-19), 21.0 (C-11), 23.6 (C-15), 27.8 (C-4), 29.5 (C-6), 30.8 (C-16), 31.6 (C-7), 35.6 (C-1), 36.1 (C-8), 36.6 (C-10), 37.0 (C-12), 37.2 (N-CH_3_), 42.8 (C-5), 43.0 (C-13), 51.2 (C-14), 54.3 (C-9), 82.1 (C-17), 114.6 (C-2), 129.1 (2C, C-2” and C-6”), 129.5 (C-1”), 130.6 (2C, C-3” and C-5”), 134.3 (C-4”), 139.0 (C-5’), 147.5 (C-3); ESI-MS 439 [M+H]^+^; Anal. Calcd. for C_27_H_35_ClN_2_O C 73.87; H 8.04. Found C 73.91; H 8.03.

##### 4.2.2.7. 17β-Hydroxy-5’-(4”-bromophenyl)-Γ-methylpyrazolo[3’,4’:3,2]-5α-androstane (**8f**)

According to Section *4.2.2., Method B,* 457 mg of **5f** was used. Eluent: EtOAc/CH_2_Cl_2_ = 10:90. White solid. Yield: 304 mg (63%); Mp 143–145 °C; ^1^H NMR (CDCl_3_, 500 MHz): *δ*H 0.74 (s, 3H, 18-H_3_), 0.75 (s, 3H, 19-H_3_), 0.80-1.00 (overlapping m, 3H), 1.06 (m, 1H), 1.23-1.47 (overlapping m, 5H), 1.54-1.65 (overlapping m, 4H), 1.73 (m, 1H), 1.80 (m, 1H), 2.05 (m, 1H), 2.10 (d, 1H, *J* = 15.2 Hz, 1α-H), 2.31 (dd, 1H, *J* = 16.5, 11.9 Hz, 4β-H), 2.43 (d, 1H, *J* = 15.2 Hz, 1β-H), 2.66 (dd, 1H, *J* = 16.5, 5.1 Hz, 4α-H), 3.63 (t, 1H, *J* = 8.6 Hz, 17α-H), 3.76 (s, 3H, N-CH_3_), 7.18 (d, 2H, *J* = 8.4 Hz), 7.59 (d, 2H, *J* = 8.4 Hz); ^13^C NMR (CDCl_3_, 125 MHz): *δ*_C_ 11.2 (C-18), 11.8 (C-19), 21.0 (C-11), 23.6 (C-15), 27.8 (C-4), 29.5 (C-6), 30.8 (C-16), 31.6 (C-7), 35.6 (C-1), 36.1 (C-8), 36.6 (C-10), 37.0 (C-12), 37.2 (N-CH_3_), 42.8 (C-5), 43.0 (C-13), 51.2 (C-14), 54.3 (C-9), 82.1 (C-17), 114.6 (C-2), 122.5 (C-4”), 129.9 (C-1”), 130.8 (2C, C-2” and C-6”), 132.1 (2C, C-3” and C-5”), 139.0 (C-5’), 147.5 (C-3); ESI-MS 485 [M+H]^+^; Anal. Calcd. for C_27_H_35_BrN_2_O C 67.07; H 7.30. Found C 66.91; H 7.29.

##### 4.2.2.8. 17β-Hydroxy-5’-(furan-2”-yl)-Γ-methylpyrazolo[3’,4’:3,2]-5α-androstane (**8g**)

According to Section *4.2.2.,Method B,* 369 mg of **5g** was used. Eluent: EtOAc/CH_2_Cl_2_ = 10:90. Light brown solid. Yield: 207 mg (52%); Mp 112–116 °C; ^1^H NMR (CDCl_3_, 500 MHz): *δ*H 0.76 (s, 3H, 18-H_3_), 0.77 (s, 3H, 19-H_3_), 0.83-1.01 (overlapping m, 3H), 1.12 (m, 1H), 1.24-1.49 (overlapping m, 5H), 1.53-1.65 (overlapping m, 4H), 1.71 (m, 1H), 1.85 (m, 1H), 2.07 (m, 1H), 2.17 (d, 1H, *J* = 15.6 Hz, 1α-H), 2.29 (dd, 1H, *J* = 16.4, 11.9 Hz, 4β-H), 2.62 (overlapping m, 2H, 1β-H and 4α-H), 3.65 (t, 1H, *J* = 8.6 Hz, 17α-H), 3.99 (s, 3H, N-CH_3_), 6.44 (d, 1H, *J* = 3.4 Hz, 3”-H), 6.51 (dd, 1H, *J* = 3.4, 1.8 Hz, 4”-H), 7.52 (d, 1H, *J* = 1.8 Hz, 5”-H); ^13^C NMR (CDCl_3_, 125 MHz): *δC* 11.2 (C-18), 11.9 (C-19), 21.0 (C-11), 23.6 (C-15), 27.6 (C-4), 29.4 (C-6), 30.7 (C-16), 31.5 (C-7), 36.0 (C-8), 36.1 (C-1), 36.4 (C-10), 37.0 (C-12), 38.6 (N-CH_3_), 42.5 (C-5), 43.0 (C-13), 51.1 (C-14), 54.3 (C-9), 82.1 (C-17), 108.5 (C-3”), 111.3 (C-4”), 114.8 (C-2), 130.8 (C-5’), 142.3 (C-5”), 145.5 (C-3), 147.1 (C-2”); ESI-MS 395 [M+H]^+^; Anal. Calcd. for C_25_H_34_N_2_O_2_ C 76.10; H 8.69. Found C 76.31; H 8.71.

##### 4.2.2.9. 17β-Hydroxy-1’-methyl-5’-(tiophen-2”-yl) -pyrazolo[3’,4’:3,2]-5α-androstane (**8h**)

According to Section *4.2.2.,Method B,* 385 mg of **5h** was used. Eluent: EtOAc/CH_2_Cl_2_ = 10:90. Light yellow solid. Yield: 237 mg (58%); Mp 124–126 °C; ^1^H NMR (CDCl_3_, 500 MHz): *δ*_H_ 0.75 (s, 3H, 18-H_3_), 0.77 (s, 3H, 19-H_3_), 0.83-1.01 (overlapping m, 3H), 1.09 (m, 1H), 1.24-1.48 (overlapping m, 5H), 1.55-1.65 (overlapping m, 4H), 1.74 (m, 1H), 1.83 (m, 1H), 2.06 (m, 1H), 2.15 (d, 1H, *J* = 15.4 Hz, 1α-H), 2.30 (dd, 1H, *J* = 16.4, 11.9 Hz, 4β-H), 2.58-2.65 (overlapping m, 2H, 1β-H and 4α-H), 3.64 (t, 1H, *J* = 8.5 Hz, 17α-H), 3.88 (s, 3H, N-CH_3_), 7.06 (d, 1H, *J* = 3.5 Hz, 5”-H), 7.14 (dd, 2H, *J* = 5.0, 3.7 Hz, 4”-H), 7.41 (d, 1H, *J* = 5.1 Hz, 3”-H); ^13^C NMR (CDCl_3_, 125 MHz): *δ*_C_ 11.2 (C-18), 11.8 (C-19), 21.0 (C-11), 23.6 (C-15), 27.8 (C-4), 29.4 (C-6), 30.8 (C-16), 31.6 (C-7), 36.0 (C-1), 36.1 (C-8), 36.6 (C-10), 37.0 (C-12), 37.6 (N-CH_3_), 42.7 (C-5), 43.0 (C-13), 51.2 (C-14), 54.3 (C-9), 82.1 (C-17), 115.6 (C-2), 126.5 (C-5”), 127.2 (C-3”), 127.5 (C-4”), 131.6 (C-2”), 133.5 (C-5’), 147.3 (C-3); ESI-MS 411 [M+H]^+^; Anal. Calcd. for C_25_H_34_N_2_OS C 73.13; H 8.35. Found C 73.09; H 8.38.

##### 4.2.2.10. 17β-Hydroxy-1’,5’-dimethylpyrazolo[3’,4’:3,2]-5α-androstane (**8l**)

According to Section *4.2.2.,Method B,* 316 mg of **5j** was used. Eluent: EtOAc/CH_2_Cl_2_ = 80:20. White solid. Yield: 207 mg (60%); Mp 240–243 °C; ^1^H NMR (CDCl_3_, 500 MHz): *δ*_H_ 0.74 (s, 3H, 18-H_3_), 0.76 (s, 3H, 19-H_3_), 0.80-1.00 (overlapping m, 3H), 1.10 (m, 1H), 1.23-1.53 (overlapping m, 5H), 1.56-1.73 (overlapping m, 5H), 1.85 (m, 1H), 1.98 (d, 1H, *J* = 14.9 Hz, 1α-H), 2.05 (m, 1H), 2.10 (s, 3H, 5’-CH_3_), 2.23 (dd, 1H, *J* = 16.4, 12.1 Hz, 4β-H) 2.43 (d, 1H, *J* = 14.9 Hz, 1β-H), 2.54 (dd, 1H, *J* = 16.4, 5.2 Hz, 4α-H), 3.64 (t, 1H, *J* = 8.6 Hz, 17α-H), 3.70 (s, 3H, N-CH_3_); ^13^C NMR (CDCl_3_, 125 MHz): *δ*_C_ 9.6 (5’-CH_3_), 11.2 (C-18), 11.8 (C-19), 21.0 (C-11), 23.6 (C-15), 27.7 (C-4), 29.4 (C-6), 30.7 (C-16), 31.5 (C-7), 35.0 (C-1), 35.9 (N-CH_3_), 36.0 (C-8), 36.4 (C-10), 37.0 (C-12), 42.8 (C-5), 43.0 (C-13), 51.1 (C-14), 54.3 (C-9), 82.1 (C-17), 113.1 (C-2), 135.0 (C-5’) 146.7 (C-3); ESI-MS 343 [M+H]^+^; Anal. Calcd. for C_22_H_34_N_2_O C 77.14; H 10.01. Found C 77.08; H 9.99.

#### 4.2.3. General Procedure for the One-Pot Synthesis of Ring A-Condensed Pyrazoles

Arylidene derivative (1.0 mmol, **5a**, **5e**, or **5i**) was dissolved in absolute EtOH (5 mL), then I2 (254 mg, 1 equiv.) and (substituted) phenylhydrazine hydrochloride (2 equiv.) were added, and the mixture was irradiated in a closed vessel at 100 °C for 2 min. After completion of the reaction, the mixture was poured into saturated aqueous solution of Na_2_S_2_O_3_ (10 mL) and extracted with CH_2_Cl_2_ (3 × 10 mL). The combined organic layers were dried over anhydrous Na_2_SO_4_ and concentrated *in vacuo.* The crude product was purified by column chromatography.

##### 4.2.3.1. 17β-Hydroxy-1’,5’-diphenylpyrazolo[3’,4’:3,2]-5α-androstane (**8i**)

According to Section *4.2.3.*, 290 mg phenylhydrazine hydrochloride and 379 mg of **5a** was used. Eluent: EtOAc/CH_2_Cl_2_ = 5:95. White solid. Yield: 394 mg (84%); Mp 224–226 °C; ^1^H NMR (CDCl_3_, 500 MHz): *δ*_H_ 0.75 (s, 3H, 18-H_3_), 0.81 (s, 3H, 19-H_3_), 0.85-1.02 (overlapping m, 3H, 9α-H, 7α-H and 14α-H), 1.09 (m, 1H, 12α-H), 1.28 (m, 1H, 15β-H), 1.35-1.48 (overlapping m, 4H, 6β-H, 8β-H, 16β-H and 11β-H), 1.64 (overlapping m, 4H, 11α-H, 5α-H, 15α-H and 6α-H), l. 75 (m, 1H, 7β-H), 1.82 (m, 1H, 12β-H), 2.07 (m, 1H, 16α-H), 2.25 (d, 1H, *J* = 15.4 Hz, 1α-H), 2.41 (dd, 1H, *J* = 16.7, 11.9 Hz, 4β-H), 2.59 (d, 1H, *J* = 15.4 Hz, 1β-H), 2.77 (dd, 1H, *J* = 16.7, 5.1 Hz, 4α-H), 3.65 (t, 1H, *J* = 8.4 Hz, 17α-H), 7.24 (overlapping m, 10H, aromatic Hs); ^13^C NMR (CDCl_3_, 125 MHz): *δ*C 11.2 (C-18), 11.9 (C-19), 21.0 (C-11), 23.6 (C-15), 27.9 (C-4), 29.4 (C-6), 30.7 (C-16), 31.5 (C-7), 35.8 (C-1), 36.0 (C-8), 36.6 (C-10), 36.9 (C-12), 42.6 (C-5), 43.0 (C-13), 51.1 (C-14), 54.3 (C-9), 82.1 (C-17), 116.4 (C-2), 124.9 (2C), 126.6 (C-4”’), 127.8 (C-4”), 128.5 (2C), 128.8 (2C), 129.4 (2C), 131.1 (C-1”), 139.1 (C-1”’), 140.6 (C-5’), 149.4 (C-3); ESI-MS 467 [M+H]^+^; Anal. Calcd. for C_32_H_38_N_2_O C 82.36; H 8.21. Found C 82.40; H 8.19.

##### 4.2.3.2. 17β-Hydroxy-5’-(4”-chlorophenyl)-1 ‘-phenylpyrazolo[3’,4’:3,2]-5α-androstane (**8j**)

According to Section *4.2.4.,* 290 mg phenylhydrazine hydrochloride and 413 mg of **5e** was used. Eluent: EtOAc/CH_2_Cl_2_ = 5:95. White solid. Yield: 397 mg (79%); Mp 185–188 °C; ^1^H NMR (CDCl_3_, 500 MHz): *δ*_H_ 0.75 (s, 3H, 18-H_3_), 0.80 (s, 3H, 19-H_3_), 0.85-1.02 (overlapping m, 3H, 9α-H, 7α-H and 14α-H), 1.09 (m, 1H, 12α-H), 1.28 (m, 1H, 15β-H), 1.35-1.48 (overlapping m, 4H, 6β-H, 8β-H, 16β-H and 11β-H), 1.59-1.68 (overlapping m, 4H, 11α-H, 5α-H, 15α-H and 6α-H), 1.75 (m, 1H, 7β-H), 1.83 (m, 1H, 12β-H), 2.07 (m, 1H, 16α-H), 2.22 (d, 1H, *J* = 15.4 Hz, 1α-H), 2.40 (dd, 1H, *J* = 16.7, 12.1 Hz, 4β-H), 2.54 (d, 1H, *J* = 15.4 Hz, 1β-H), 2.76 (dd, 1H, *J* = 16.7, 5.1 Hz, 4α-H), 3.65 (m, 1H, 17α-H), 7.08 (d, 2H, *J* = 8.5 Hz, 3”-H and 5”-H), 7.22 (m, 3H), 7.27-7.30 (m, 4H); ^13^C NMR (CDCl_3_, 125 MHz): *δ*C 11.2 (C-18), 11.9 (C-19), 21.0 (C-11), 23.6 (C-15), 27.8 (C-4), 29.4 (C-6), 30.7 (C-16), 31.5 (C-7), 35.8 (C-1), 36.0 (C-8), 36.5 (C-10), 36.9 (C-12), 42.6 (C-5), 43.0 (C-13), 51.1 (C-14), 54.2 (C-9), 82.1 (C-17), 116.6 (C-2), 124.9 (2C, C-2”’ and C-6”’), 126.9 (C-4”’), 128.9 (2C), 129.0 (2C), 129.5 (C-1”), 130.6 (2C), 133.9 (C-4”), 137.9 (C-1”’), 140.3 (C-5’), 149.6 (C-3); ESI-MS 501 [M+H]^+^; Anal. Calcd. for C_32_H_37_ClN_2_O C 76.70; H 7.44. Found C 76.62; H 7.46.

##### 4.2.3.3. 17β-Hydroxy-5’-(4”-hydroxyphenyl)-1 ‘-phenylpyrazolo[3’,4’:3,2]-5α-androstane (**8k**)

According to Section *4.2.4.,* 290 mg phenylhydrazine hydrochloride and 395 mg of **5i** was used. Eluent: EtOAc/CH_2_Cl_2_ = 20:80. Light brown solid. Yield: 433 mg (90%); Mp 290–292 °C; ^1^H NMR (DMSO-*d*_6_, 500 MHz): *δ*H 0.64 (s, 3H, 18-H_3_), 0.71 (s, 3H, 19-H_3_), 0.82-1.01 (overlapping m, 4H), 1.18 (m, 1H), 1.28-1.35 (overlapping m, 4H), 1.50-1.60 (overlapping m, 4H), 1.67 (m, 1H), 1.74 (m, 1H), 1.84 (m, 1H), 2.19 (d, 1H, *J* = 15.4 Hz, 1α-H), 2.25 (dd, 1H, *J* = 16.7, 12.4 Hz, 4β-H), 2.42 (d, 1H, *J* = 15.4 Hz, 1β-H), 2.61 (dd, 1H, *J* = 16.7, 5.1 Hz, 4α-H), 3.44 (m, 1H, 17α-H), 4.40 (d, 1H, *J* = 4.82, 17-OH), 6.73 (d, 2H, *J* = 8.6 Hz, 3”-H and 5”-H), 6.94 (d, 2H, *J* = 8.6 Hz, 2”-H and 6”-H), 7.16 (d, 2H, *J* = 7.5 Hz, 3”’-H and 5”’-H), 7.23 (t-like m, 1H, 4”’-H), 7.31 (t-like m, 2H, 2”’-H and 6”’-H), 9.65 (s, 1H, 4’-OH); ^13^C NMR (DMSO-*d*_6_, 125 MHz): *δC* 11.2 (C-18), 11.6 (C-19), 20.4 (C-11), 23.1 (C-15), 27.3 (C-4), 28.8 (C-6), 29.8 (C-16), 31.0 (C-7), 35.0 (C-1), 35.4 (C-8), 35.8 (C-10), 36.5 (C-12), 41.8 (C-5), 42.4 (C-13), 50.5 (C-14), 53.4 (C-9), 80.0 (C-17), 115.1 (C-2), 115.4 (2C, C-3” and C-5”), 120.8 (C-1”), 124.2 (2C, C-2”’ and C-6”’), 126.3 (C-4”’), 128.6 (2C), 130.2 (2C), 138.6 (C-1”’), 140.3 (C-5’), 148.0 (C-3), 157.1 (C-4”); ESI-MS 483 [M+H]^+^; Anal. Calcd. for C_32_H_38_N_2_O_2_ C 79.63; H 7.94. Found C 79.81; H 7.96.

#### 4.2.4. General procedure for the synthesis of heterocyclic 17-keto steroids by Jones oxidation

The crude product (**8a–l**) of the heterocyclization (*4.2.3.*) was dissolved in acetone (10 mL) and Jones reagent (0.2 mL) was added dropwise into the solution, which was then stirred at room temperature for 30 min, after which it was poured into ice-cold water. NH4Cl was added and the resulting precipitate was filtered off and dried. The crude product was purified by column chromatography.

##### 4.2.4.1. 1’-methyl-5’-phenylpyrazolo[3’,4’:3,2]-5α-androstan-17-one (**10a**)

Eluent: EtOAc/CH_2_Cl_2_ = 10:90. White solid. Yield: 307 mg (76%); Mp 200–203 °C; ^1^H NMR (CDCl_3_, 500 MHz): *δ*H 0.77 (s, 3H, 19-H_3_), 0.86 (s, 3H, 18-H_3_), 0.89 (m, 1H), 1.04 (m, 1H), 1.22-1.45 (overlapping m, 5H), 1.50-1.71 (overlapping m, 4H), 1.80 (m, 1H), 1.85 (m, 1H), 1.97 (m, 1H), 2.08 (m, 1H, 16α-H), 2.15 (d, 1H, *J* = 15.2 Hz, 1α-H), 2.33 (dd, 1H, *J* = 16.5, 12.0 Hz, 4β-H), 2.45 (dd, 1H, *J* = 19.4, 8.6 Hz, 16β-H), 2.46 (d, 1H, *J* = 15.2 Hz, 1β-H), 2.69 (dd, 1H, *J* = 16.5, 5.2 Hz, 4α-H), 3.78 (s, 3H, N-CH_3_), 7.31 (d-like m, 2H, 3”-H and 5”-H), 7.39 (t-like m, 1H, 4”-H), 7.46 (t-like m, 2H, 2”-H and 6”-H); ^13^C NMR (CDCl_3_, 125 MHz): *δ*_C_ 11.7 (C-19), 13.9 (C-18), 20.6 (C-11), 22.0 (C-15), 27.8 (C-4), 29.3 (C-6), 30.8 (C-16), 31.7 (C-7), 35.5 (2C, C-8 and C-1), 36.0 (C-12), 36.6 (C-10), 37.2 (N-CH_3_), 42.7 (C-5), 47.8 (C-13), 51.6 (C-14), 54.2 (C-9), 114.1 (C-2), 128.2 (C-4”), 128.8 (2C, C-3” and C-5”), 129.3 (2C, C-2” and C-6”), 130.9 (C-1”), 140.2 (C-5’) 147.1 (C-3), 221.4 (C-17); ESI-MS 403 [M+H]^+^; Anal. Calcd. for C_27_H_34_N_2_O C 80.55; H 8.51. Found C 80.65; H 8.52.

##### 4.2.4.2. 1’-methyl-5’-(4”-tolyl)-pyrazolo[3’,4’:3,2]-5α-androstan-17-one (**10b**)

Eluent: EtOAc/CH_2_Cl_2_ = 10:90. White solid. Yield: 307 mg (74%); Mp 180–183 °C; ^1^H NMR (CDCl_3_, 500 MHz): *δ*_H_ 0.76 (s, 3H, 19-H_3_), 0.86 (s, 3H, 18-H_3_), 0.88 (m, 1H), 1.03 (m, 1H), 1.21-1.44 (overlapping m, 5H), 1.48-1.71 (overlapping m, 4H), 1.80 (m, 1H), 1.85 (m, 1H), 1.97 (m, 1H), 2.06 (m, 1H, 16α-H), 2.14 (d, 1H, *J* = 15.4 Hz, 1α-H), 2.33 (dd, 1H, *J* = 16.4, 12.2 Hz, 4β-H), 2.41 (s, 4”-CH_3_), 2.43 (m, 1H, 16β-H), 2.47 (d, 1H, *J* = 15.4 Hz, 1β-H), 2.68 (dd, 1H, *J* = 16.4, 5.2 Hz, 4α-H), 3.77 (s, 3H, N-CH_3_), 7.20 (d, 2H, *J* = 8.0 Hz, 2”-H and 6”-H), 7.27 (d, 2H, *J* = 8.4 Hz, 3”-H and 5”-H); ^13^C NMR (CDCl_3_, 125 MHz): *δ*_C_ 11.7 (C-19), 13.9 (C-18), 20.6 (C-11), 21.4 (4”-CH_3_), 22.0 (C-15), 27.8 (C-4), 29.3 (C-6), 30.8 (C-16), 31.7 (C-7), 35.5 (C-8), 35.6 (C-1), 36.0 (C-12), 36.6 (C-10), 37.2 (N-CH_3_), 42.7 (C-5), 47.8 (C-13), 51.6 (C-14), 54.2 (C-9), 113.9 (C-2), 127.9 (C-4”), 129.1 (2C, C-2” and C-6”), 129.5 (2C, C-3” and C-5”), 138.1 (C-1”), 140.2 (C-5’) 147.0 (C-3), 221.5 (C-17); ESI-MS 417 [M+H]^+^; Anal. Calcd. for C_28_H_36_N_2_O C 80.73; H 8.71. Found C 80.75; H 8.70.

##### 4.2.4.3. 5’-(4”-methoxyphenyl)-1’-methylpyrazolo[3’,4’:3,2]-5α-androstan-17-one (**10c**)

Eluent: EtOAc/CH_2_Cl_2_ = 20:80. White solid. Yield: 321 mg (74%); Mp 102–105 °C; ^1^H NMR (CDCl_3_, 500 MHz): *δ*_H_ 0.77 (s, 3H, 19-H_3_), 0.86 (s, 3H, 18-H_3_), 0.88 (m, 1H), 1.04 (m, 1H), 1.22-1.44 (overlapping m, 5H), 1.48-1.70 (overlapping m, 4H), 1.80 (m, 1H), 1.86 (m, 1H), 1.97 (m, 1H), 2.07 (m, 1H, 16α-H), 2.13 (d, 1H, *J* = 15.2 Hz, 1α-H), 2.33 (dd, 1H, *J* = 16.5, 12.1 Hz, 4β-H), 2.42-2.48 (overlapping m, 2H, 16β-H and 1β-H), 2.68 (dd, 1H, *J* = 16.5, 5.2 Hz, 4α-H), 3.76 (s, 3H, N-CH_3_), 3.86 (s, 3H, 4”-OCH_3_), 6.99 (d, 2H, *J* = 8.7 Hz, 3”-H and 5”-H), 7.23 (d, 2H, *J* = 8.7 Hz, 2”-H and 6”-H); ^13^C NMR (CDCl_3_, 125 MHz): *δ*_C_ 11.7 (C-19), 13.9 (C-18), 20.6 (C-11), 22.0 (C-15), 27.8 (C-4), 29.3 (C-6), 30.8 (C-16), 31.7 (C-7), 35.5 (C-8), 35.6 (C-1), 36.0 (C-12), 36.6 (C-10), 37.1 (N-CH_3_), 42.7 (C-5), 47.8 (C-13), 51.6 (C-14), 54.2 (C-9), 55.5 (4”-CH_3_), 113.8 (C-2), 114.3 (2C, C-3” and C-5”), 123.1 (C-1”), 130.5 (2C, C-2” and C-6”), 140.0 (C-5’), 147.0 (C-3) 159.6 (C-4”), 221.5 (C-17); ESI-MS 433 [M+H]^+^; Anal. Calcd. for C_28_H_36_N_2_O_2_ C 77.74; H 8.39. Found C 77.80; H 8.37.

##### 4.2.4.4. 5’-(4”-fluorophenyl)-Γ-methylpyrazolo[3’,4’:3,2]-5α-androstan-17-one (**10d**)

Eluent: EtOAc/CH_2_Cl_2_ = 20:80. White solid. Yield: 251 mg (60%); Mp 185–187 °C; ^1^H NMR (CDCl_3_, 500 MHz): *δ*_H_ 0.77 (s, 3H, 19-H_3_), 0.87 (s, 3H, 18-H_3_), 0.88 (m, 1H), 1.04 (m, 1H), 1.22-1.44 (overlapping m, 5H), 1.48-1.71 (overlapping m, 4H), 1.80 (m, 1H), 1.86 (m, 1H), 1.97 (m, 1H), 2.07 (m, 1H, 16α-H), 2.12 (d, 1H, *J* = 15.4 Hz, 1α-H), 2.33 (dd, 1H, *J* = 16.5, 12.1 Hz, 4β-H), 2.43 (d, 1H, *J* = 15.4 Hz, 1β-H), 2.44 (dd, 1H, *J* = 19.2, 8.8 Hz, 16β-H), 2.68 (dd, 1H, *J* = 16.5, 5.2 Hz, 4α-H), 3.75 (s, 3H, N-CH_3_), 7.16 (t-like m, 2H, 2”-H and 6”-H), 7.27 (m, 2H, 3”-H and 5”-H); ^13^C NMR (CDCl_3_, 125 MHz): *δ*C 11.7 (C-19), 13.8 (C-18), 20.6 (C-11), 22.0 (C-15), 27.8 (C-4), 29.2 (C-6), 30.8 (C-16), 31.7 (C-7), 35.5 (C-1), 35.5 (C-8), 36.0 (C-12), 36.6 (C-10), 37.1 (N-CH_3_), 42.6 (C-5), 47.8 (C-13), 51.6 (C-14), 54.2 (C-9), 114.2 (C-2), 115.9 (d, 2C, *J* = 21.7 Hz, C-3” and C-5”), 126.9 (d, *J* = 3.3 Hz, C-1”), 131.1 (d, 2C, *J* = 8.2 Hz, C-2” and C-6”), 139.2 (C-5’), 147.2 (C-3), 162.7 (d, *J* = 248.5 Hz, C-4”), 221.4 (C-17); ESI-MS 421 [M+H]^+^; Anal. Calcd. for C_27_H_33_FN_2_O C 77.11; H 7.91. Found C 77.05; H 7.90.

##### 4.2.4.5. 5’-(4”-chlorophenyl)-1’-methylpyrazolo[3’,4’:3,2]-5α-androstan-17-one (**10e**)

Eluent: EtOAc/CH_2_Cl_2_ = 20:80. White solid. Yield: 280 mg (64%); Mp 211–215 °C; ^1^H NMR (CDCl_3_, 500 MHz): *δ*_H_ 0.76 (s, 3H, 19-H_3_), 0.86 (s, 3H, 18-H_3_), 0.88 (m, 1H), 1.04 (m, 1H), 1.23-1.44 (overlapping m, 5H), 1.48-1.73 (overlapping m, 4H), 1.79-1.87 (overlapping m, 2H), 1.97 (m, 1H), 2.06 (m, 1H, 16α-H), 2.11 (d, 1H, *J* = 14.7 Hz, 1α-H), 2.32 (dd, 1H, *J* = 16.5, 12.4 Hz, 4β-H), 2.42 (d, 1H, *J* = 14.7 Hz, 1β-H), 2.45 (m, 1H, 16β-H), 2.68 (dd, 1H, *J* = 16.5, 5.1 Hz, 4α-H), 3.75 (s, 3H, N-CH_3_), 7.23 (d, 2H, *J* = 8.2 Hz, 3”-H and 5”-H), 7.43 (d, 2H, *J* = 8.2 Hz, 2”-H and 6”-H); ^13^C NMR (CDCl_3_, 125 MHz): *δ*C 11.7 (C-19), 13.9 (C-18), 20.7 (C-11), 22.0 (C-15), 27.8 (C-4), 29.3 (C-6), 30.8 (C-16), 31.8 (C-7), 35.6 (C-8), 35.6 (C-1), 36.0 (C-12), 36.7 (C-10), 37.2 (N-CH_3_), 42.7 (C-5), 47.8 (C-13), 51.6 (C-14), 54.3 (C-9), 114.4 (C-2), 129.1 (2C, C-2” and C-6”), 129.3 (C-1”), 130.5 (2C, C-3” and C-5”), 134.4 (C-4”), 139.1 (C-5’) 147.3 (C-3), 221.1 (C-17); ESI-MS 437 [M+H]^+^; Anal. Calcd. for C_27_H_33_ClN_2_O C 74.21; H 7.61. Found C 77.31; H 7.59.

##### 4.2.4.6. 5’-(4”-bromophenyl)-1’-methylpyrazolo[3’,4’:3,2]-5α-androstan-17-one (**10f**)

Eluent: EtOAc/CH_2_Cl_2_ = 20:80. White solid. Yield: 273 mg (57%); Mp 126–128 °C; ^1^H NMR (CDCl_3_, 500 MHz): *δ*H 0.76 (s, 3H, 19-H_3_), 0.87 (s, 3H, 18-H_3_), 0.88 (m, 1H), 1.04 (m, 1H), 1.22-1.44 (overlapping m, 5H), 1.48-1.73 (overlapping m, 4H), 1.81 (m, 1H), 1.86 (m, 1H), 1.97 (m, 1H), 2.07 (m, 1H, 16α-H), 2.13 (d, 1H, *J* = 15.2 Hz, 1α-H), 2.33 (dd, 1H, *J* = 16.5, 12.1 Hz, 4β-H), 2.42 (d, 1H, *J* = 15.2 Hz, 1β-H), 2.45 (dd, 1H, *J* = 16.5, 5.3 Hz, 16β-H), 2.68 (dd, 1H, *J* = 16.5, 5.3 Hz, 4α-H), 3.76 (s, 3H, N-CH_3_), 7.18 (d, 2H, *J* = 8.4 Hz, 2”-H and 6”-H), 7.60 (d, 2H, *J* = 8.4 Hz, 3”-H and 5”-H); ^13^C NMR (CDCl_3_, 125 MHz): *δ*C 11.7 (C-19), 13.9 (C-18), 20.6 (C-11), 22.0 (C-15), 27.7 (C-4), 29.2 (C-6), 30.8 (C-16), 31.7 (C-7), 35.5 (C-8), 35.5 (C-1), 36.0 (C-12), 36.6 (C-10), 37.2 (N-CH_3_), 42.6 (C-5), 47.8 (C-13), 51.6 (C-14), 54.2 (C-9), 114.3 (C-2), 122.5 (C-4”), 129.7 (C-1”), 130.8 (2C, C-2” and C-6”), 132.1 (2C, C-3” and C-5”), 139.0 (C-5’) 147.3 (C-3), 221.4 (C-17); ESI-MS 483 [M+H]^+^; Anal. Calcd. for C_27_H_33_BrN_2_O C 67.35; H 6.91. Found C 67.37; H 6.93.

##### 4.2.4.7. 5’-(furan-2”-yl)-1’-methylpyrazolo[3’,4’:3,2]-5α-androstan-17-one (**10g**)

Eluent: EtOAc/CH_2_Cl_2_ = 5:95. Light brown solid. Yield: 180 mg (46%); Mp > 80 °C (decomp.); ^1^H NMR (CDCl_3_, 500 MHz): *δ*H 0.78 (s, 3H, 19-H_3_), 0.89 (s, 3H, 18-H_3_), 0.93 (m, 1H), 1.04 (m, 1H), 1.25-1.70 (overlapping m, 9H), 1.78 (m, 1H), 1.86 (m, 1H), 1.98 (m, 1H), 2.09 (m, 1H, 16α-H), 2.18 (d, 1H, *J* = 15.6 Hz, 1α-H), 2.31 (dd, 1H, *J* = 16.4, 12.1 Hz, 4β-H), 2.46 (dd, 1H, *J* = 19.2, 8.8 Hz, 16β-H), 2.63 (dd, 1H, *J* = 16.4, 5.2 Hz, 4α-H), 2.66 (d, 1H, *J* = 15.6 Hz, 1β-H), 3.99 (s, 3H, N-CH_3_), 6.44 (d, 1H, *J* = 3.3 Hz, 3”-H), 6.51 (dd, 1H, *J* = 3.3, 1.8 Hz, 4”-H), 7.52 (d, 1H, *J* = 1.3 Hz, 5”-H); ^13^C NMR (CDCl_3_, 125 MHz): *δ*C 11.9 (C-19), 13.9 (C-18), 20.7 (C-11), 22.0 (C-15), 27.5 (C-4), 29.2 (C-6), 30.8 (C-16), 31.8 (C-7), 35.5 (C-8), 36.0 (C-1), 36.0 (C-12), 36.5 (C-10), 38.6 (N-CH_3_), 42.4 (C-5), 47.8 (C-13), 51.6 (C-14), 54.2 (C-9), 108.6 (C-3”), 111.4 (C-4”), 114.6 (C-2), 130.9 (C-5’), 142.3 (C-5”), 145.4 (C-3), 147.0 (C-2”), 221.4 (C-17); ESI-MS 393 [M+H]^+^; Anal. Calcd. for C_25_H_32_N_2_O_2_ C 76.49; H 8.22. Found C 76.37; H 8.20.

##### 4.2.4.8. 1’-methyl-5’-(tiophen-2”-yl)-pyrazolo[3’,4’:3,2]-5α-androstan-17-one (**10h**)

Eluent: EtOAc/CH_2_Cl_2_ = 5:95. Light yellow solid. Yield: 218 mg (53%); Mp 90–2 °C; ^1^H NMR (CDCl_3_, 500 MHz): *δ*_H_ 0.77 (s, 3H, 19-H_3_), 0.87 (s, 3H, 18-H_3_), 0.91 (m, 1H), 1.04 (m, 1H), 1.25-1.73 (overlapping m, 9H), 1.84 (overlapping m, 2H), 1.97 (m, 1H), 2.08 (m, 1H, 16α-H), 2.16 (d, 1H, *J* = 15.4 Hz, 1α-H), 2.31 (dd, 1H, *J* = 16.4, 12.1 Hz, 4β-H), 2.45 (dd, 1H, *J* = 19.2, 8.8 Hz, 16β-H), 2.60 (d, 1H, *J* = 15.3 Hz, 1β-H), 2.65 (dd, 1H, *J* = 16.4, 5.2 Hz, 4α-H), 3.88 (s, 3H, N-CH_3_), 7.07 (d, 1H, *J* = 3.5 Hz, 5”-H), 7.14 (dd, 1H, *J* = 5.1, 3.6 Hz, 4”-H), 7.43 (d, 1H, *J* = 5.1 Hz, 3”-H); ^13^C NMR (CDCl_3_, 125 MHz): *δ*C 11.8 (C-19), 13.9 (C-18), 20.7 (C-11), 22.0 (C-15), 27.7 (C-4), 29.2 (C-6), 30.8 (C-16), 31.7 (C-7), 35.5 (C-8), 35.9 (C-1), 36.0 (C-12), 36.6 (C-10), 37.7 (N-CH_3_), 42.5 (C-5), 47.8 (C-13), 51.5 (C-14), 54.2 (C-9), 115.3 (C-2), 126.5 (C-5”), 127.2 (C-3”), 127.6 (C-4”), 131.3 (C-2”), 133.6 (C-5’), 147.1 (C-3), 221.4 (C-17); ESI-MS 409 [M+H]^+^; Anal. Calcd. for C_25_H_32_N_2_OS C 73.49; H 7.89. Found C 73.53; H 7.88.

##### 4.2.4.9. 1’,5’-diphenylpyrazolo[3’,4’:3,2]-5α-androstan-17-one (**10i**)

According to Section *4.2.4.,* 117 mg of **8i** was used. Eluent: EtOAc/CH_2_Cl_2_ = 2:98. White solid. Yield: 110 mg (95%); Mp 267-270 °C; ^1^H NMR (CDCl_3_, 500 MHz): *δ*_H_ 0.82 (s, 3H, 19-H_3_), 0.88 (s, 3H, 18-H_3_), 0.94 (m, 1H), 1.07 (m, 1H), 1.25-1.34 (overlapping m, 2H), 1.38-1.61 (overlapping m, 4H), 1.70 (m, 3H), 1.83 (m, 1H), 1.88 (m, 1H), 1.99 (m, 1H), 2.09 (m, 1H, 16α-H), 2.26 (d, 1H, *J* = 15.4 Hz, 1α-H), 2.40-2.49 (overlapping m, 2H, 4β-H, 16β-H), 2.59 (d, 1H, *J* = 15.4 Hz, 1β-H), 2.80 (dd, 1H, *J* = 16.7, 5.1 Hz, 4α-H), 7.15-7.34 (overlapping m, 10H, aromatic Hs); ^13^C NMR (CDCl_3_, 125 MHz): *δ*C 11.8 (C-19), 13.9 (C-18), 20.7 (C-11), 22.0 (C-15), 27.9 (C-4), 29.3 (C-6), 30.8 (C-16), 31.7 (C-7), 35.5 (C-8), 35.7 (C-1), 36.0 (C-12), 36.6 (C-10), 42.6 (C-5), 47.8 (C-13), 51.6 (C-14), 54.2 (C-9), 116.2 (C-2), 124.9 (2C), 126.7 (C-4”’), 127.9 (C-4”), 128.6 (2C), 128.8 (2C), 129.4 (2C), 131.0 (C-1”), 139.2 (C-1”’), 140.6 (C-5’), 149.2 (C-3), 221.4 (C-17); ESI-MS 465 [M+H]^+^; Anal. Calcd. for C_32_H_36_N_2_O C 82.72; H 7.81. Found C 82.54; H 7.80.

##### 4.2.4.10. 5’-(4”-chlorophenyl)-1 ‘-phenylpyrazolo[3 ‘,4 ‘:3,2]-5α-androstan-17-one (**10j**)

According to Section *4.2.5.,* 125 mg of **8j** was used. Eluent: CH_2_Cl_2_. White solid. Yield: 114 mg (91%); Mp > 250 °C (decomp.); ^1^H NMR (CDCl_3_, 500 MHz): *δ*_H_ 0.81 (s, 3H, 19-H_3_), 0.88 (s, 3H, 18-H_3_), 0.94 (m, 1H), 1.07 (m, 1H), 1.24-1.34 (overlapping m, 2H), 1.39-1.59 (overlapping m, 4H), 1.70 (m, 3H), 1.82-1.90 (overlapping m, 2H), 1.99 (m, 1H), 2.09 (m, 1H, 16α-H), 2.24 (d, 1H, *J* = 15.4 Hz, 1α-H), 2.39-2.49 (overlapping m, 2H, 4β-H, 16β-H), 2.55 (d, 1H, *J* = 15.4 Hz, 1β-H), 2.79 (dd, 1H, *J* = 16.7, 5.1 Hz, 4α-H), 7.08 (d, 2H, *J* = 8.5 Hz, 3”-H and 5”-H), 7.25 (m, 7H); ^13^C NMR (CDCl_3_, 125 MHz): *δ*_C_ 11.8 (C-19), 13.9 (C-18), 20.7 (C-11), 22.0 (C-15), 27.8 (C-4), 29.2 (C-6), 30.8 (C-16), 31.7 (C-7), 35.5 (C-8), 35.7 (C-1), 36.0 (C-12), 36.6 (C-10), 42.5 (C-5), 47.8 (C-13), 51.5 (C-14), 54.2 (C-9), 116.4 (C-2), 124.9 (2C, C-2”’ and C-6”’), 127.0 (C-4”’), 128.9 (2C), 129.0 (2C), 129.4 (C-1”), 130.6 (2C), 134.0 (C-4”), 138.0 (C-1”’), 140.3 (C-5’), 149.4 (C-3), 221.3 (C-17); ESI-MS 499 [M+H]^+^; Anal. Calcd. for C_32_H_35_ClN_2_O C 77.01; H 7.07. Found C 77.20; H 7.05.

##### 4.2.4.11. 5’-(4”-hydroxyphenyl)-1 ‘-phenylpyrazolo[3’,4 ‘:3,2]-5α-androstan-17-one (**10k**)

According to Section *4.2.5.,* 121 mg of **8k** was used. Eluent: EtOAc/CH_2_Cl_2_ = 10:90. White solid. Yield: 109 mg (91%); Mp > 250 °C (decomp.); ^1^H NMR (CDCl_3_, 500 MHz): *δ*_H_ 0.81 (s, 3H, 19-H_3_), 0.89 (s, 3H, 18-H_3_), 0.93 (m, 1H), 1.06 (m, 1H), 1.26-1.35 (overlapping m, 2H), 1.39-1.47 (overlapping m, 2H), 1.51-1.61 (overlapping m, 2H), 1.69 (m, 3H), 1.82-1.89 (overlapping m, 2H), 1.99 (m, 1H), 2.10 (m, 1H, 16α-H), 2.22 (d, 1H, *J* = 15.3 Hz, 1α-H), 2.38-2.50 (overlapping m, 2H, 4β-H, 16β-H), 2.57 (d, 1H, *J* = 15.3 Hz, 1β-H), 2.77 (dd, 1H, *J* = 16.7, 5.0 Hz, 4α-Hz), 6.58 (bs, 1H, 4”-OH), 6.75 (d, 2H, *J* = 8.6 Hz, 3”-H and 5”-H), 6.98 (d, 2H, *J* = 8.6 Hz, 2”-H and 6”-H), 7.22 (m, 5H); ^13^C NMR (CDCl_3_, 125 MHz): *δ*_C_ 11.9 (C-19), 13.9 (C-18), 20.7 (C-11), 22.0 (C-15), 27.7 (C-4), 29.2 (C-6), 30.8 (C-16), 31.7 (C-7), 35.5 (C-8), 35.7 (C-1), 36.1 (C-12), 36.6 (C-10), 42.5 (C-5), 47.9 (C-13), 51.6 (C-14), 54.2 (C-9), 115.6 (C-2), 115.7 (2C, C-3” and C-5”), 122.7 (C-1”), 125.0 (2C, C-2”’ and C-6”’), 126.7 (C-4”’), 128.8 (2C), 130.8 (2C), 139.4 (C-1”’), 140.4 (C-5’), 149.2 (C-3), 156.1 (C-4”), 222.0 (C-17); ESI-MS 481 [M+H]^+^; Anal. Calcd. for C_32_H_36_N_2_O_2_ C 79.96; H 7.55. Found C 79.79; H 7.57.

##### 4.2.4.12. 1’,5’-dimethylpyrazolo[3’,4’:3,2]-5α-androstan-17-one (**10l**)

Eluent: EtOAc/CH_2_Cl_2_ = 50:50. Light yellow solid. Yield: 184 mg (54%); Mp 90–92 °C; ^1^H NMR (CDCl_3_, 500 MHz): *δ*_H_ 0.76 (s, 3H, 19-H_3_), 0.89 (s, 3H, 18-H_3_), 0.90 (m, 1H), 1.03 (m, 1H), 1.25-1.78 (overlapping m, 10H), 1.84 (overlapping m, 1H), 1.97 (m, 1H), 1.99 (d, 1H, *J* = 14.8Hz, 1α-H), 2.08 (m, 1H, 16α-H), 2.11 (s, 3H, 5’-CH_3_), 2.25 (dd, 1H, *J* = 16.4, 12.1 Hz, 4β-H), 2.43 (d, 1H, *J* = 14.8 Hz, 1β-H), 2.45 (dd, 1H, *J* = 19.2, 8.8 Hz, 16β-H), 2.57 (dd, 1H, *J* = 16.4, 5.2 Hz, 4α-H), 3.71 (s, 3H, N-CH_3_); ^13^C NMR (CDCl_3_, 125 MHz): *δ*_C_ 9.6 (5’-CH_3_), 11.8 (C-19), 13.9 (C-18), 20.7 (C-11), 22.0 (C-15), 27.7 (C-4), 29.3 (C-6), 30.8 (C-16), 31.8 (C-7), 35.0 (C-1), 35.5 (C-8), 35.9 (N-CH_3_), 36.0 (C-12), 36.5 (C-10), 42.8 (C-5), 47.8 (C-13), 51.6 (C-14), 54.3 (C-9), 112.9 (C-2), 135.1 (C-5’), 146.5 (C-3), 221.5 (C-17); ESI-MS 341 [M+H]^+^; Anal. Calcd. for C_22_H_32_N_2_O C 77.60; H 9.47. Found C 77.55; H 9.48.

### 4.3. Cell lines

The ARE14 reporter cell line [20] (kindly gifted by prof. Zdeněk Dvořák from Palacky University Olomouc, Czech Republic) and the LNCaP cells (purchased from ECACC) were grown in RPMI-1640 medium. The 22Rv1, DuCaP, LAPC-4 cell lines (kindly gifted by prof. Zoran Culig, Innsbruck Medical University) and DU145 (purchased from ECACC) were grown in DMEM medium. All media were supplemented with 10% normal or charcoal-stripped fetal bovine serum (steroid-depleted serum), 100 IU/ml penicillin, 100 μg/ml streptomycin, 4 mM glutamine and 1 mM sodium pyruvate. Cells were cultivated in a humidified incubator at 37 °C and in 5% CO_2_ atmosphere.

### 4.4. AR transcriptional luciferase assay

ARE14 cells were seeded (40 000 cells/well) into the Nunc™ MicroWell™ 96-well optical flat-bottom plate (Thermo Fisher Scientific) for luciferase assay. The second day, the cultivation medium (supplemented with FBS) was discarded, and cells were washed with PBS. Cells were then incubated in presence of analysed compounds dissolved in medium supplemented with CSS (agonist mode) or CSS with 1 nM R1881 (antagonist mode), including CSS and 1 nM R1881 controls. Upon 24 h of incubation, cells were washed with PBS again and lysed for 10 min in a lysis buffer (10 mM Tris pH = 7.4, 2 mM DCTA, 1% nonidet P40, 2 mM DTT) at 37 °C. Next, reaction buffer (20 mM tricine pH = 7.8, 1.07 mM MgSO_4_ · 7H_2_0, 5 mM ATP, 9.4 μM luciferin) was added and the luminescence of the samples was measured using a Tecan M200 Pro microplate reader (Biotek).

### 4.4. Cell viability assay

Cells were seeded into the 96-well tissue culture plates, the other day, solutions of compounds were added in different concentrations in duplicate for 72 h. Upon treatment, the resazurin solution (Sigma Aldrich) was added for 4 h, and then the fluorescence of resorufin was measured at 544 nm/590 nm (excitation/emission) using a Fluoroskan Ascent microplate reader (Labsystems). Percentual viability was calculated and in the separate experiment, GI_50_ value was calculated from the dose response curves that resulted from the measurements using GraphPad Prism 5.

### 4.5. Colony formation assay

PCa cells 22Rv1 and DU-145 (5000 cells per well), LAPC-4 and PC-3 (10 000 cells per well) were seeded into 6-well plates. After two days of cultivation, the medium was replaced with fresh medium containing different concentrations of the compounds. Cells were cultivated for 10 days at the presence of compounds. After the treatment, the medium was discarded and colonies were fixed with 70% ethanol for 15 min, washed with PBS and stained with crystal violet (1% solution in 96% ethanol) for 1 h. Finally, wells were washed with PBS and colonies’ photograph was captured.

### 4.6. Immunoblotting

After all treatments, cells were harvested by centrifugation, washed twice with PBS, pelleted and kept frozen at −80 °C. Pellets were thawed, resuspended in ice-cold RIPA lysis buffer supplemented with phosphatase and protease inhibitors. Ultrasound sonication (10 sec with 30% amplitude) of cells was performed on ice and soluble proteins in supernatants we obtained by centrifugation at 14.000 *g* for 30 min. Protein concentration in supernatants was measured and balanced within samples. Proteins were denatured by addition of SDS-loading buffer, separated by SDS-PAGE and electroblotted onto nitrocellulose membranes. Immunodetection of proteins was performed as usual, membranes were blocked in BSA solution, incubated overnight with primary antibodies, washed and incubated with secondary antibodies conjugated with peroxidase. Then, peroxidase activity was detected by SuperSignal West Pico reagents (Thermo Scientific) using a CCD camera LAS-4000 (Fujifilm). Primary antibodies purchased from Merck (anti-α-tubulin, clone DM1A; anti-phosphorylated AR (S81)). Primary antibodies purchased from Santa Cruz Biotechnology (anti-β-actin, clone C4). Specific antibodies purchased from Cell Signaling Technology (anti-AR, clone D6F11; anti-PSA/KLK3, clone D6B1; anti-Nkx3.1, clone D2Y1A); anti-rabbit secondary antibody (porcine anti-rabit immunoglobulin serum); anti-mouse secondary antibody (rabbit anti-mouse IgG, clone D3V2A)). All antibodies were diluted in 4% BSA and 0.1% Tween 20 in TBS.

### 4.7. Analyses of mRNA expression

Cells were treated and harvested into lysis buffer and total RNA was isolated using RNeasy plus mini kit (QIAGEN) based on the manufacturer’s instruction. RNA concentration and purity was evaluated using DeNovix DS-11 spectrophotometer, while quality of RNA was determined by gel electrophoresis (high-quality samples displayed intact ribosomal RNA). The RNA (0.5–1 μg) of those samples was used for reverse transcription into first-strand cDNA which was carried out by SensiFast cDNA Synthesis Kit (Bioline). RNA Spike I template (TATAA) was used as a transcriptional inhibition control. Quantitative RT-PCR was performed on CFX96 Real-Time PCR Detection System (Biorad) with a SensiFAST SYBR No-Rox Kit (Bioline). The suitable primers were designed using Primer-BLAST [28] and synthesized by Generi Biotech. Primary data were analyzed using Bio-rad CFX Maestro 2.2. Relative gene expression levels were determined using ΔΔCt method [29]. Expressions were normalized per *ACTB* and *SDHA* genes which were selected as the most stable by Bio-rad CFX Maestro 2.2.

Used primers:

ACTB (F: GCACCACACCTTCTACAAT; R: GTCTCAAACATGATCTGGGT);
AR-FL (F: TTCGCCCCTGATCTGGTTTT; R: TGCCTCATTCGGACACACTG);
KLK3 (F: CCACACCCGCTCTACGATATG; R: GGAGGTCCACACACTGAAGTT);
SDHA (F: TACAAGGTGCGGATTGATGA; R: GTTTTGTCGATCACGGGTCT).

### 4.8. Molecular docking

Molecular docking was performed with the crystal structure of AR-antagonist model. The 3D structures of all compounds were obtained and their energy was minimized by molecular mechanics with Avogadro 1.90.0, a software used for the drawing and characterization of chemical structures. Polar hydrogens were added to ligands and proteins with the AutoDock Tools program [30] and docking studies were performed using AutoDock Vina 1.05 [31]. Interaction between ligand and amino acid residues were modelled in PLIP software [32]. Figures were generated in Pymol ver. 2.0.4 (Schrödinger, LLC).

### 4.9. Ex vivo tissue culture

This pilot study was approved by the Ethical Committee of the University Hospital Olomouc (Ref. No. 127/14) and all five patients signed an informed consent. Prostate tissue from radical prostatectomy was examined by a pathologist and a small piece of PCa lesion was selected for a short time *ex vivo* culture according to the recently optimized protocol [33]. Briefly, PCa tissue was cut into 300 μm slices on vibratome Leica VT1200S (Leica Biosystems). One slice was fixed in 10% formaldehyde at time 0 (T0) as a control. Two slices for each treatment were put on 70 μm pores strainer (MACS SmartStrainer, Miltenyi Biotec) in a tissue culture plate with 1.5 ml of 10% DMEM medium supplemented with 10% fetal bovine serum, 4 mM glutamine, 100 IU/ml penicillin, 100 μg/ml streptomycin, and 1 mM sodium pyruvate, 5 μg/ml insulin, 0.5 μg/ml hydrocortisone, 0.05 μg/ml EGF and enriched with tested compounds or DMSO as vehicle. *Ex vivo* cultures were incubated in standard conditions (37 °C, 5% CO_2_) on a 3D Mini-Shaker (BioSan) that provided continuous mixing for 72 h. The slices were then fixed in 10% formalin and FFPE blocks were prepared. Standard hematoxylin-eosin staining and immunohistochemistry for AR (antibody clone AR441, Dako, dilution 1:25), and Ki-67 (antibody clone MIB1, Dako, dilution 1:200) were performed. Protein expression was assessed semiquantitatively by a pathologist using the histoscore method where the percentage of positive cells (0–100%) was multiplied by staining intensity (0–3), which resulted in a final histoscore between 0 and 300. Ki67 staining at the end-point confirmed the proliferation of cancer cells in the slice.

### 4.10. AR-LBD preparation and micro-scale thermophoresis (MST) measurements

AR-LBD (with His6-tag) was expressed using recombinant plasmid pET-15b-hAR-663-919, which was a generous gift from Elizabeth Wilson (Addgene plasmid # 89083) and expression bacteria BL21(DE3) pLysS similar to the original protocol [34]. Cells were grown in LB medium, the expression was induced by 0.1 mM isopropyl thiogalactopyranoside and suspension was further cultivated overnight at 18 °C. Cells were resuspended in lysis buffer (25 mM Tris, 300 mM NaCl, pH 8.0, 5 mM DTT, supplemented with protease inhibitors and 1% Nonidet P-40), lysed using a sonicator and lysate was clarified by centrifugation at 19 000 *g* for 30 min at 4 °C. The purification was performed on Ni2+-metal affinity-Sepharose column (His-Bind, Merck), pre-equilibrated with 25 mM Tris, 300 mM NaCl, pH 8.0, 5 mM DTT and 50 mM imidazole. Column was thoroughly washed with the equilibration buffer and with 100 mM imidazole in the equilibration buffer, subsequently. Elution was performed by 500 mM imidazole in 25 mM Tris, 300 mM NaCl, pH 8.0, 5 mM DTT. The buffer from elution fraction was exchanged to storage buffer (25 mM Tris, 300 mM NaCl, pH 8.0, 5 mM DTT) without imidazole and concentrated up to 0.5 mg/ml using centrifugal filter unit with 10 kDa cutoff (Merck). MST method was used to prove binding of **3d** in the AR-LBD, which was labelled with the His-Tag Labeling Kit RED-tris-NTA (NanoTemper) (100 nM dye + 800 nM His-tagged protein) for 30 min. The labelled protein was used for MST measurements with or without **3d** in final concentration of 400 nM His-tagged protein in the storage buffer, supplemented with 0.1% Tween. Measurements were done on a Monolith NT.115 instrument (NanoTemper Technologies).

## Acknowledgement

We would like to thank Jakub Bělíček and Veronika Vojáčková for their technical assistance. The authors gratefully acknowledge financial support from the Palacký University Olomouc (IGA_PrF_2022_007 and LF_2022_004), from the Czech Ministry of Education, Youth and Sports via the project National Institute for Cancer Research (Programme EXCELES, ID Project No. LX22NPO5102 funded by the European Union – Next Generation EU), and from the Czech Ministry of Health (grants NU20-03-00201 and DRO (FNOL, 00098892)).

## Appendix A. Supplementary data

Supplementary data related to this article can be found at http://

