## Supporting Information for "A-ring-fused pyrazoles of dihydrotestosterone targeting prostate cancer cells via the downregulation of the androgen receptor"

### Table of contents

**Scheme S1** Synthesis of 2-(hetero)arylidene derivatives of DHT **3**

**Figure S1** Partial 2D-NOESY spectrum of compound **8a** **4**

**Figure S2** Partial 2D-NOESY spectrum of compound **9a** **4**

$^{1}$H and $^{13}$C NMR spectra of the synthesized compounds **5-31**

**Table S1** AR transcriptional activity in antagonist and agonist mode **32**

**Figure S3** Western blot of AR-regulated proteins in LAPC-4 upon treatment with compounds **34**

**Figure S4** Colony formation assay of LAPC-4 treated by series **1**, **2**, **8i-8k**, **10i-10k**, standards **35**

**Figure S5** Colony formation assay of LAPC-4 treated by series **3, 4, 8a-8h, 10a-10h** **36**

**Figure S6** Dose dependent effect of **3d** on AR-signalling in 22Rv1 and LNCaP **37**

**Figure S7** Antiproliferative activity of **3d** in LAPC-4 (long time treatment, protein levels) **38**

**Figure S8** MST measurement of binding of **3d** in AR-LBD **39**

**Figure S9** Antiproliferative activity of **3d** on cell lines' panel **40**

**Figure S10** Cell cytometry of 22Rv1 cells after 48h treatment with **3d** or bavdegalutamide **41**

**Figure S11** Relative normalized expression of AR in LAPC-4 cells **42**

**Figure S12** Images of Ki67 and AR immunostaining of tumor slices after 72 h *ex vivo* treatment **43**

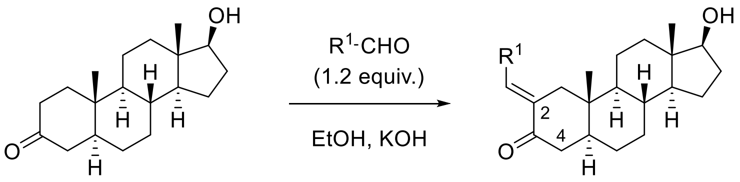


DHT 5

$R^1$: Ph (**a**); *p*-$CH_3$-$C_6H_4$ (**b**); *p*-MeO-$C_6H_4$ (**c**); *p*-F-$C_6H_4$ (**d**); *p*-Cl-$C_6H_4$ (**e**); *p*-Br-$C_6H_4$ (**f**); furan-2-yl (**g**); tiophen-2-yl (**h**); *p*-OH-$C_6H_4$ (**i**); $CH_3$ (**j**)

**Scheme S1.** Synthesis of 2-(hetero)arylidene derivatives of DHT. *Reagents and conditions: ref.* [18] *for* ***5a–e***, ***5g*** *and* ***5h****; 0 °C, 3 h for* ***5f*** *(86%); reflux, 16 h (with MOM-protected p-OH-phenol), then dil. HCl, MeOH (36% after two steps) for* ***5i****; ref* [17] *for* ***5j****.*

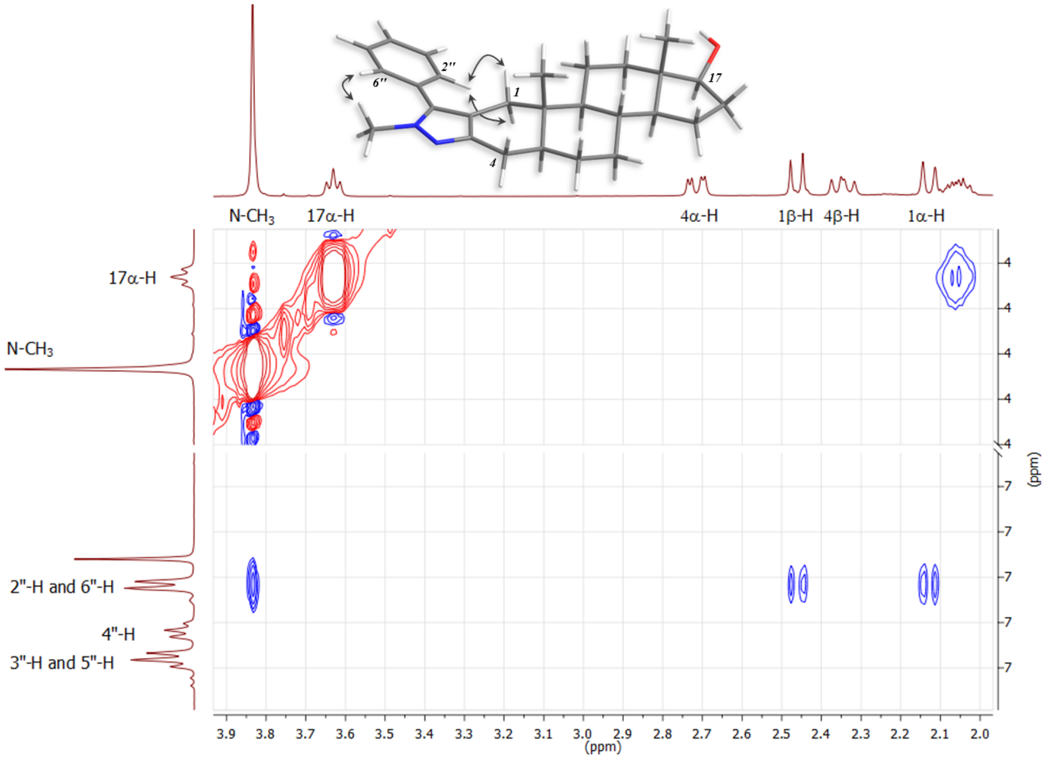


**Figure S1.** Partial 2D-NOESY spectrum of compound **8a** and the correlations between protons observed.

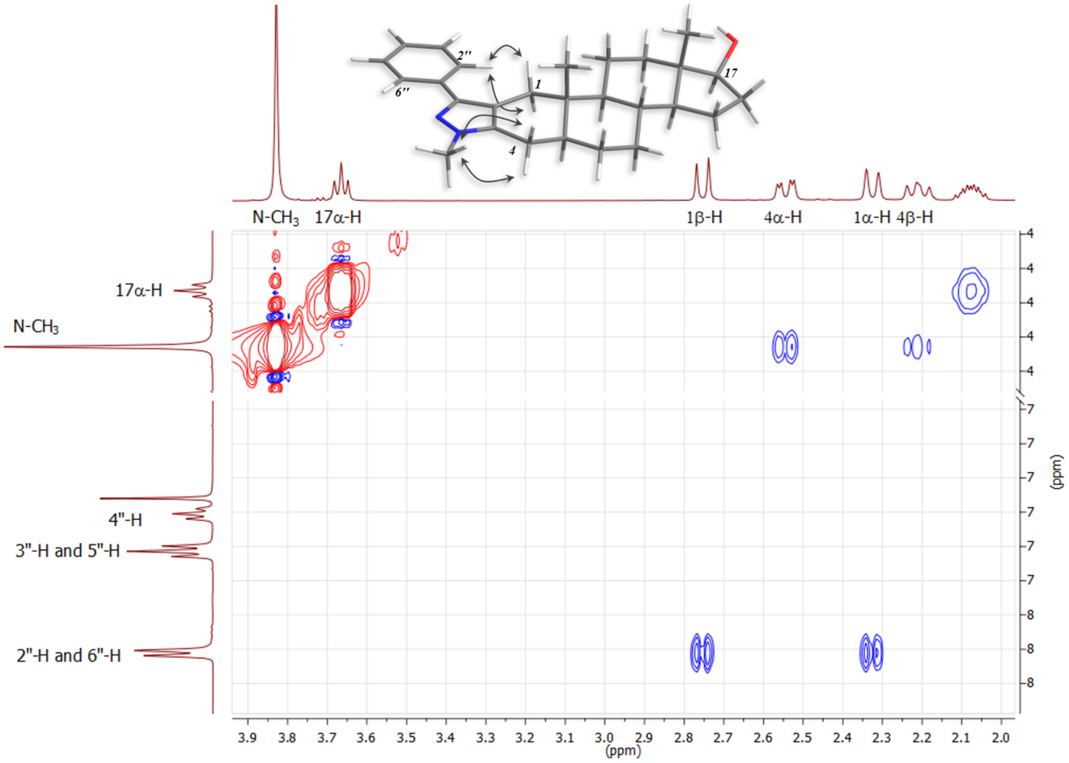


**Figure S2.** Partial 2D-NOESY spectrum of compound **9a** and the correlations between protons observed.

### $^{1}H$ and $^{13}C$ NMR spectra of the synthesized compounds

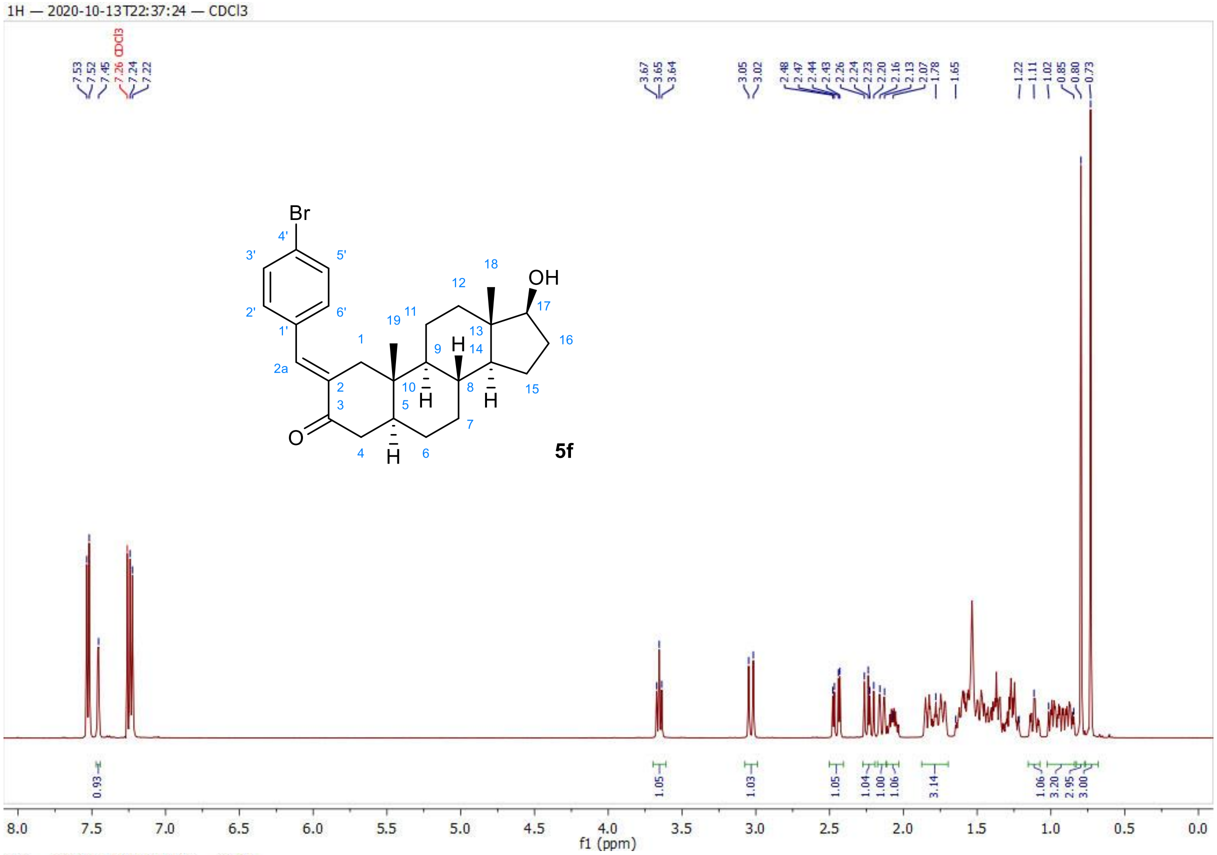


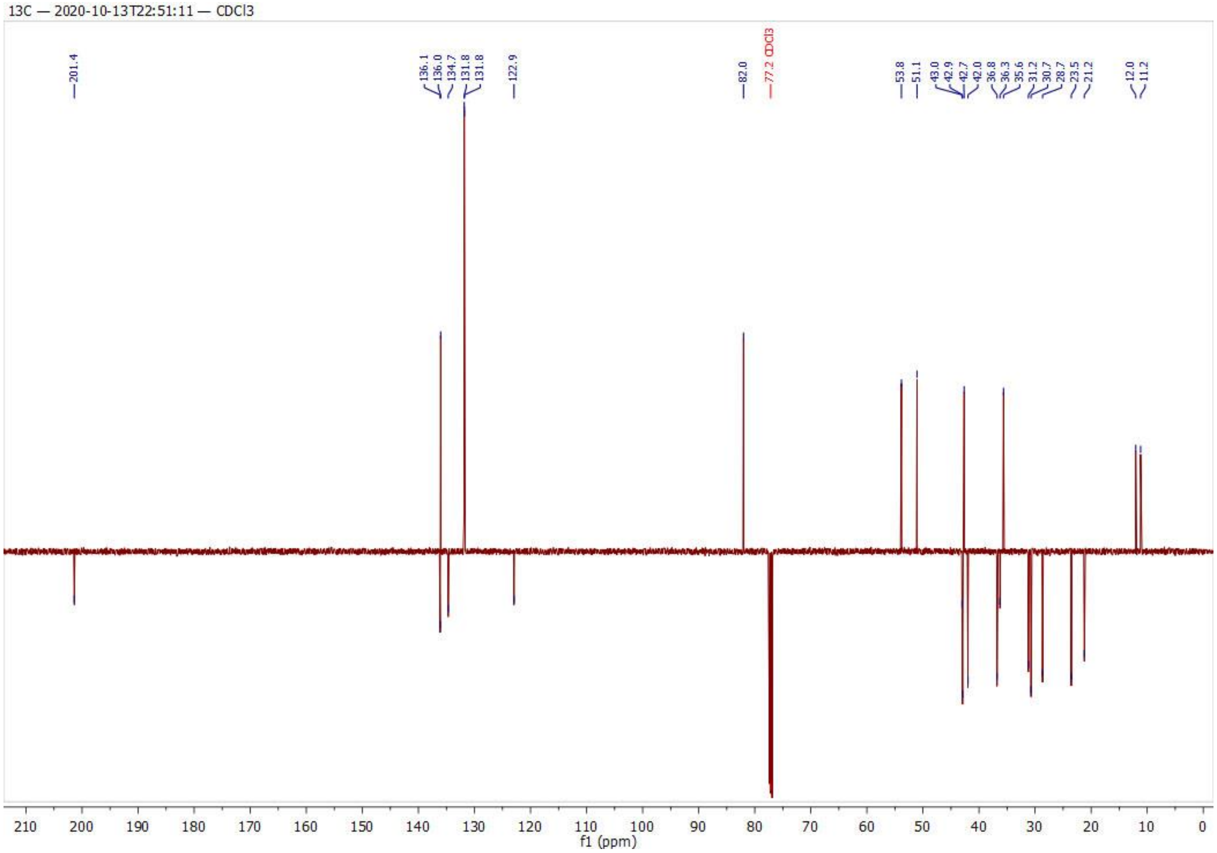

1H — 2021-04-30T22:36:55 — DMSO

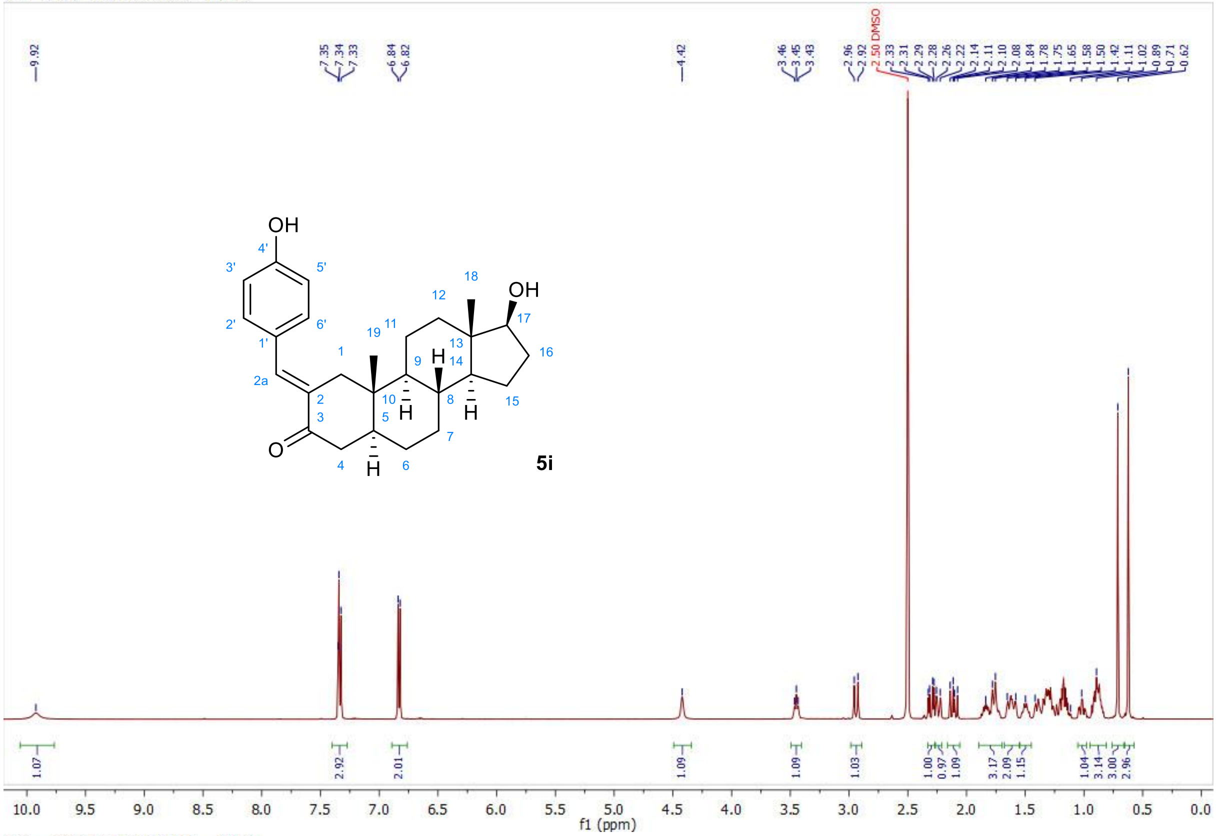


13C — 2021-04-30T22:34:45 — DMSO

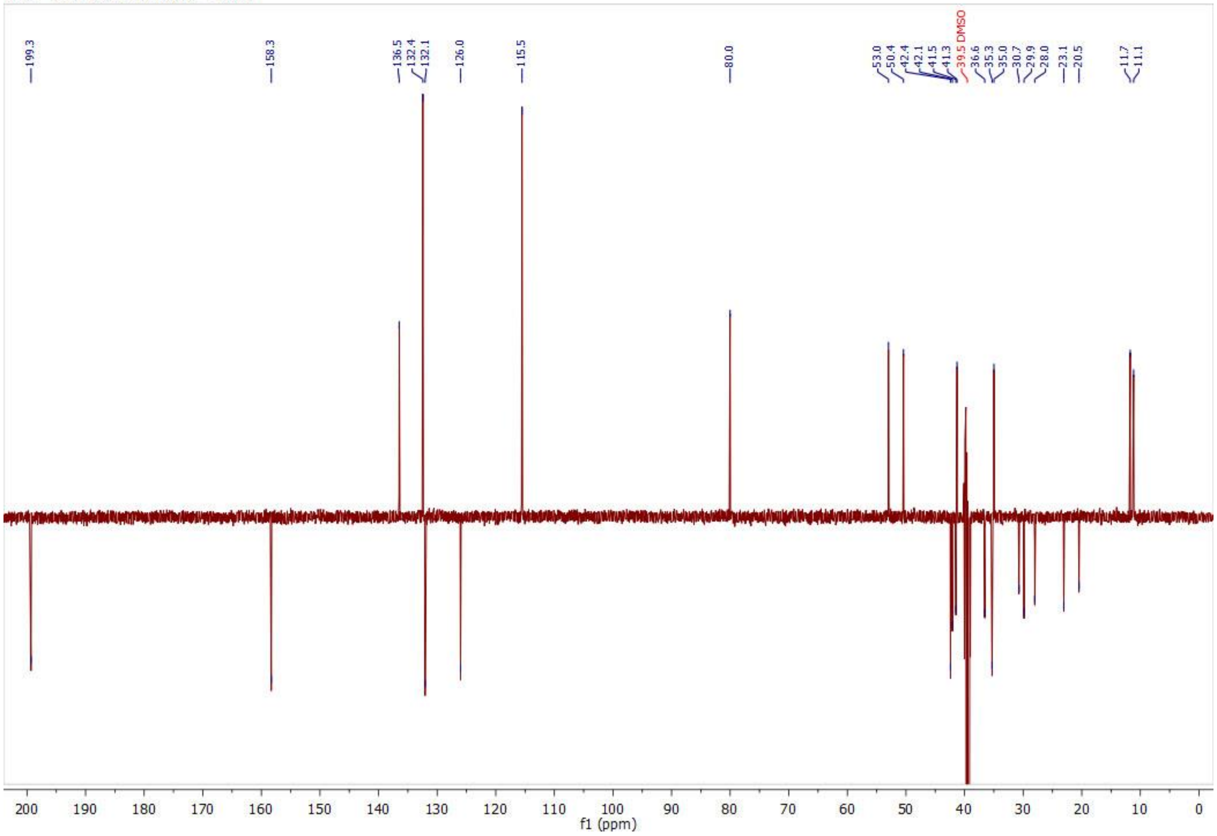

1H — 2020-11-14T13:00:19 — CDCl3

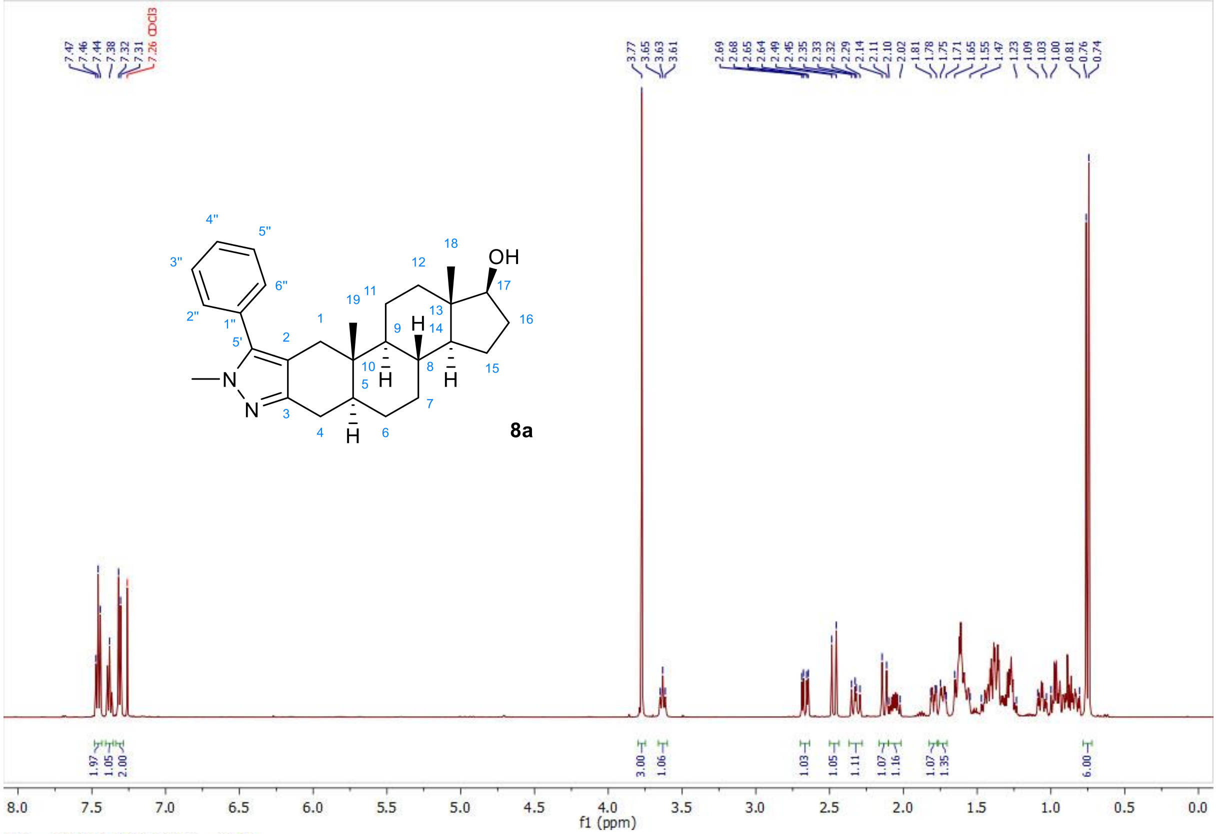


13C — 2020-11-14T13:13:57 — CDCl3

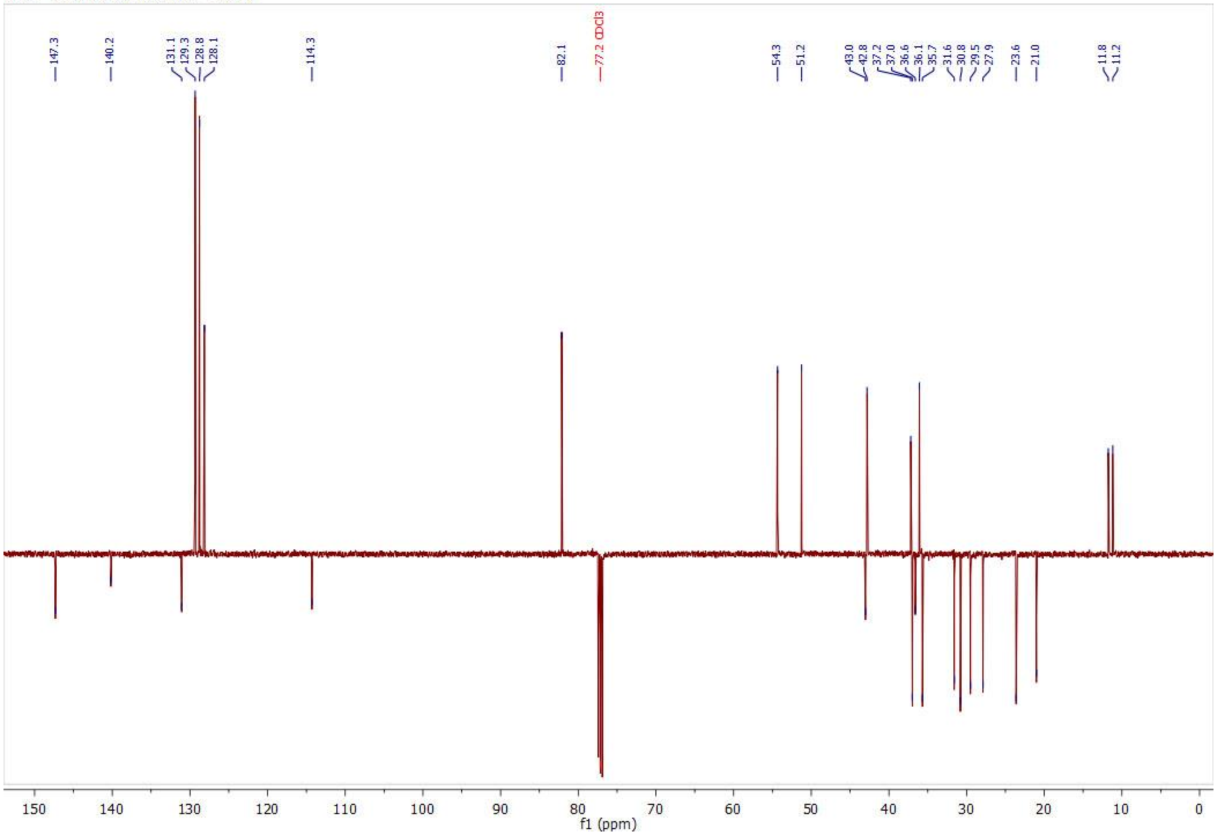

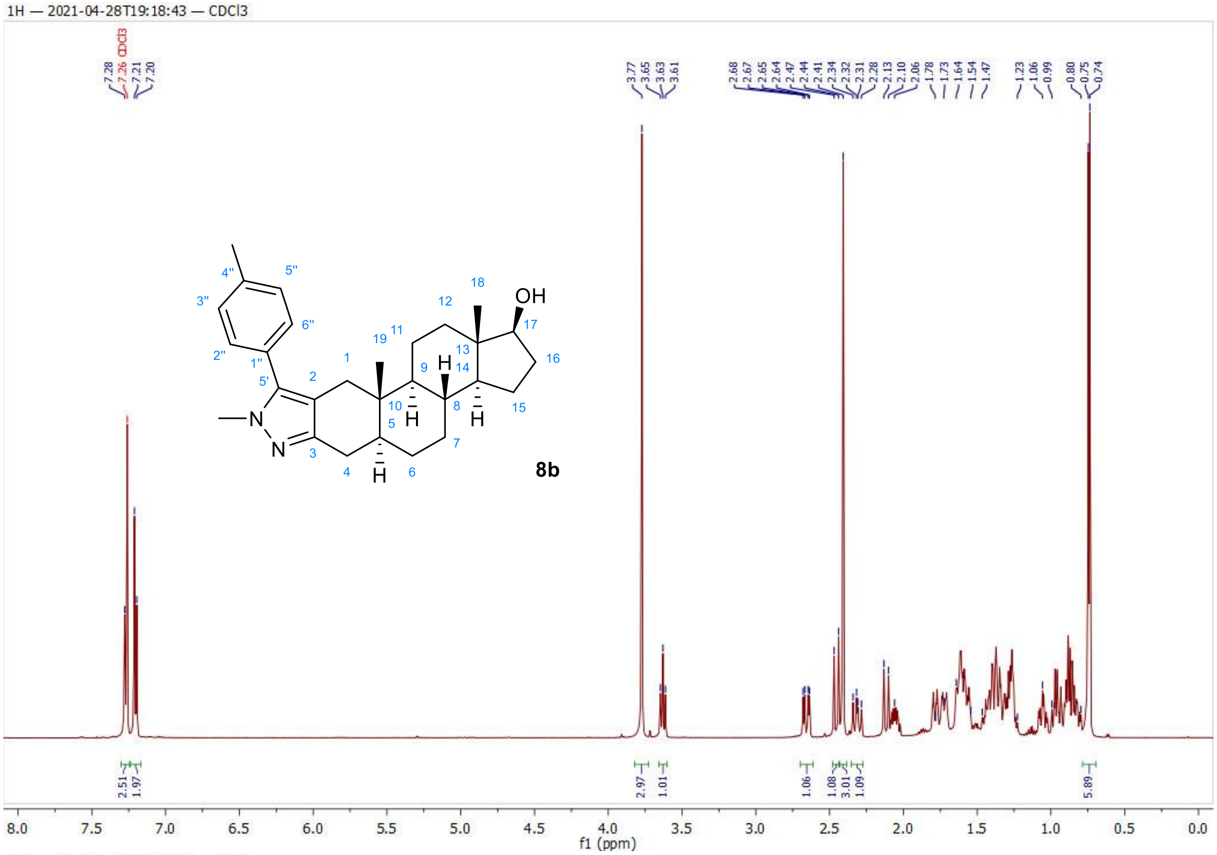
1H — 2021-04-28T19:18:43 — CDCl3
7.28
7.26 CDCl3
7.21
7.20
3.77
3.65
3.63
3.61
2.68
2.67
2.65
2.64
2.47
2.44
2.41
2.34
2.32
2.31
2.28
2.13
2.10
2.06
1.78
1.73
1.64
1.54
1.47
1.23
1.06
0.99
0.80
0.75
0.74
2.51
1.97
2.97
1.01
1.06
1.08
3.01
1.09
5.89
8b
OH
f1 (ppm)

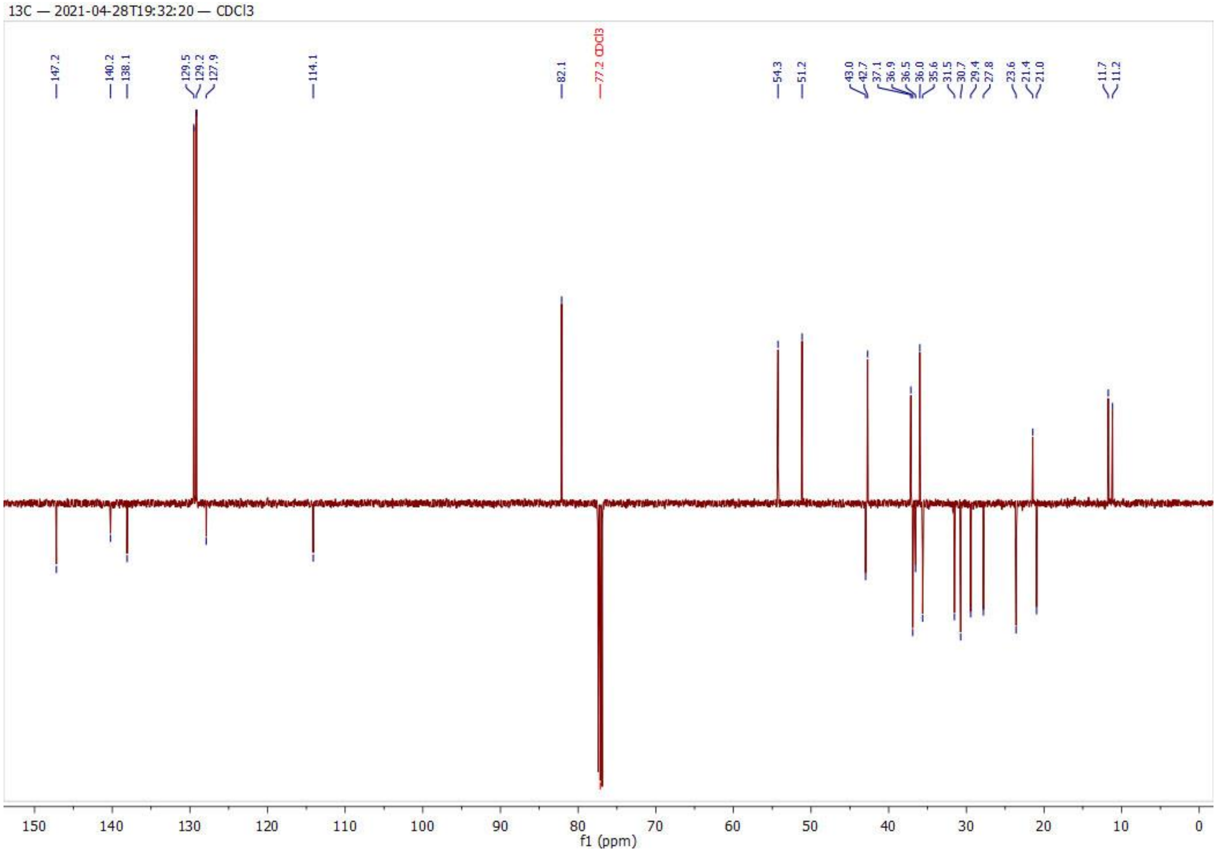
13C — 2021-04-28T19:32:20 — CDCl3
147.2
140.2
138.1
129.5
129.2
127.9
114.1
82.1
77.2 CDCl3
54.3
51.2
43.0
42.7
37.1
36.9
36.5
36.0
35.6
31.5
30.7
29.4
27.8
23.6
21.4
21.0
11.7
11.2
f1 (ppm)

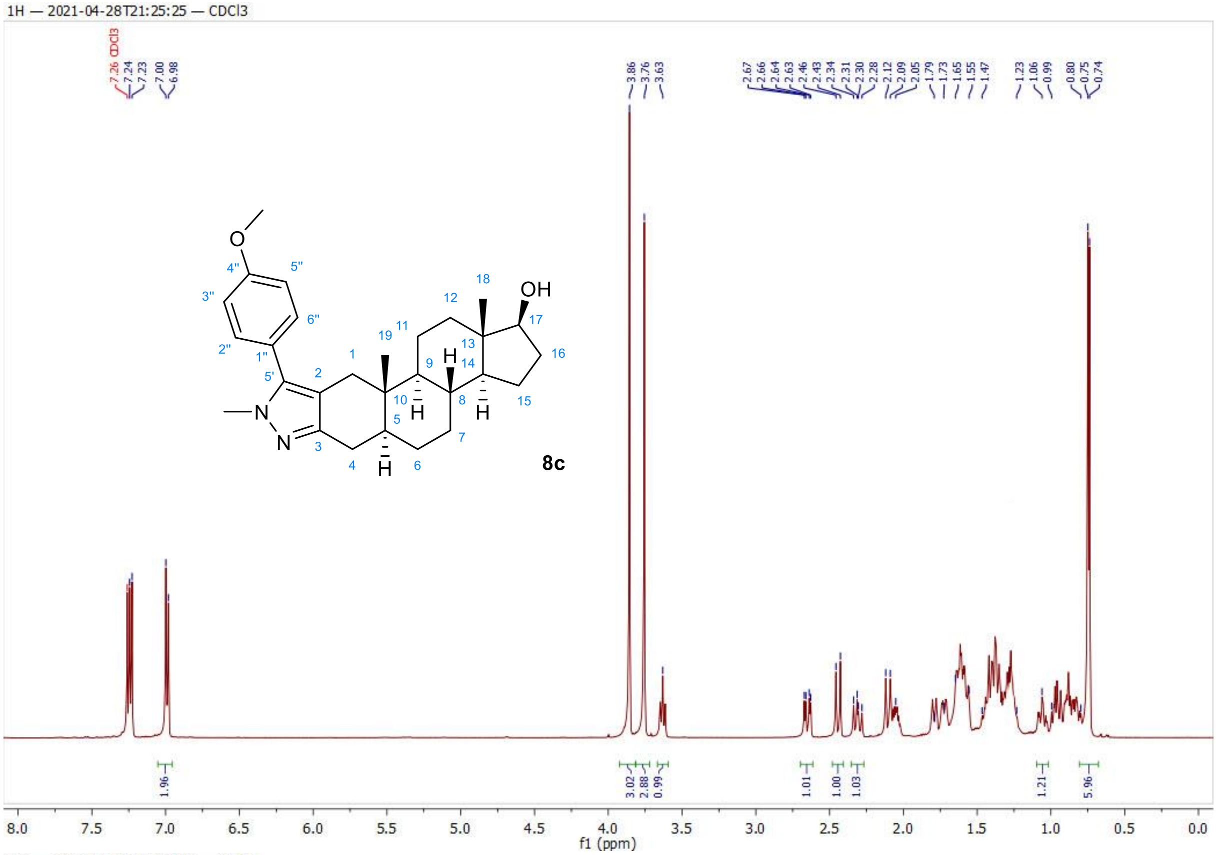
1H — 2021-04-28T21:25:25 — CDCl3
8c
OH
f1 (ppm)

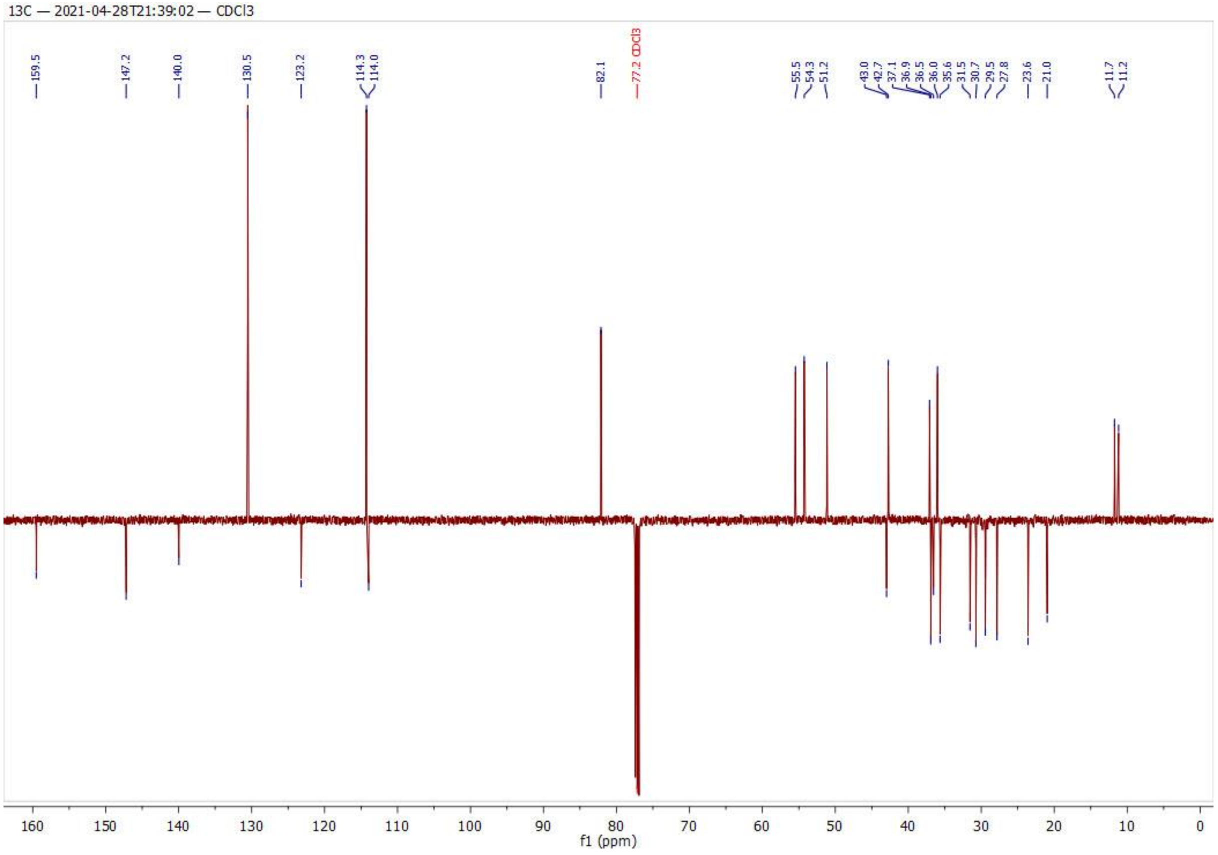
13C — 2021-04-28T21:39:02 — CDCl3
f1 (ppm)

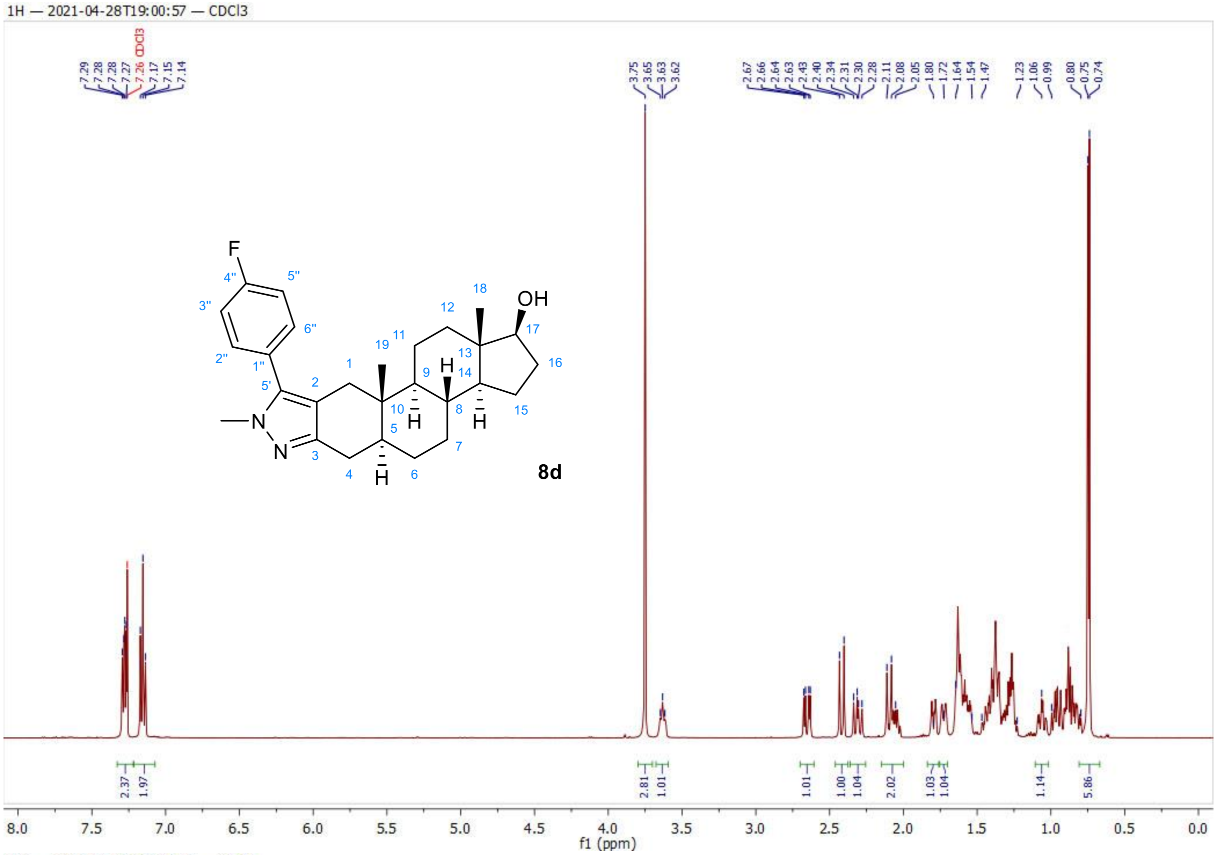
1H — 2021-04-28T19:00:57 — CDCl3
8d
OH
F
f1 (ppm)

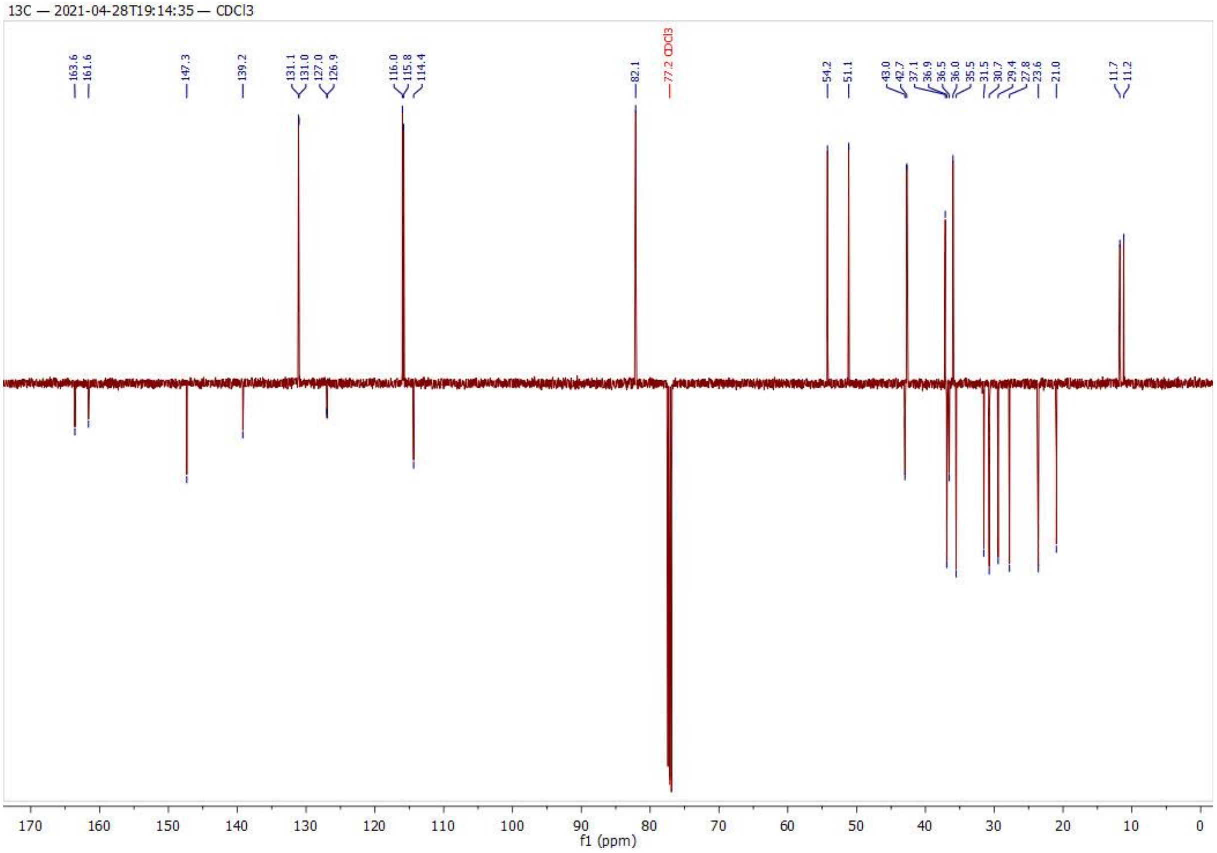
13C — 2021-04-28T19:14:35 — CDCl3
f1 (ppm)

1H — 2020-11-14T13:36:38 — CDCl3

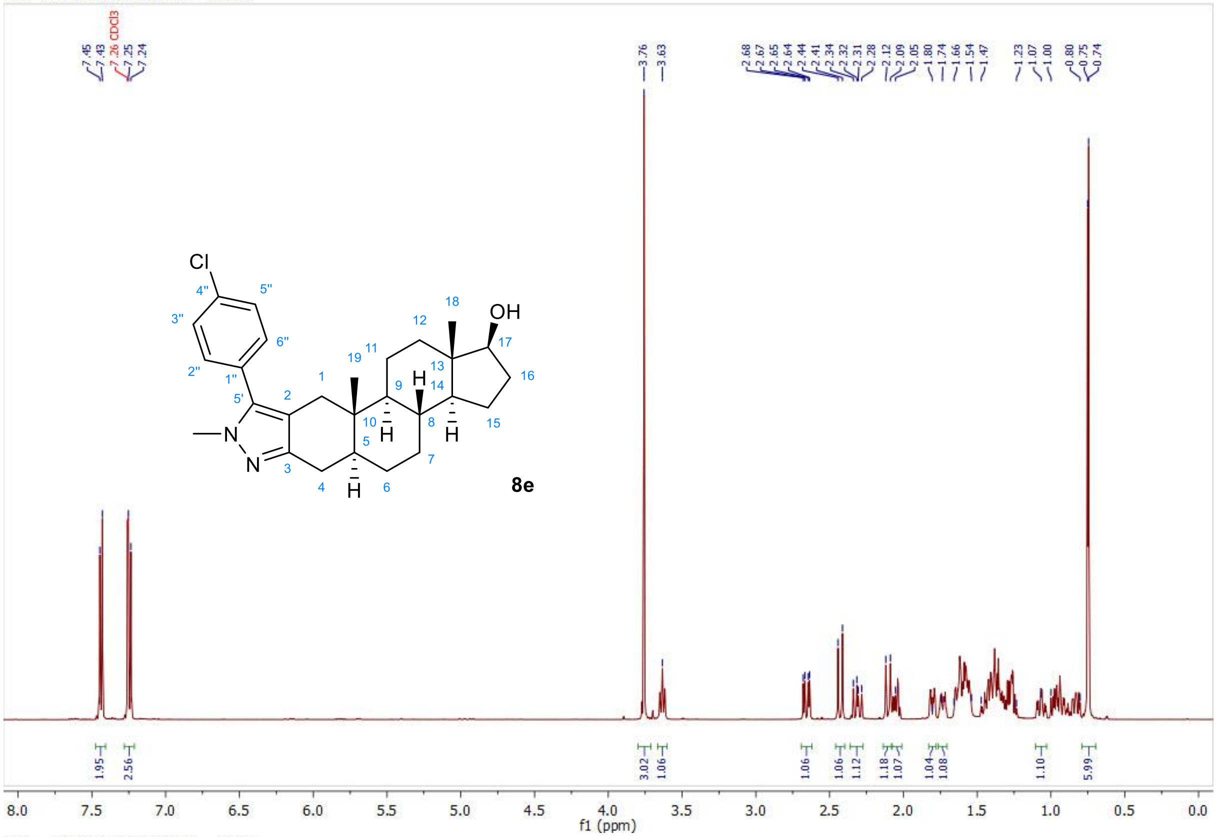


13C — 2020-11-14T13:50:37 — CDCl3

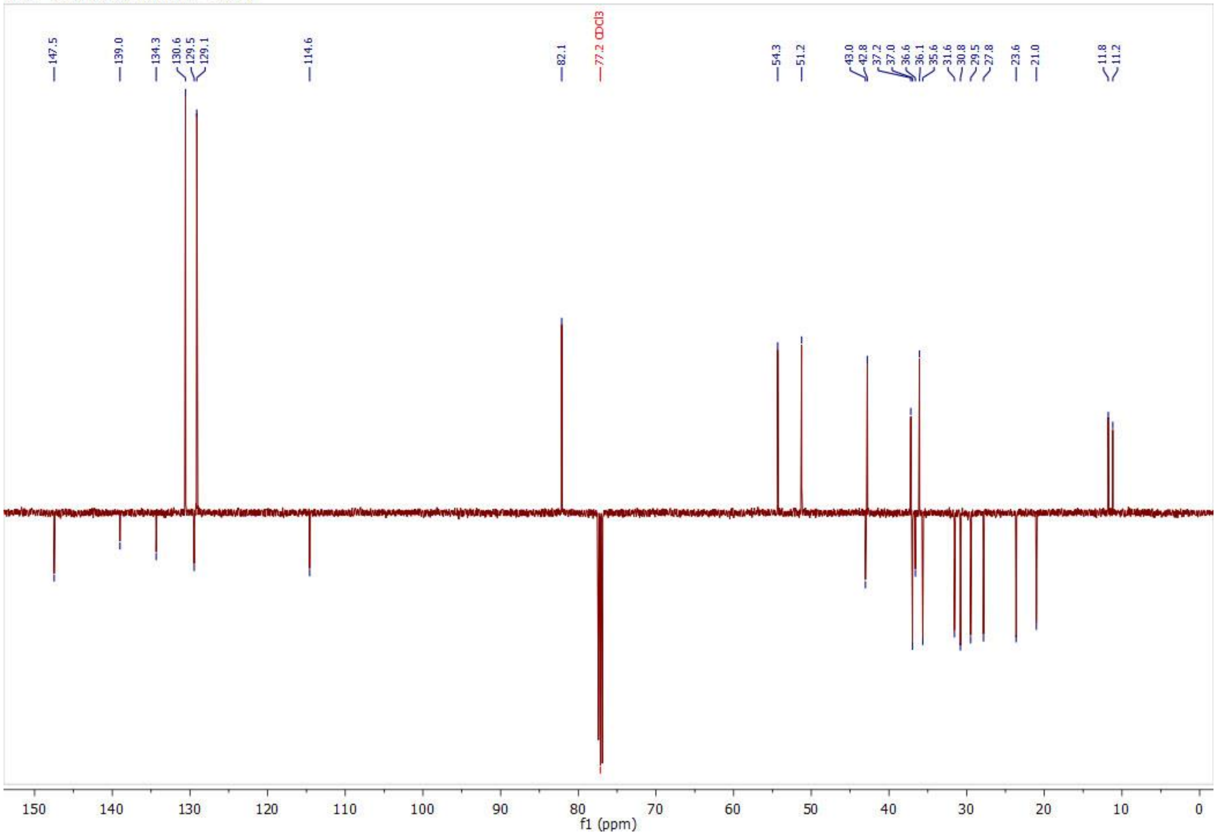

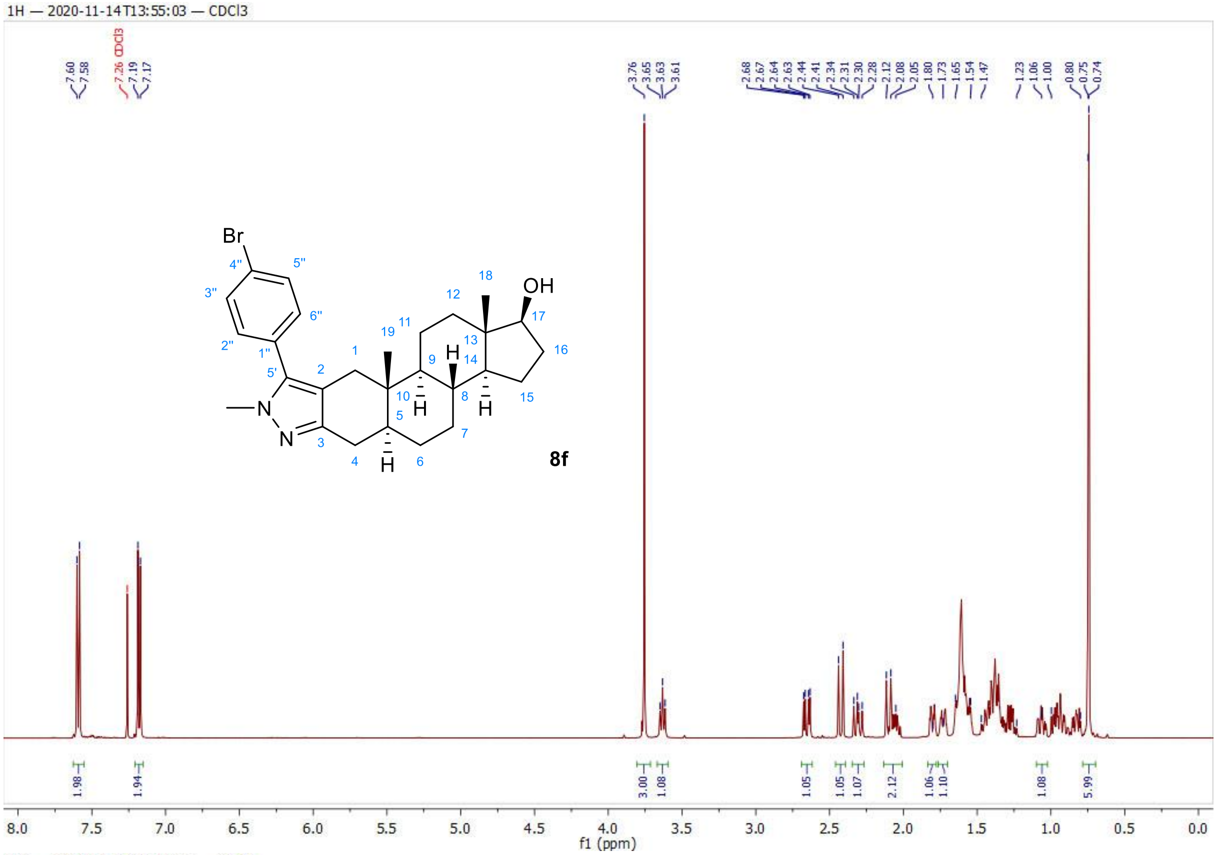
1H — 2020-11-14T13:55:03 — CDCl3
8f
f1 (ppm)

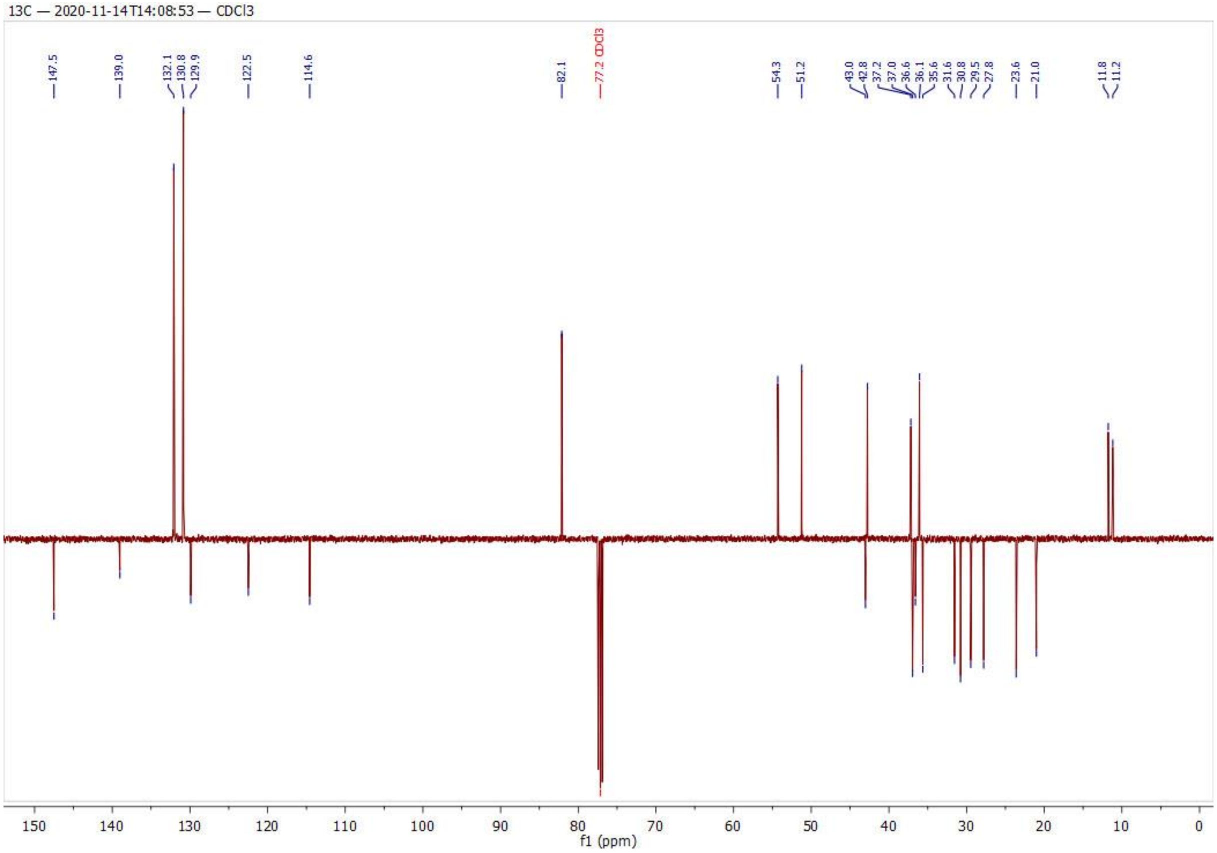
13C — 2020-11-14T14:08:53 — CDCl3
f1 (ppm)

1H — 2021-02-13T18:06:57 — CDCl3

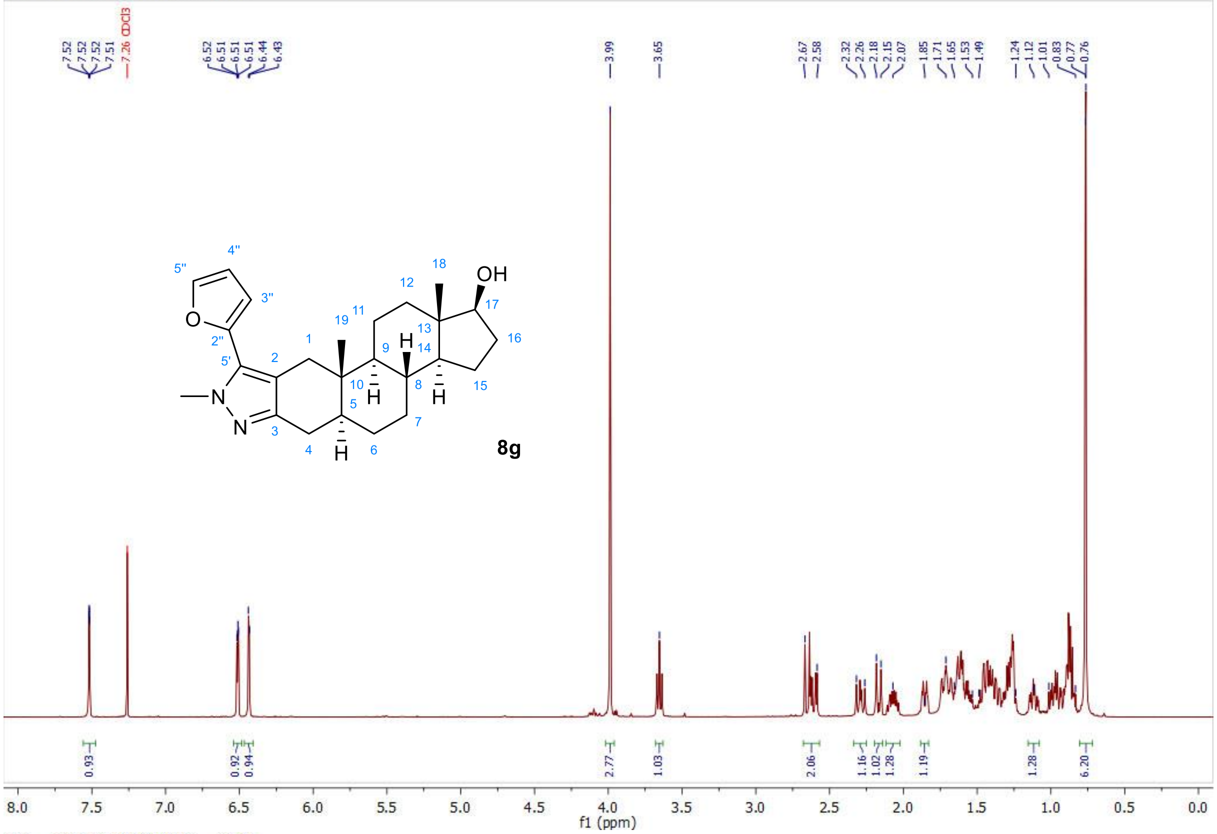


13C — 2021-02-13T18:20:32 — CDCl3

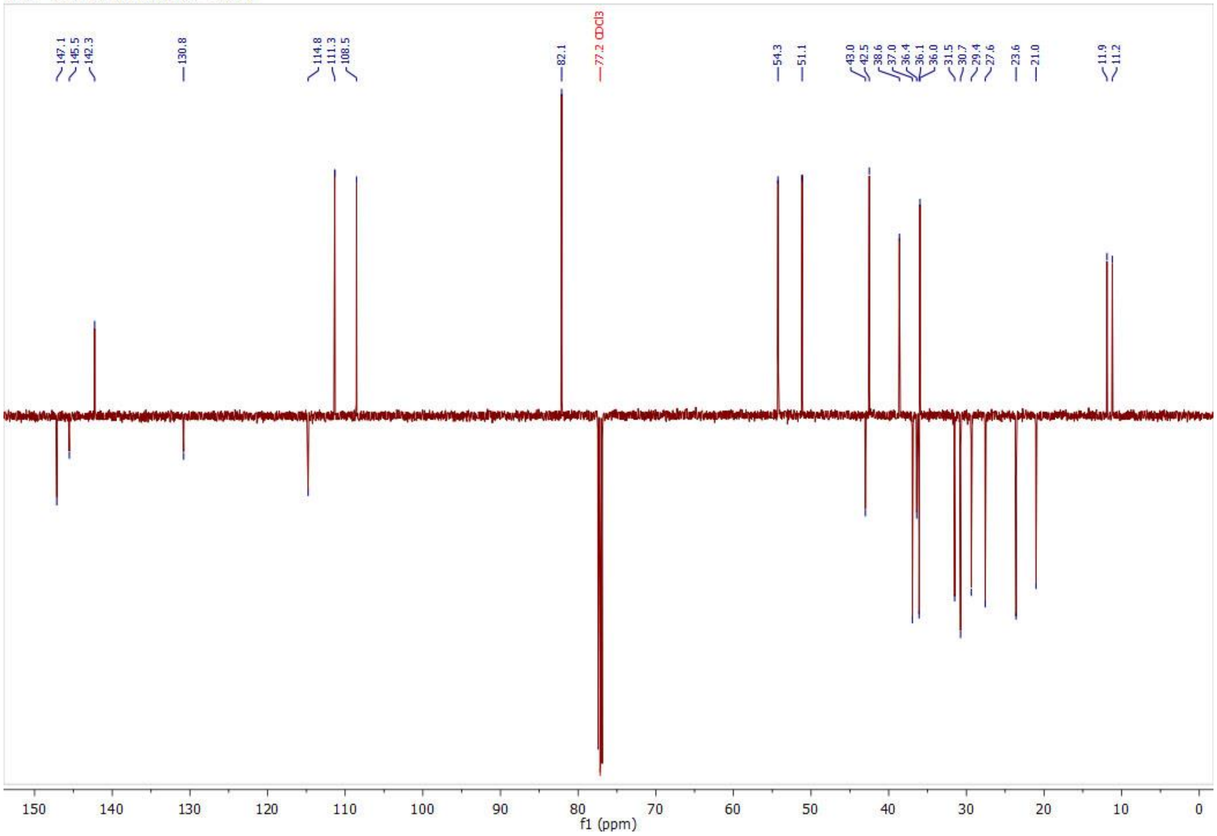

1H — 2021-09-18T00:49:18 — CDCl3

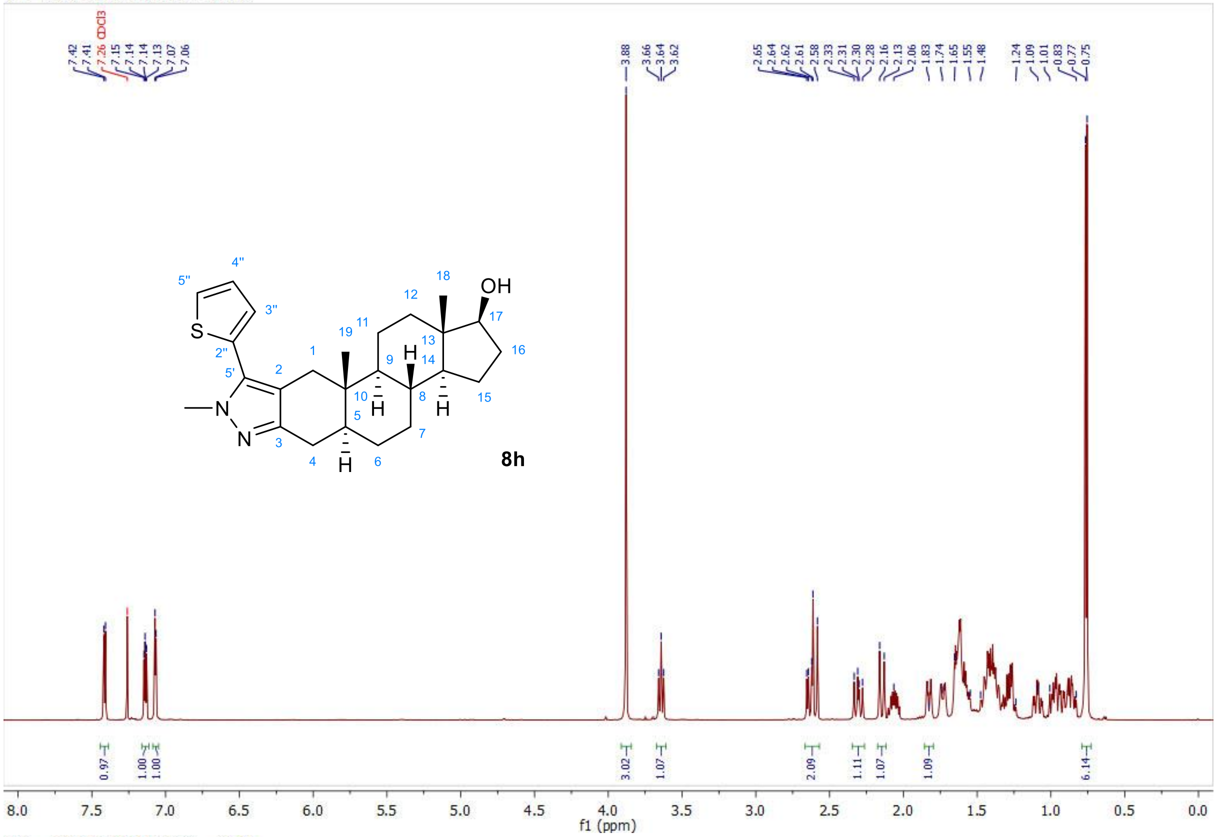


13C — 2021-09-18T01:03:06 — CDCl3

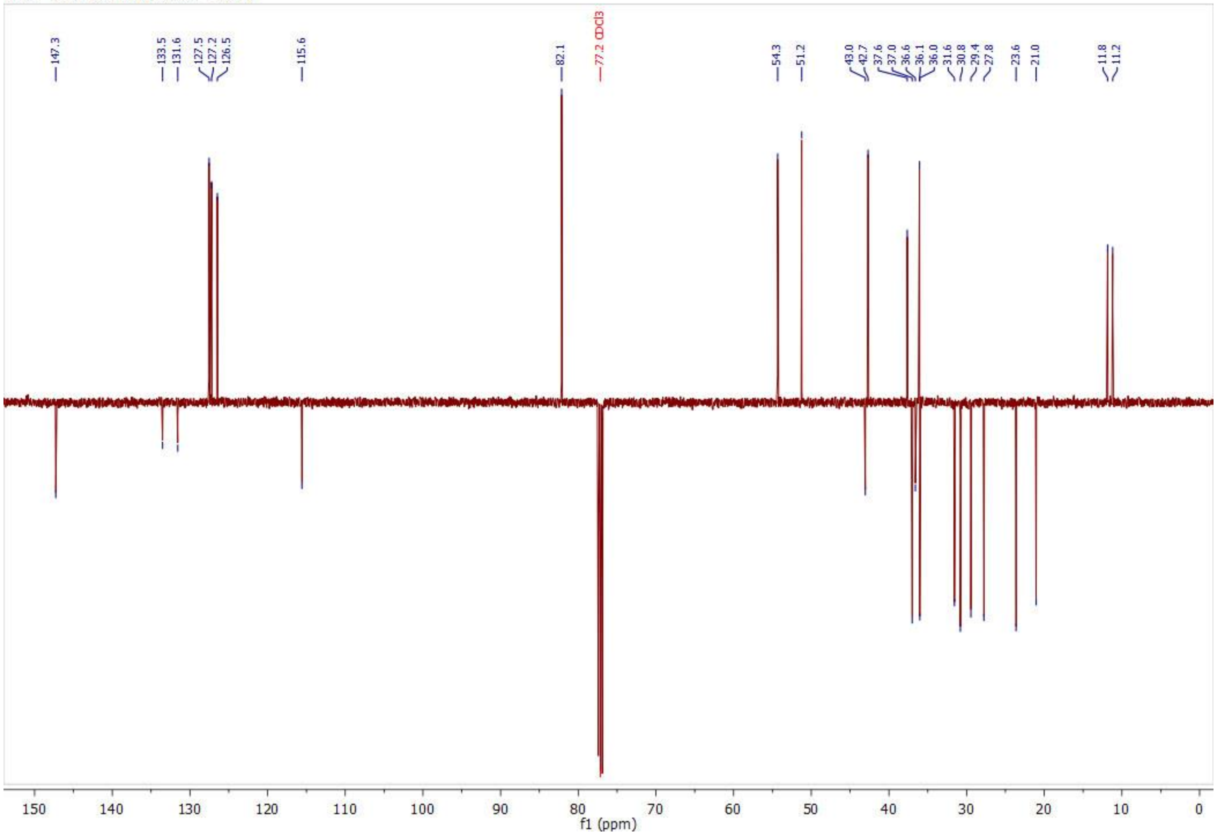

1H — 2021-04-28T21:44:05 — CDCl3

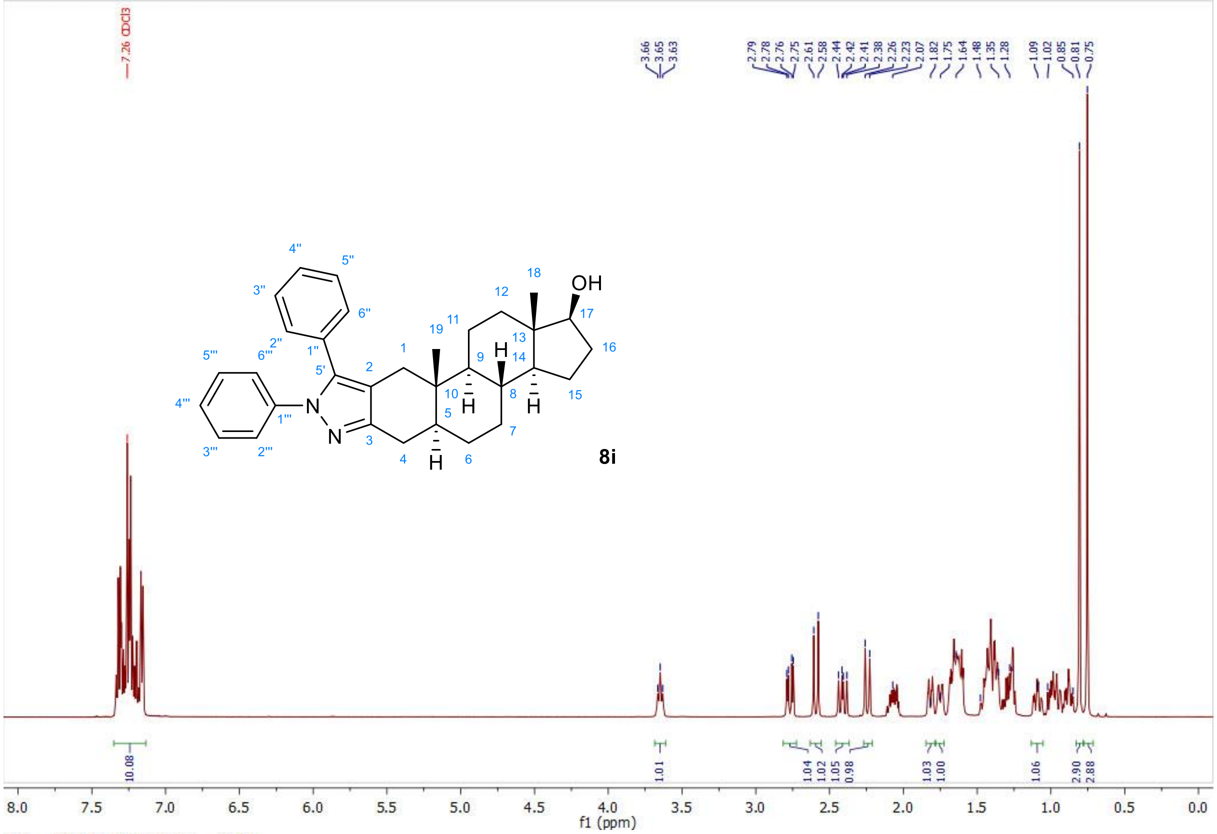


13C — 2021-04-28T21:57:43 — CDCl3

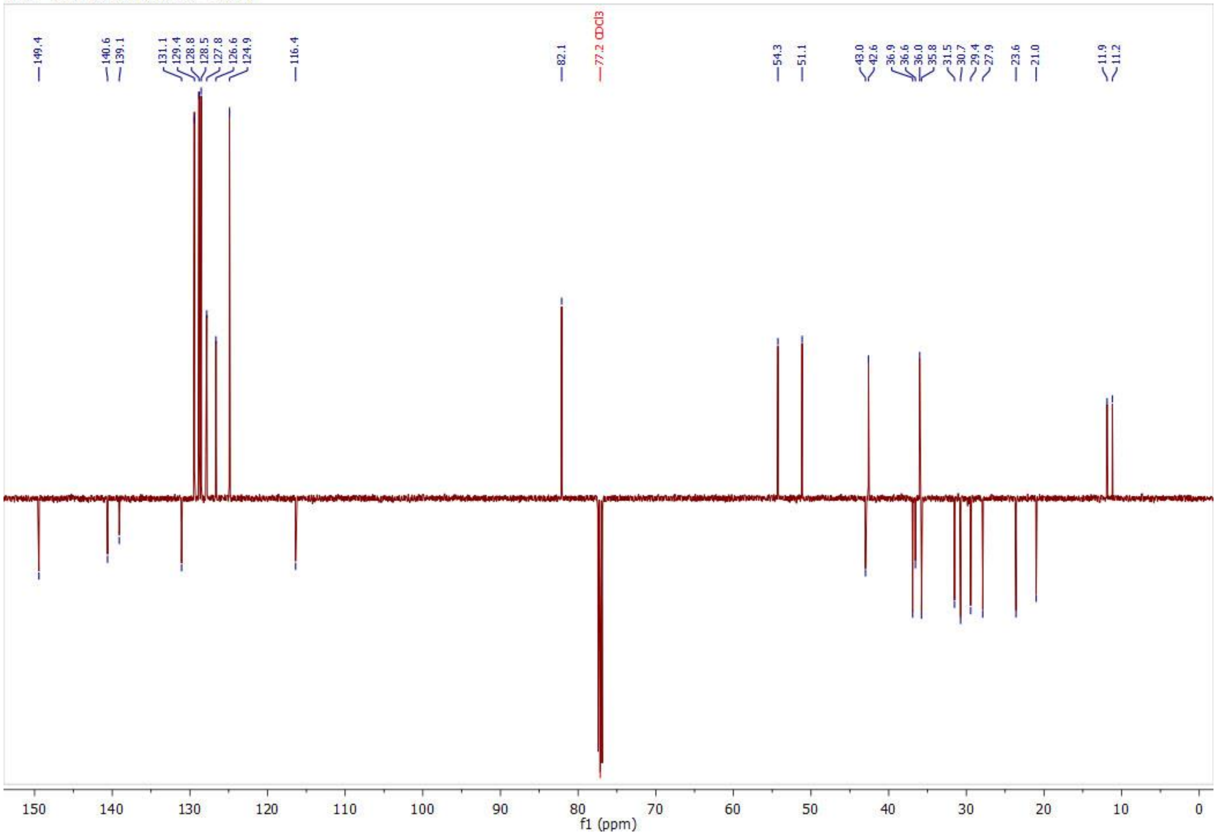

1H — 2021-04-28T22:20:05 — CDCl3

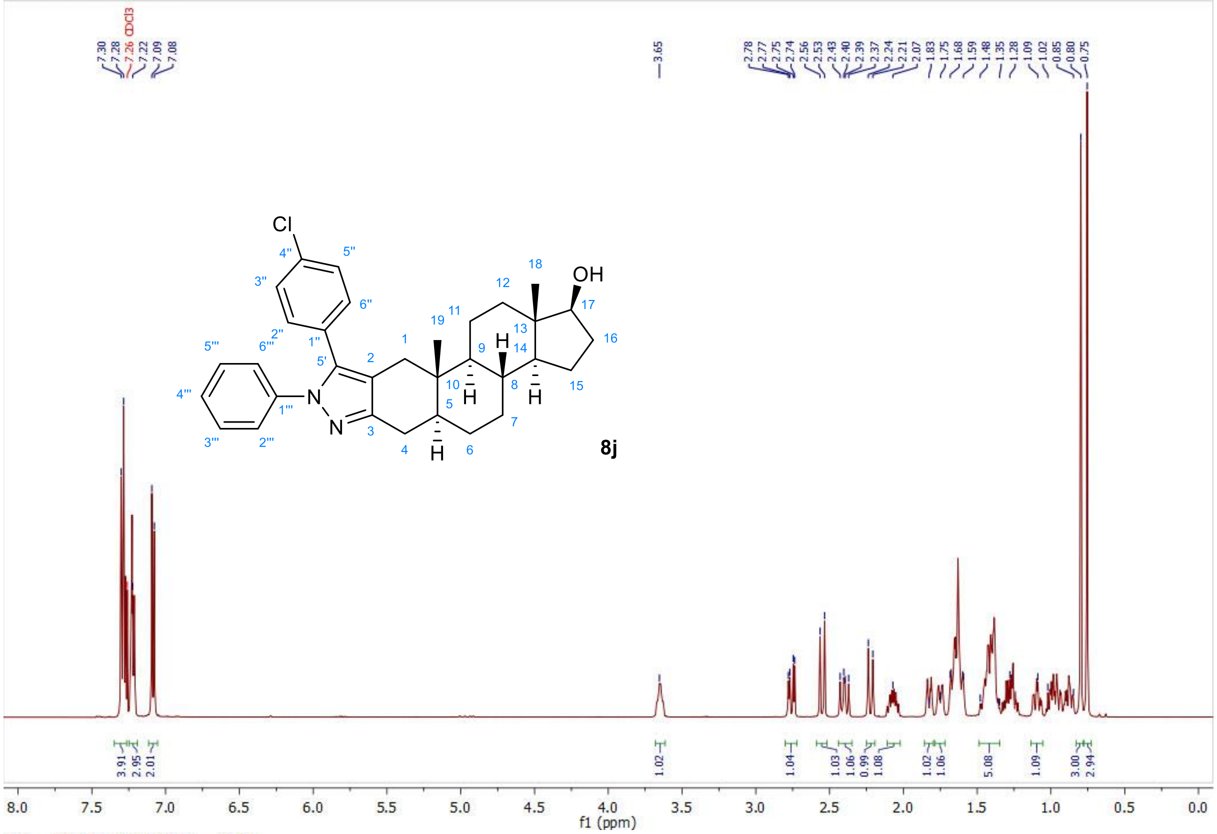


13C — 2021-04-28T22:33:42 — CDCl3

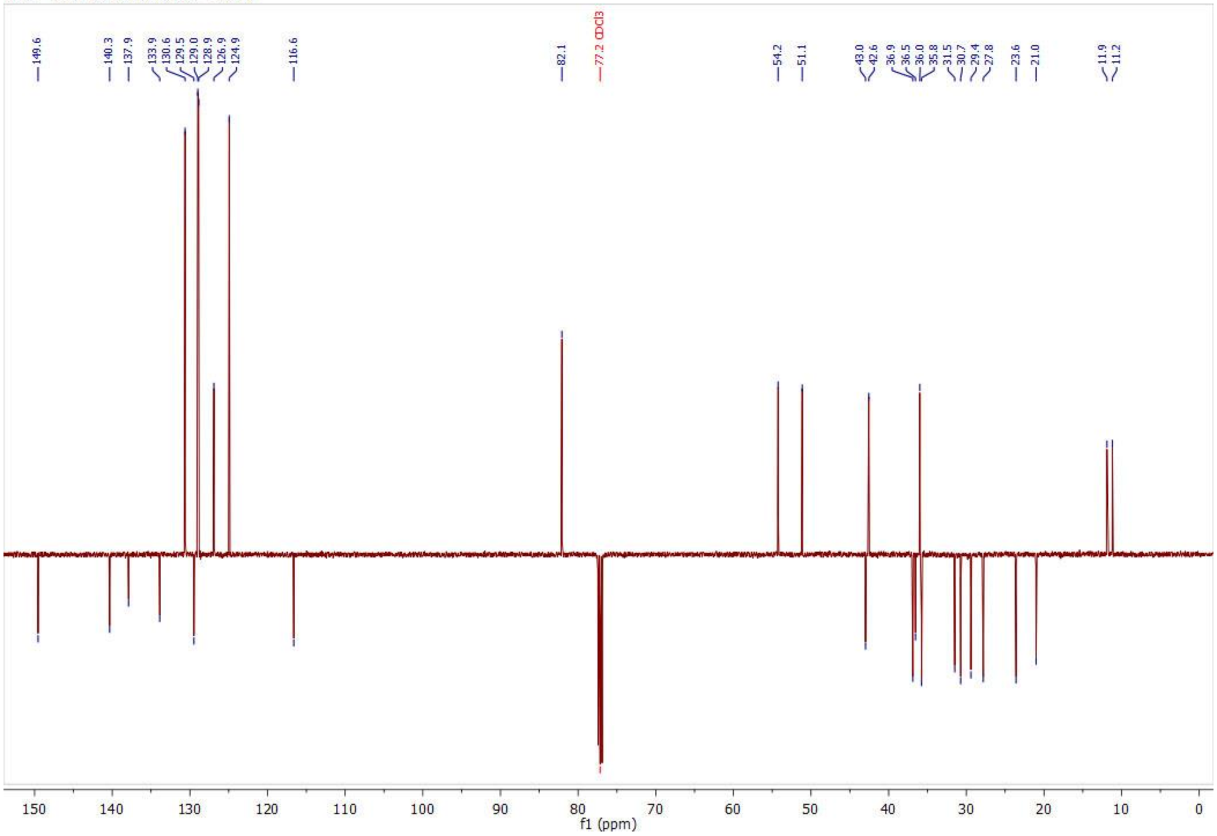

1H — 2021-05-13T21:32:57 — DMSO

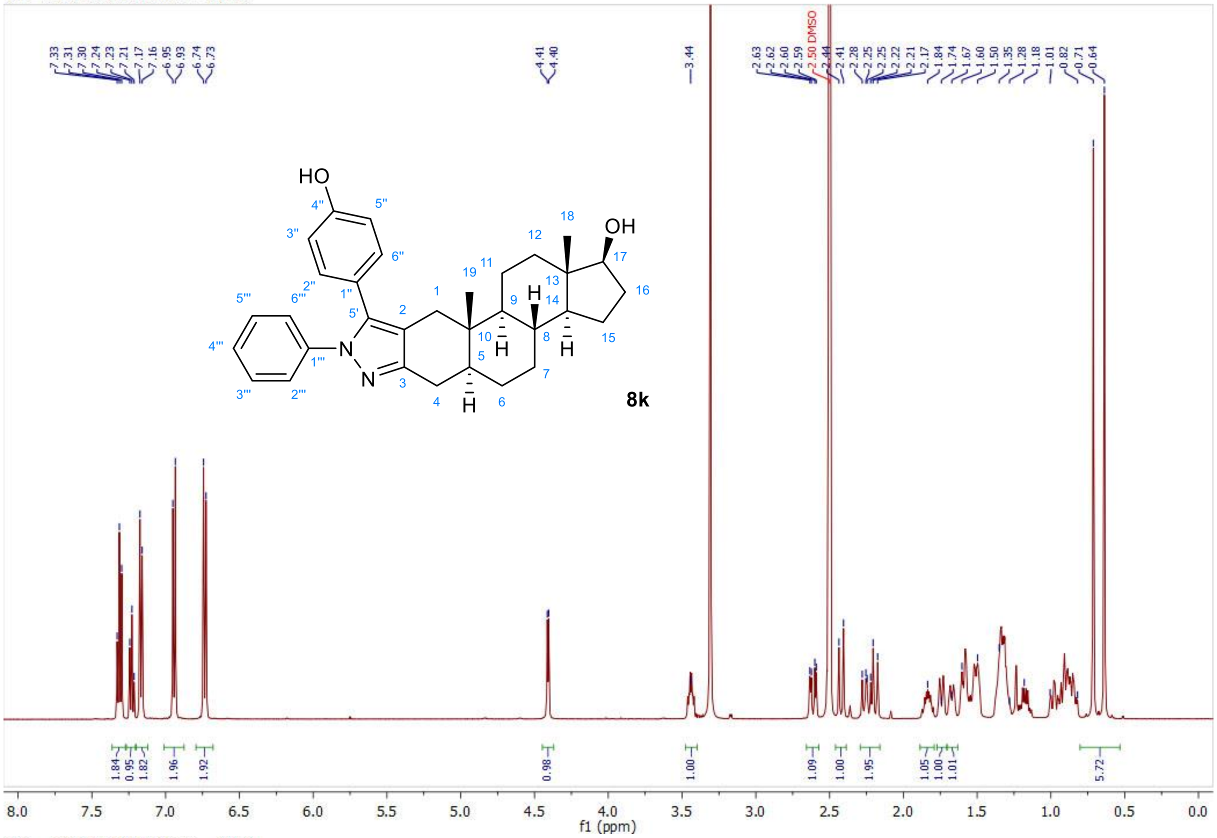


13C — 2021-05-13T21:46:51 — DMSO

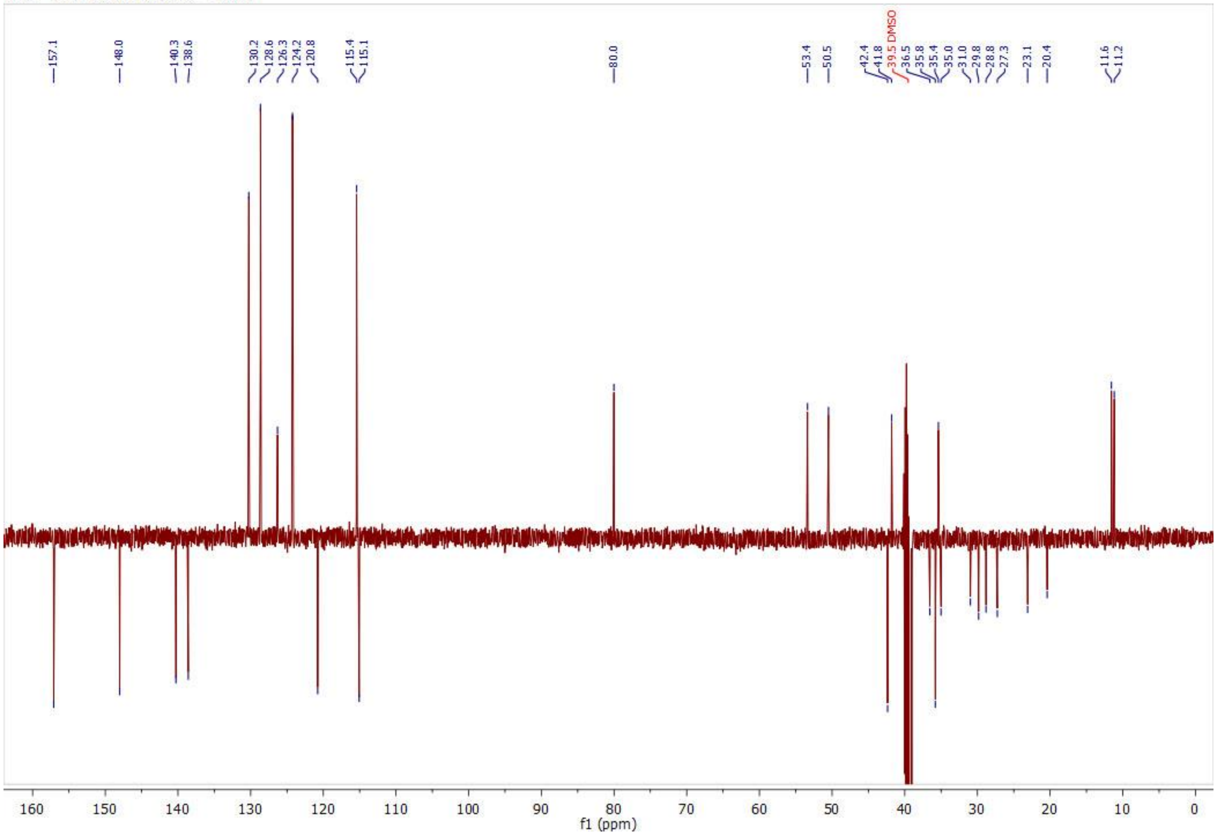

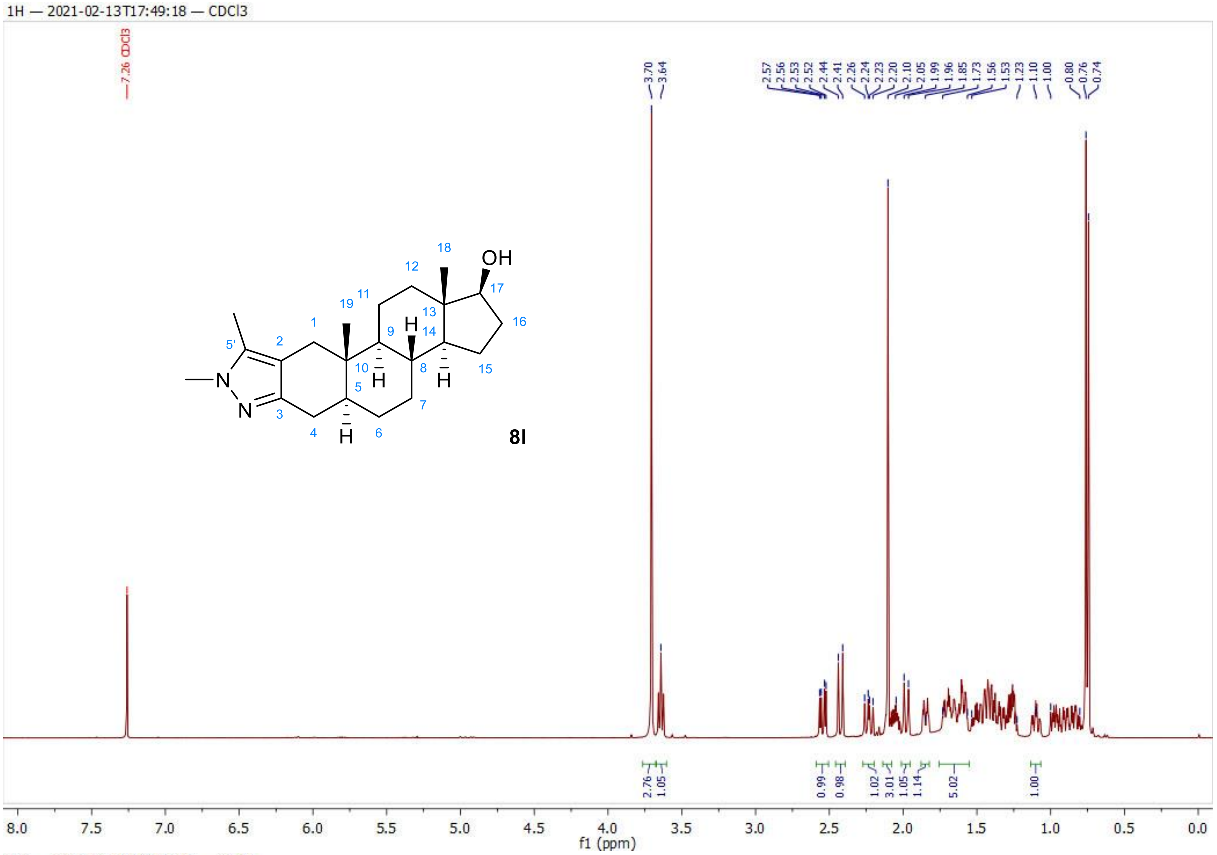
1H — 2021-02-13T17:49:18 — CDCl3
8l
f1 (ppm)

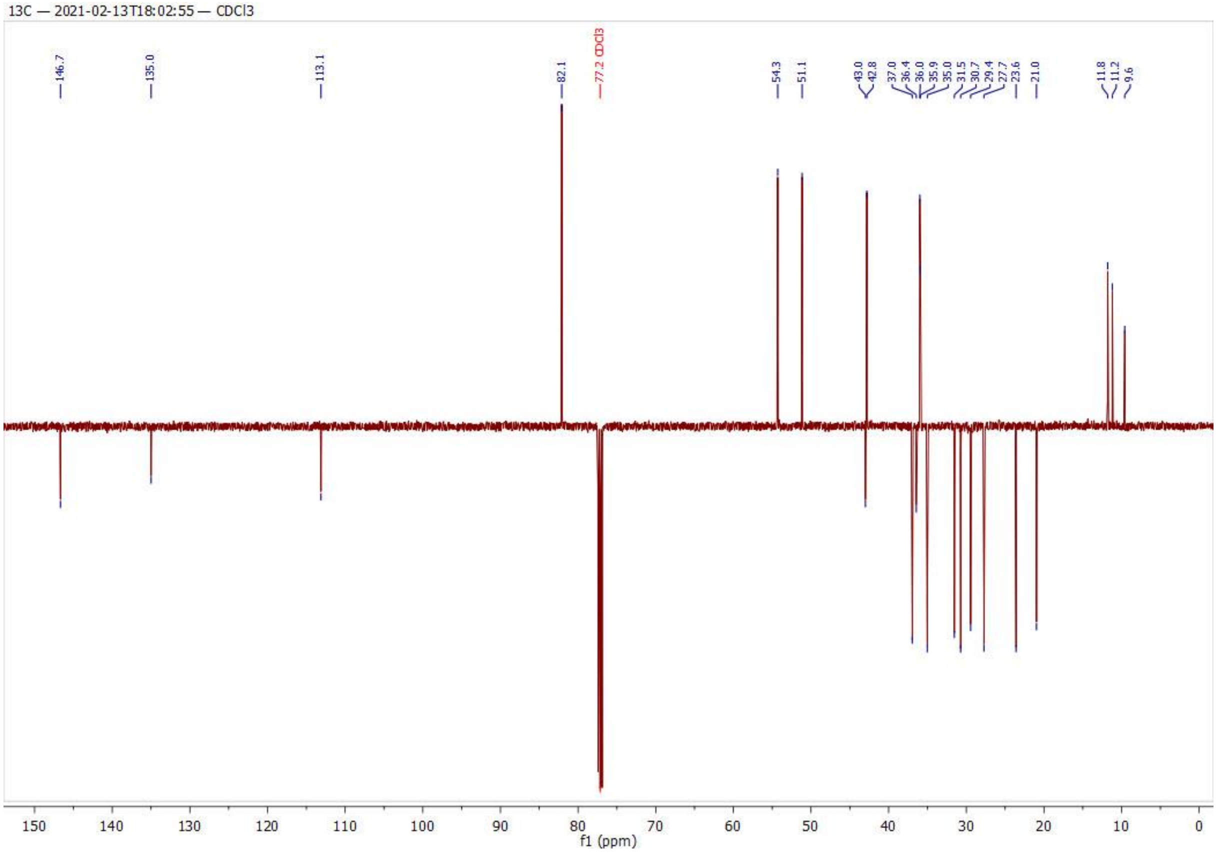
13C — 2021-02-13T18:02:55 — CDCl3
f1 (ppm)

1H — 2017-12-06T11:17:44 — CDCl3

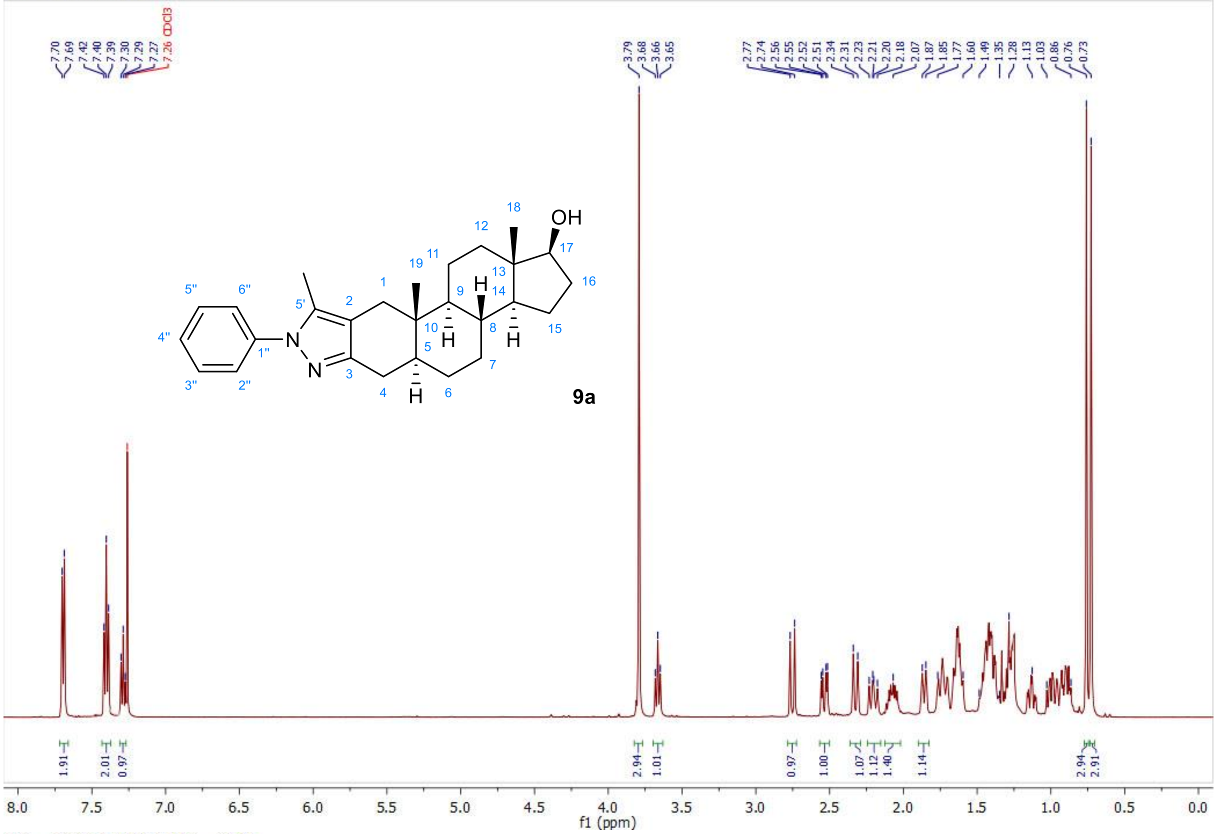


13C — 2017-11-11T13:24:44 — CDCl3

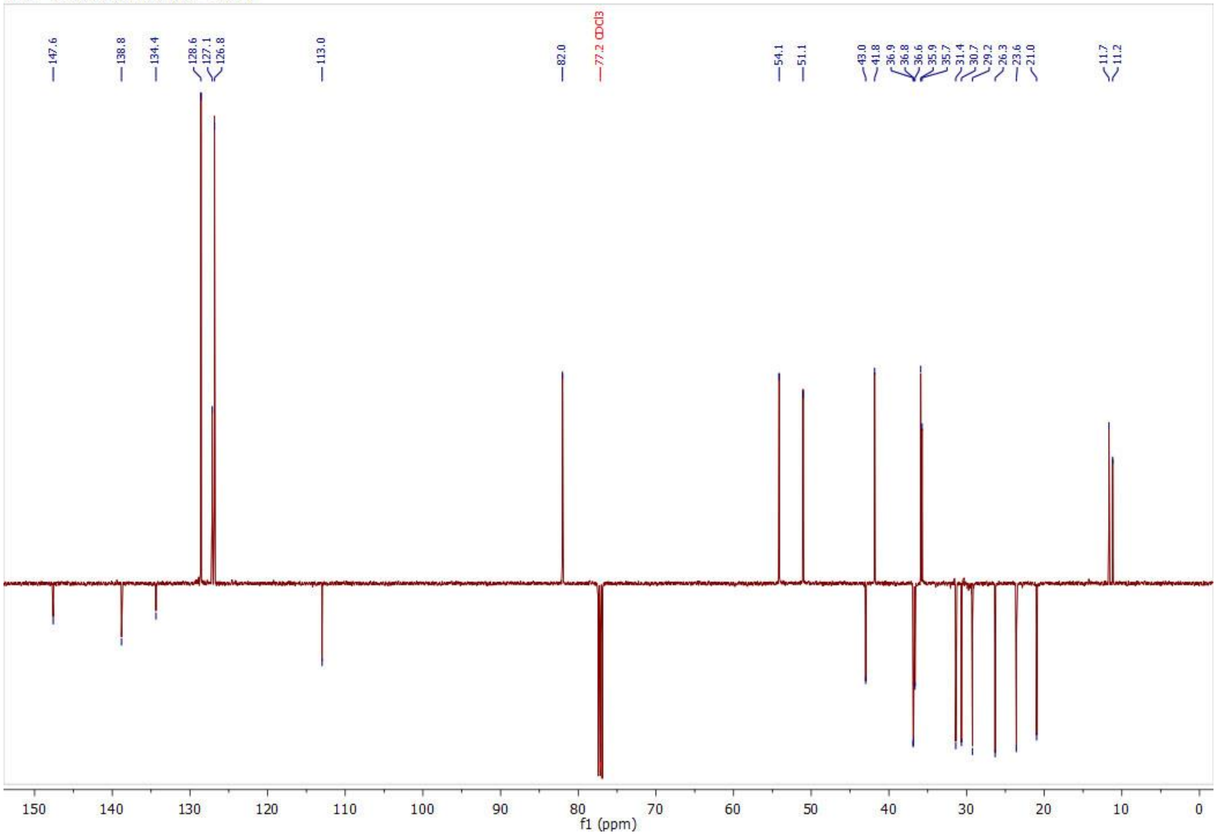

1H — 2021-04-28T19:54:26 — CDCl3

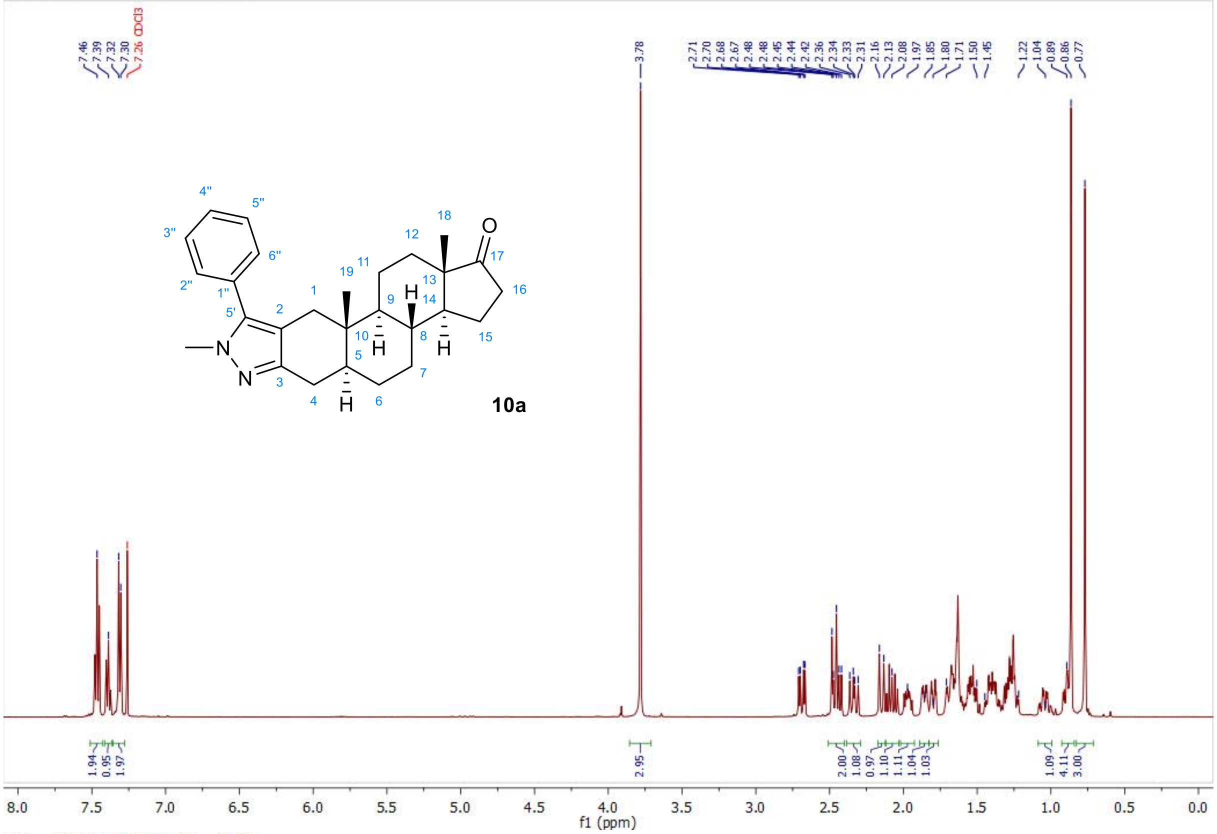


13C — 2021-04-28T20:08:04 — CDCl3

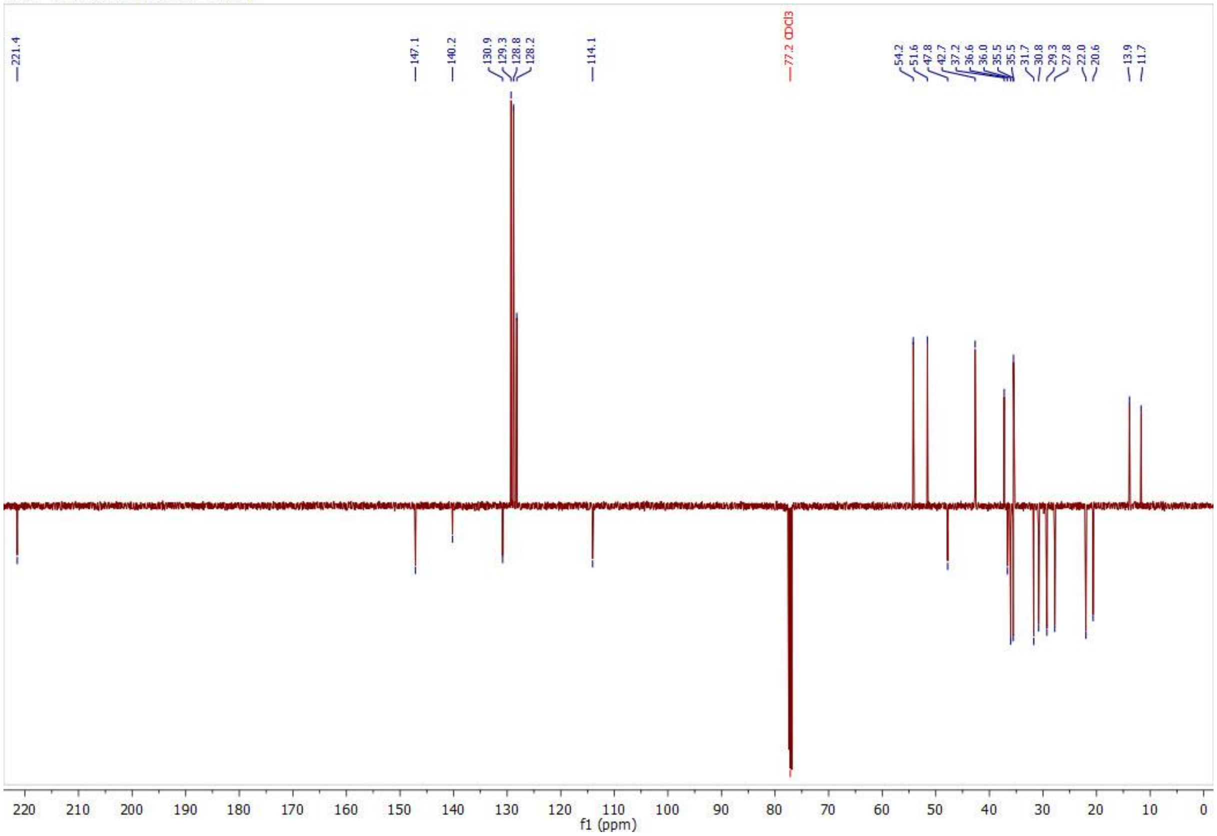

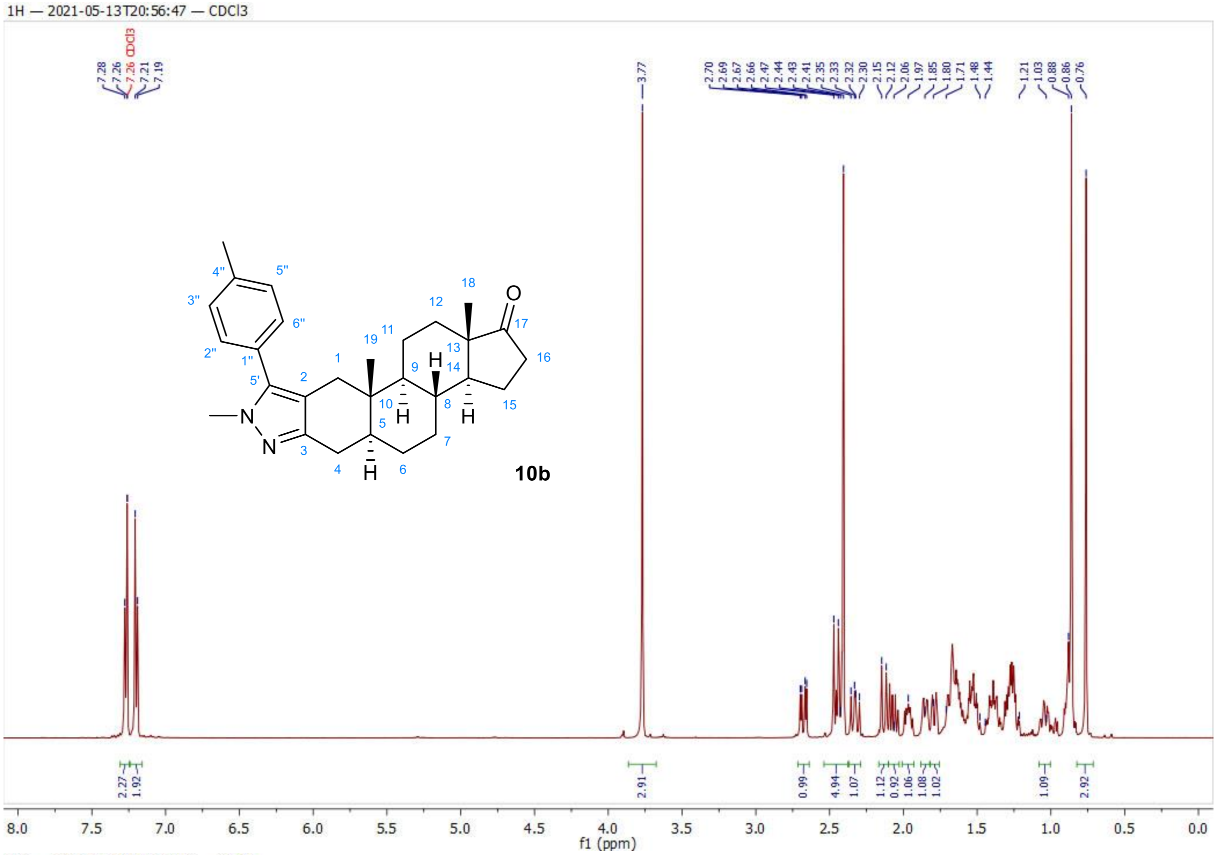
1H — 2021-05-13T20:56:47 — CDCl3
7.28
7.26
7.26 CDCl3
7.21
7.19
3.77
2.70
2.69
2.67
2.66
2.47
2.44
2.43
2.41
2.35
2.33
2.32
2.30
2.15
2.12
2.06
1.97
1.85
1.80
1.71
1.48
1.44
1.21
1.03
0.88
0.86
0.76
10b
2.27
1.92
2.91
0.99
4.94
1.07
1.12
0.92
1.06
1.08
1.02
1.09
2.92
8.0 7.5 7.0 6.5 6.0 5.5 5.0 4.5 4.0 3.5 3.0 2.5 2.0 1.5 1.0 0.5 0.0
f1 (ppm)

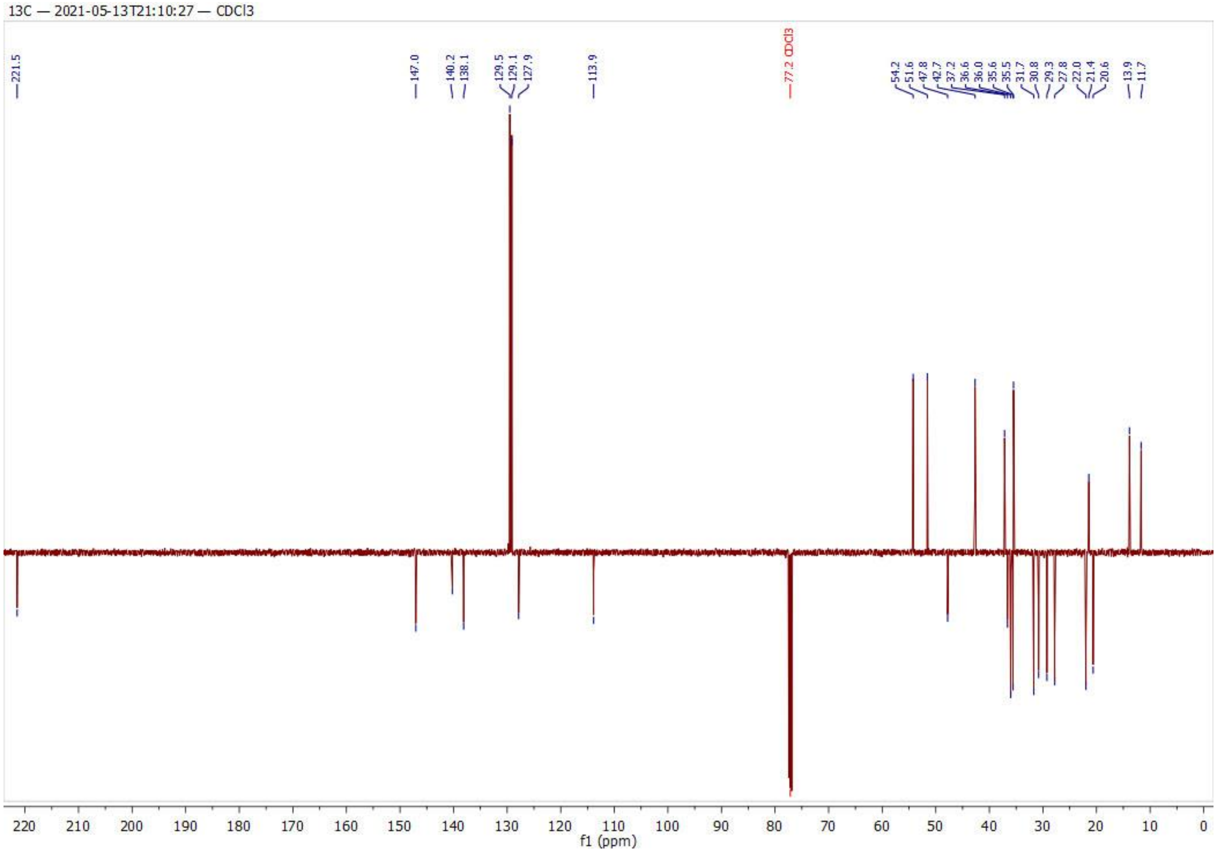
13C — 2021-05-13T21:10:27 — CDCl3
221.5
147.0
140.2
138.1
129.5
129.1
127.9
113.9
77.2 CDCl3
54.2
51.6
47.8
42.7
37.2
36.6
36.0
35.6
35.5
31.7
30.8
29.3
27.8
22.0
21.4
20.6
13.9
11.7
220 210 200 190 180 170 160 150 140 130 120 110 100 90 80 70 60 50 40 30 20 10 0
f1 (ppm)

1H — 2021-04-28T23:00:52 — CDCl3

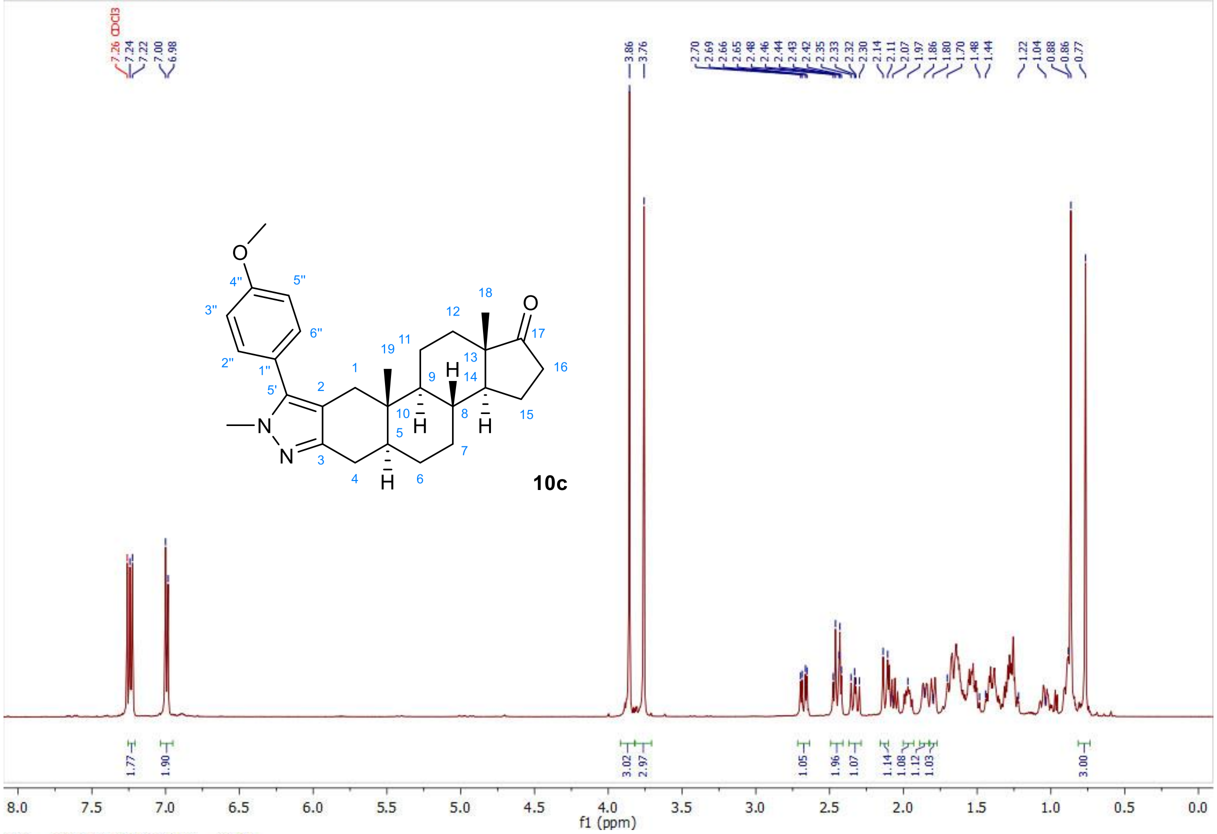


13C — 2021-04-28T23:14:29 — CDCl3

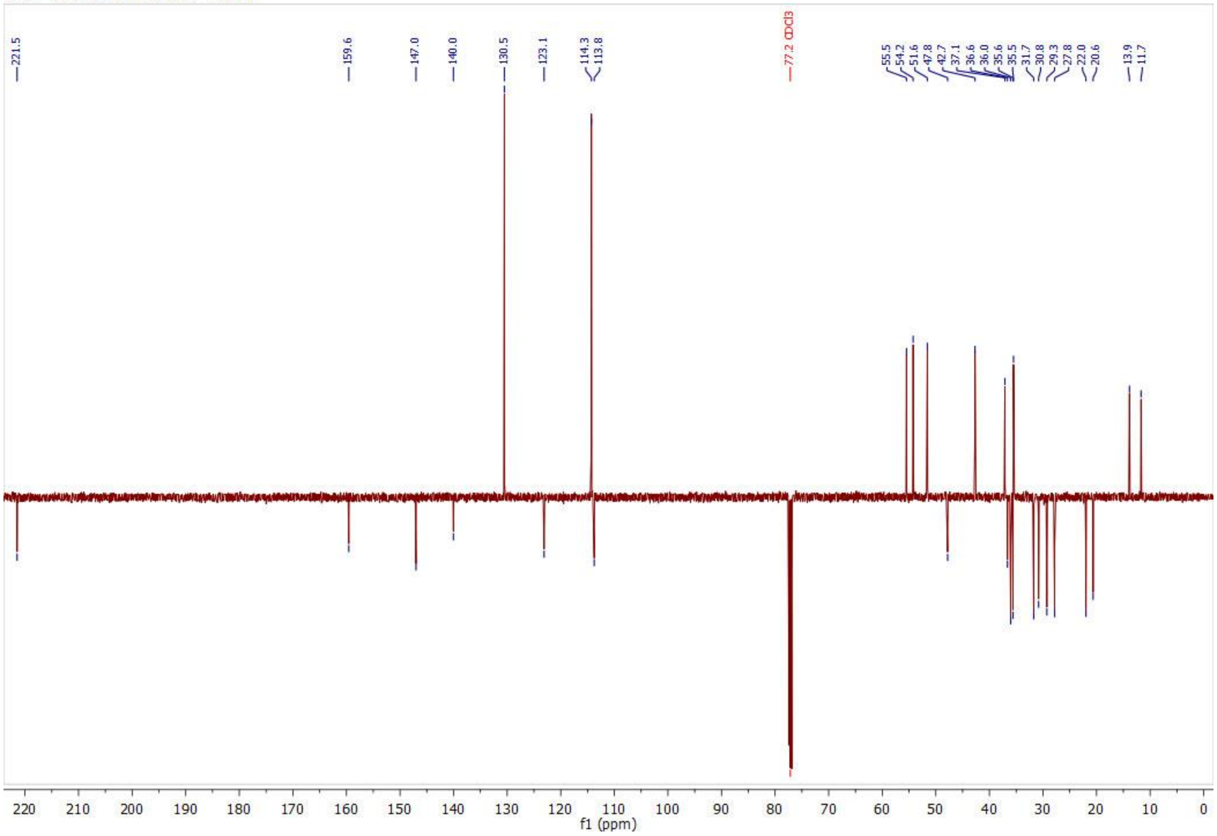

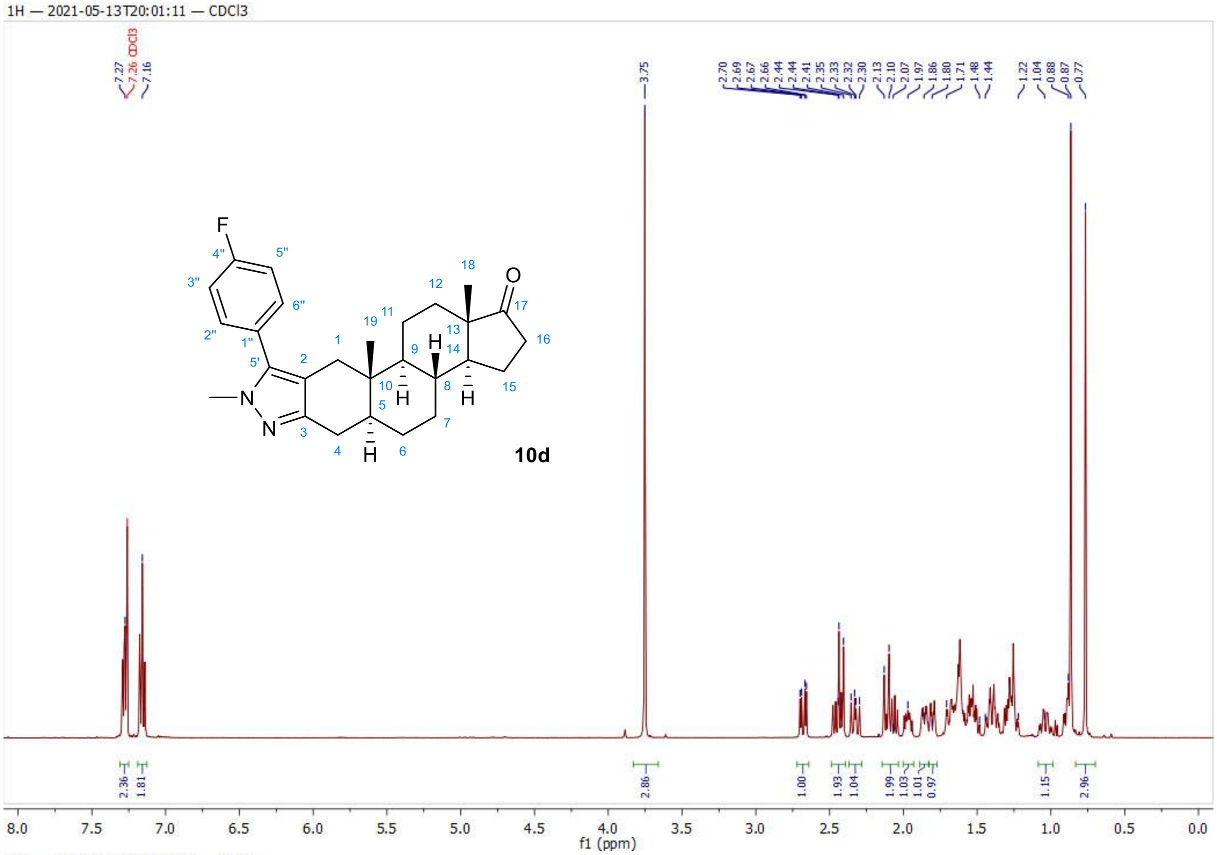
1H — 2021-05-13T20:01:11 — CDCl3
10d
f1 (ppm)

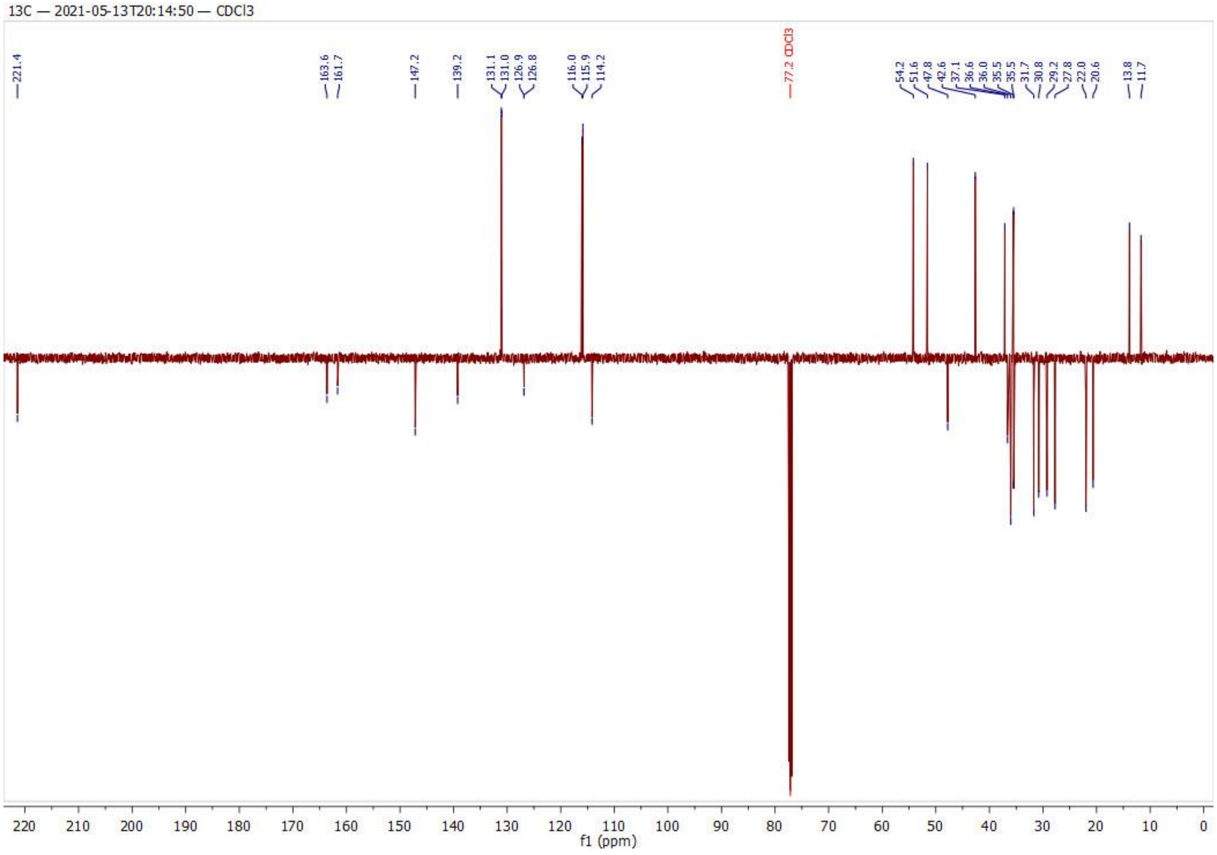
13C — 2021-05-13T20:14:50 — CDCl3
f1 (ppm)

1H — 2021-09-18T01:09:49 — CDCl3

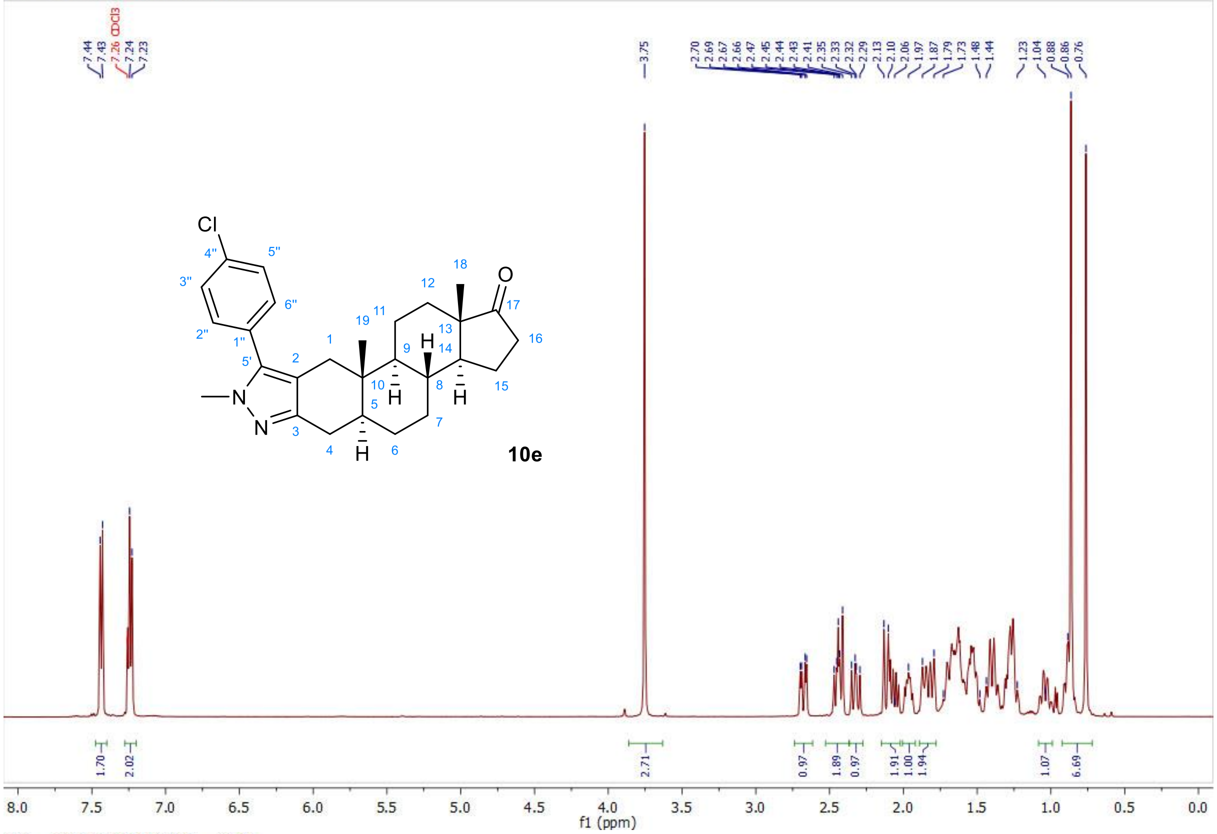


13C — 2021-09-18T01:23:25 — CDCl3

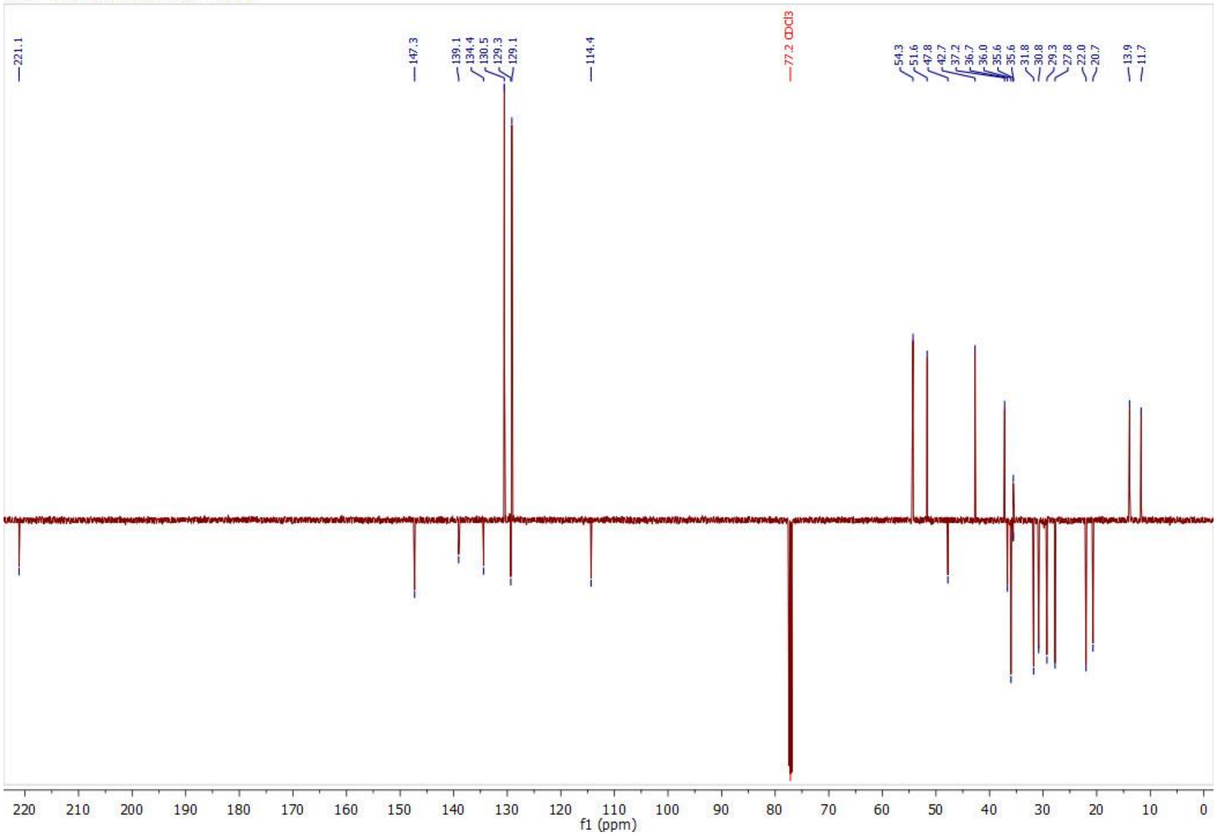

1H — 2021-04-28T19:36:38 — CDCl3

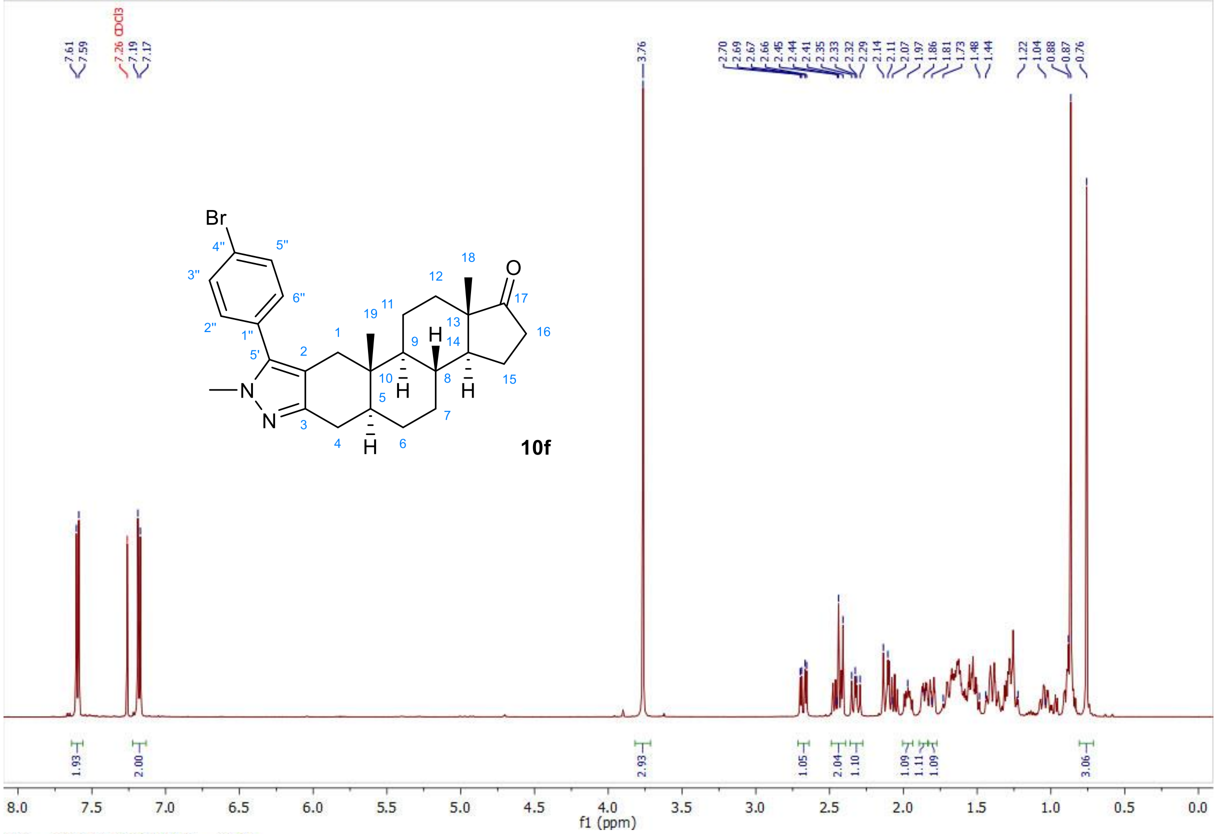


13C — 2021-04-28T19:50:15 — CDCl3

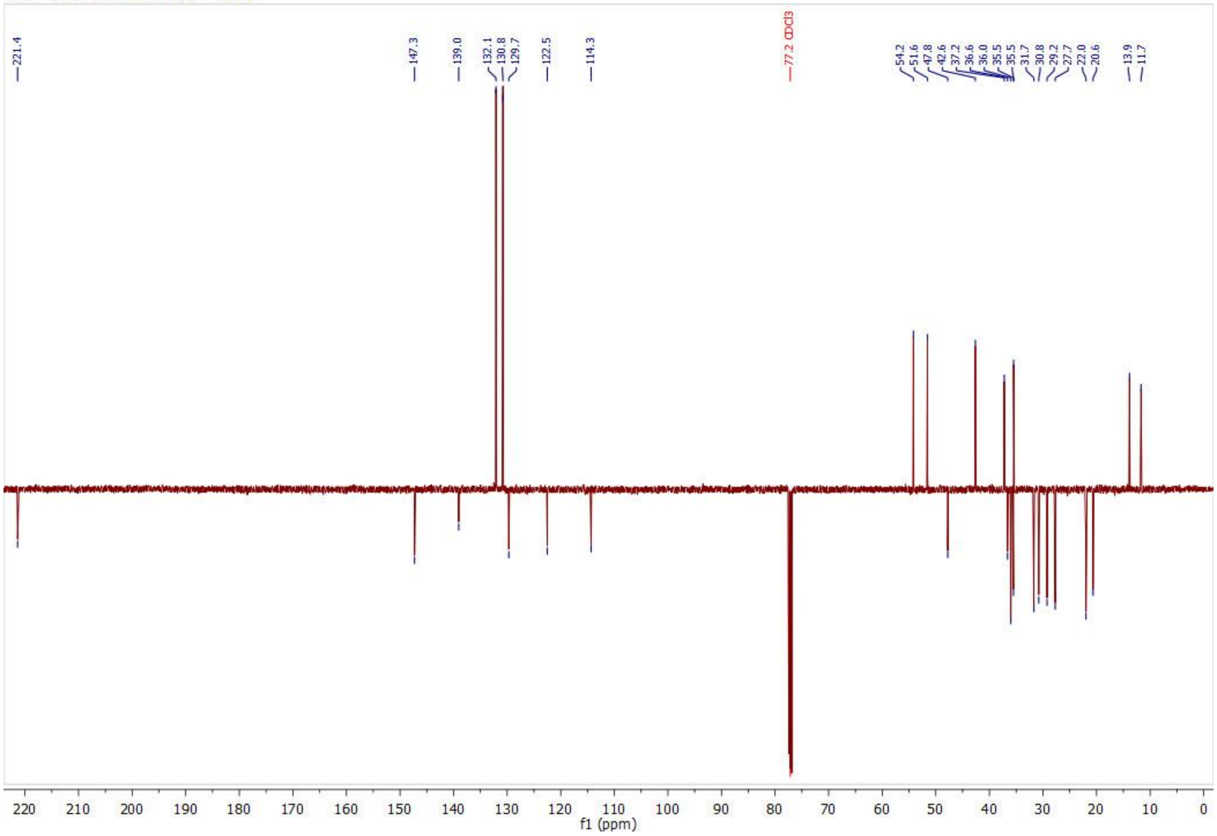

1H — 2021-05-13T20:19:12 — CDCl3

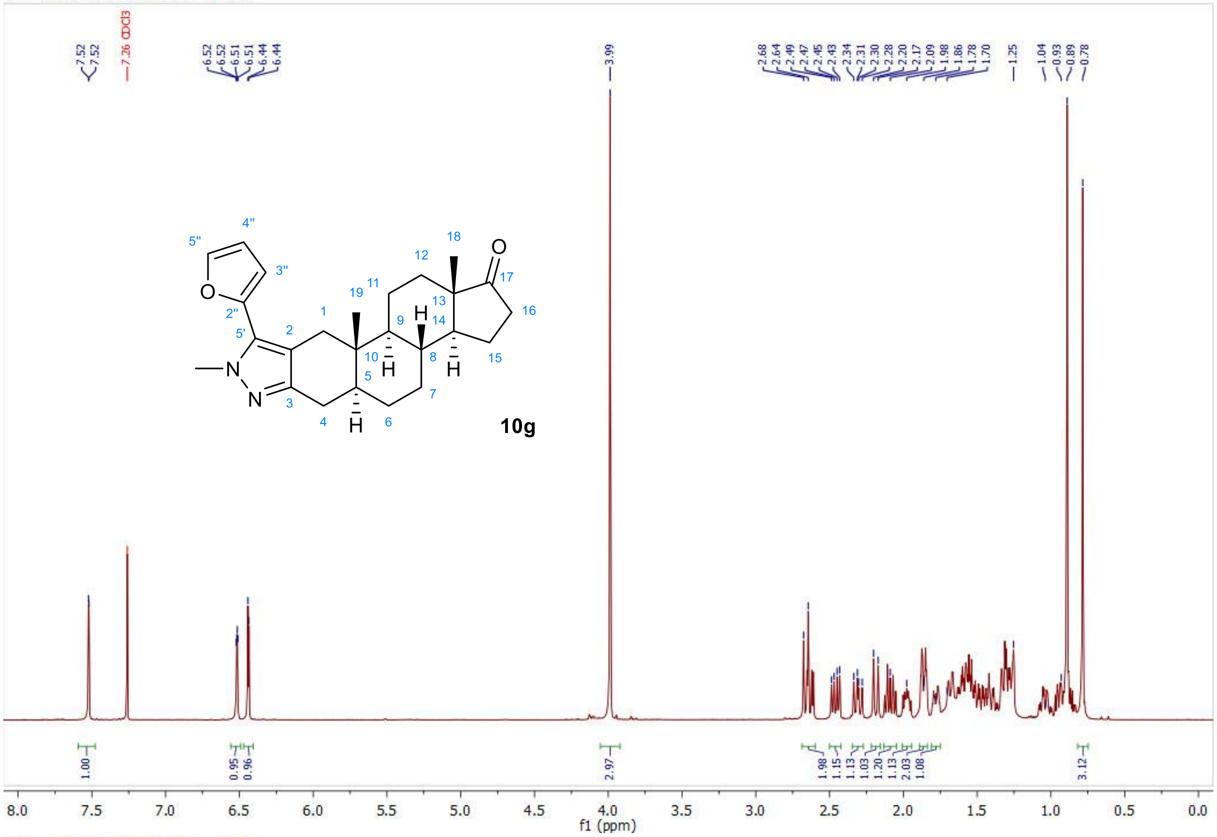


13C — 2021-05-13T20:32:53 — CDCl3

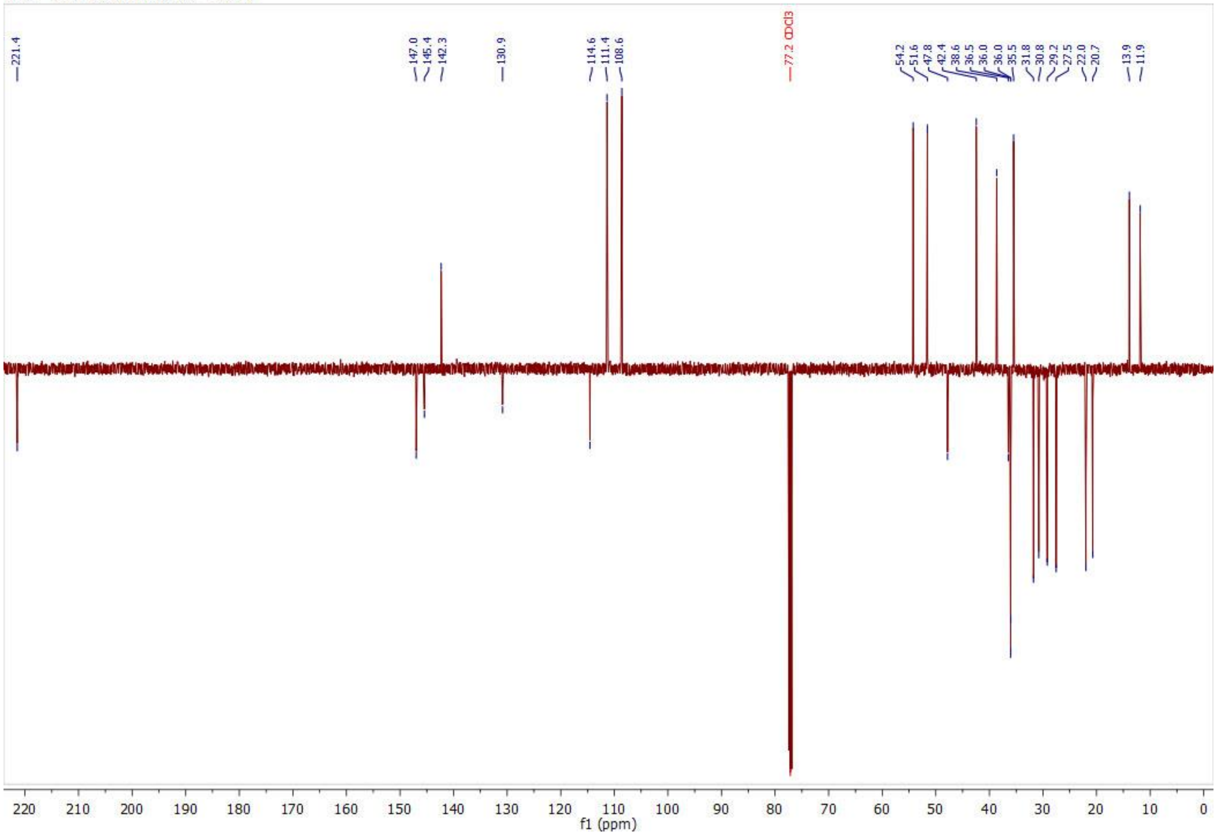

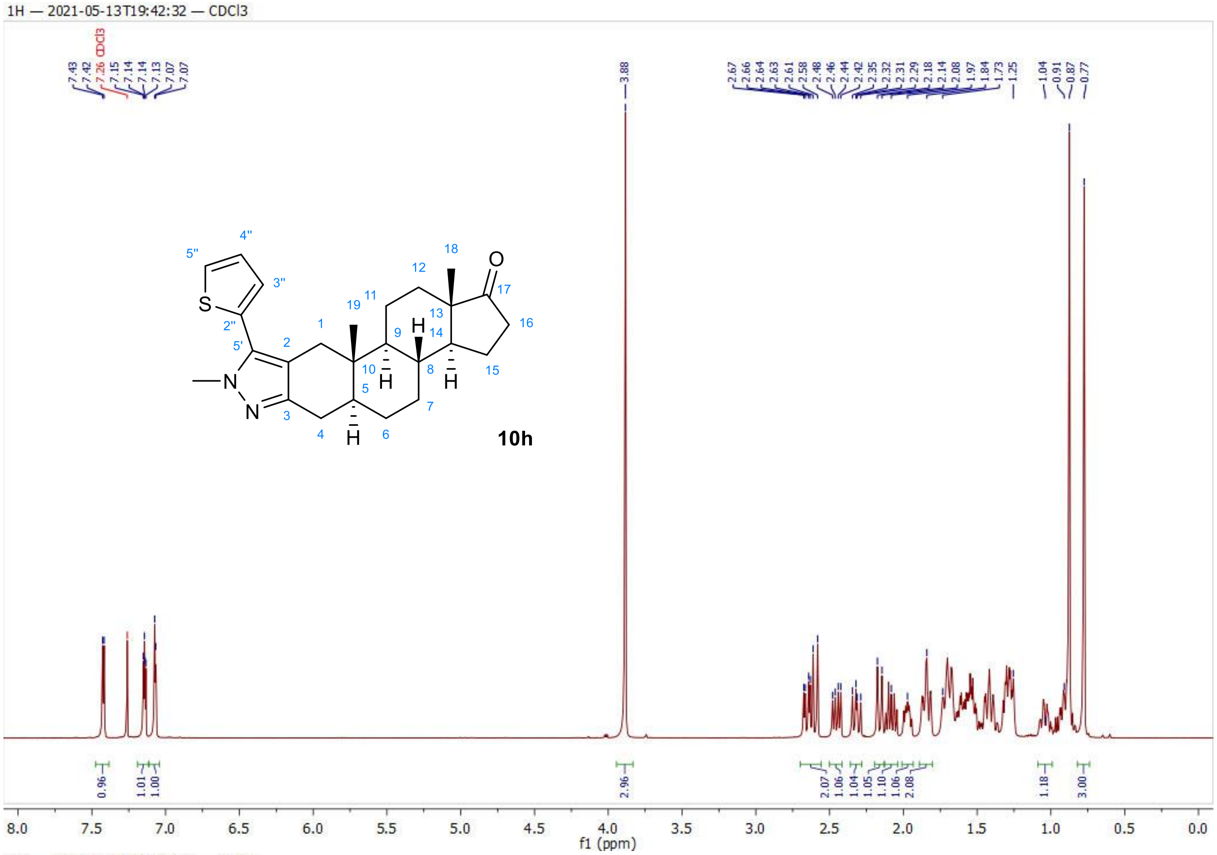
1H — 2021-05-13T19:42:32 — CDCl3
10h
f1 (ppm)

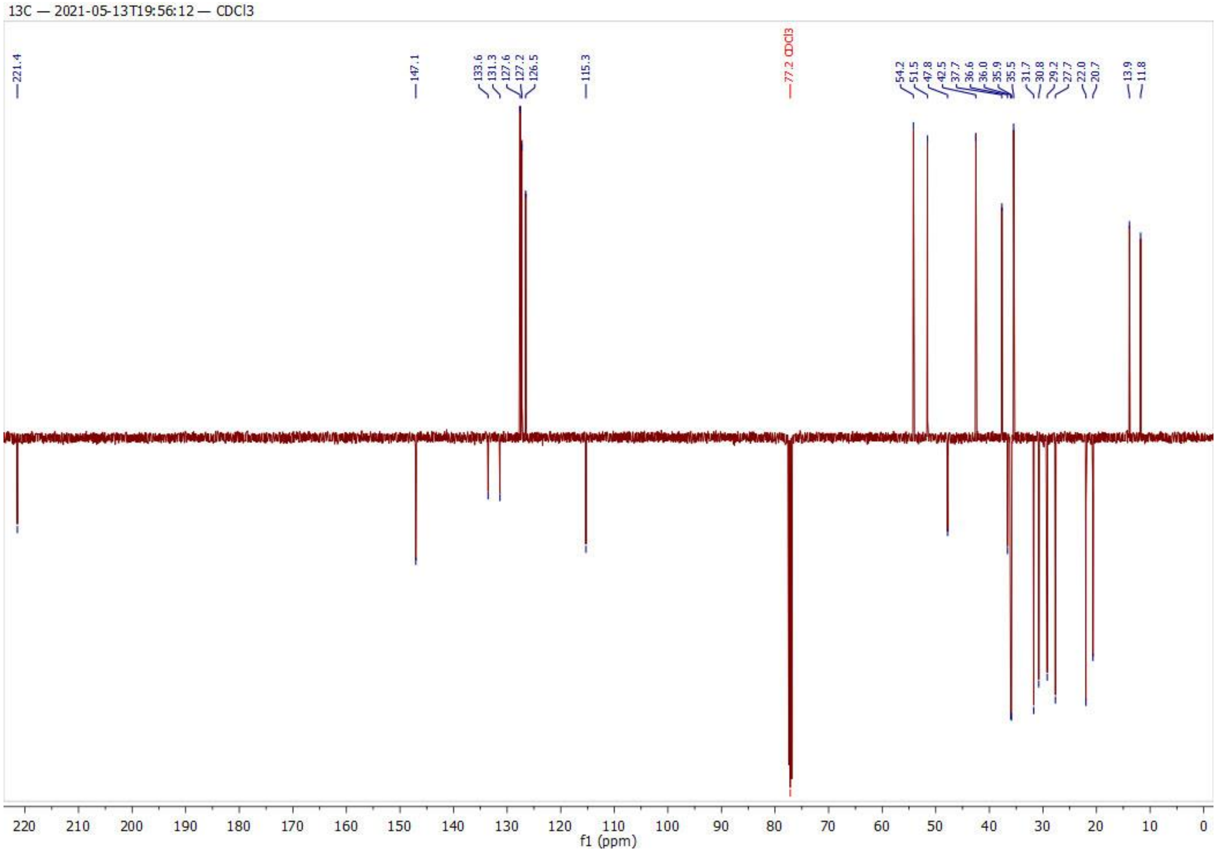
13C — 2021-05-13T19:56:12 — CDCl3
f1 (ppm)

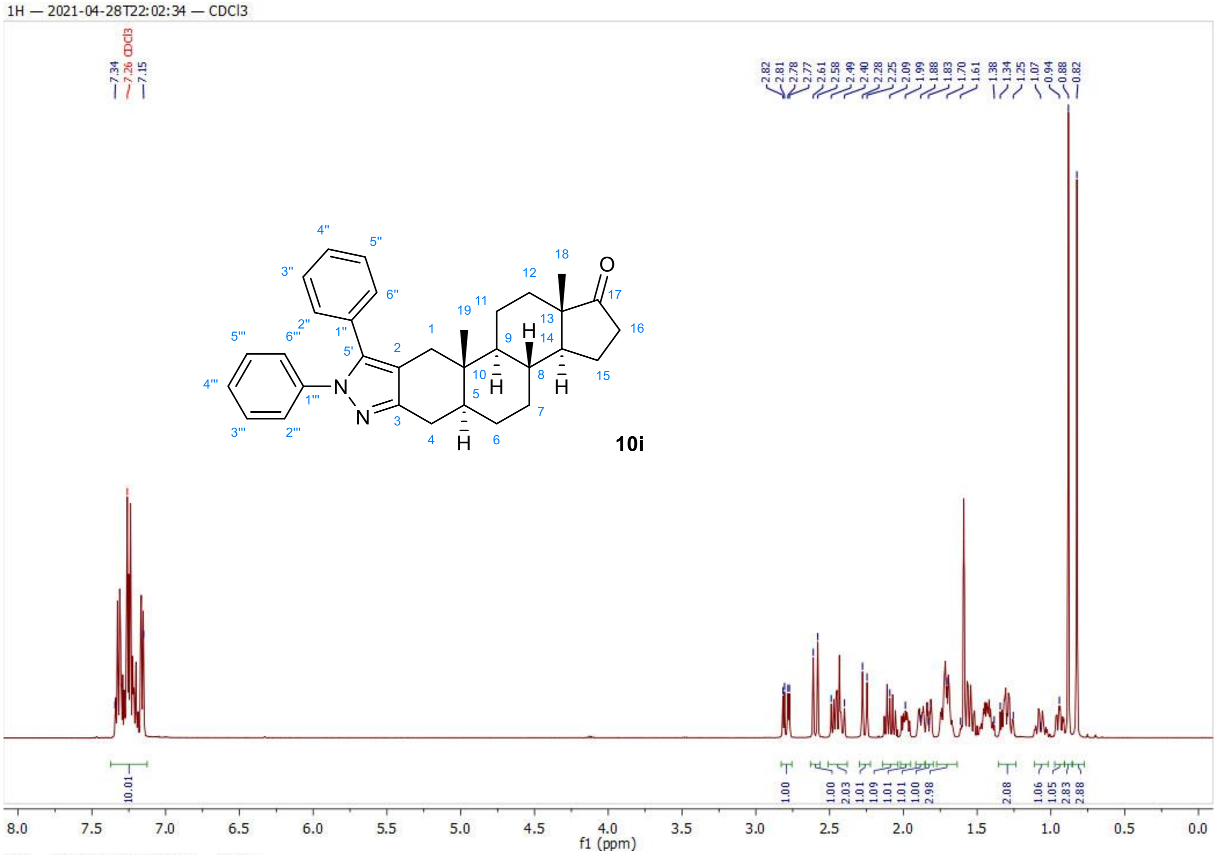
1H — 2021-04-28T22:02:34 — CDCl3
7.34
7.26 CDCl3
7.15
2.82
2.81
2.78
2.77
2.61
2.58
2.49
2.40
2.28
2.25
2.09
1.99
1.88
1.83
1.70
1.61
1.38
1.34
1.25
1.07
0.94
0.88
0.82
10i
10.01
1.00
1.00
2.03
1.01
1.09
1.01
1.01
1.00
2.98
2.08
1.06
1.05
2.83
2.88
f1 (ppm)

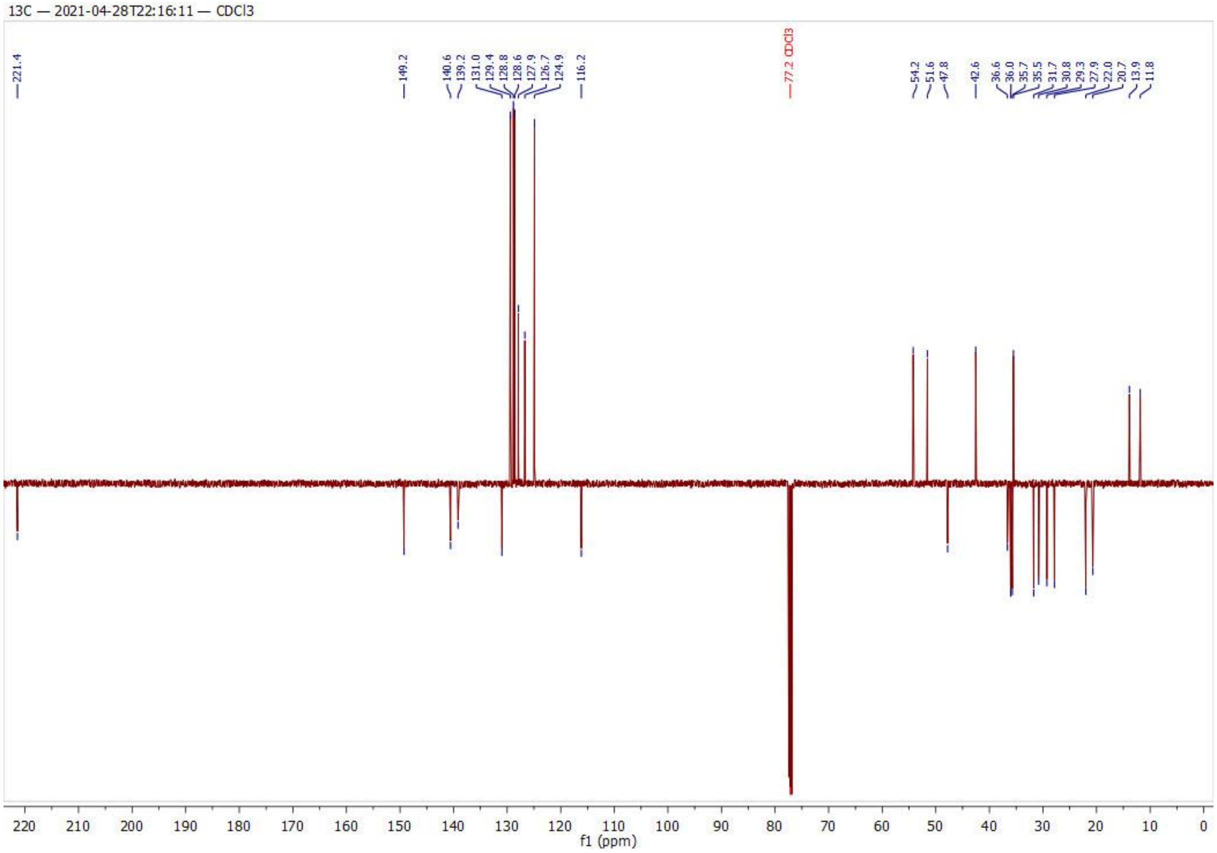
13C — 2021-04-28T22:16:11 — CDCl3
221.4
149.2
140.6
139.2
131.0
129.4
128.8
128.6
127.9
126.7
124.9
116.2
77.2 CDCl3
54.2
51.6
47.8
42.6
36.6
36.0
35.7
35.5
31.7
30.8
29.3
27.9
22.0
20.7
13.9
11.8
f1 (ppm)

1H — 2021-04-28T22:37:58 — CDCl3

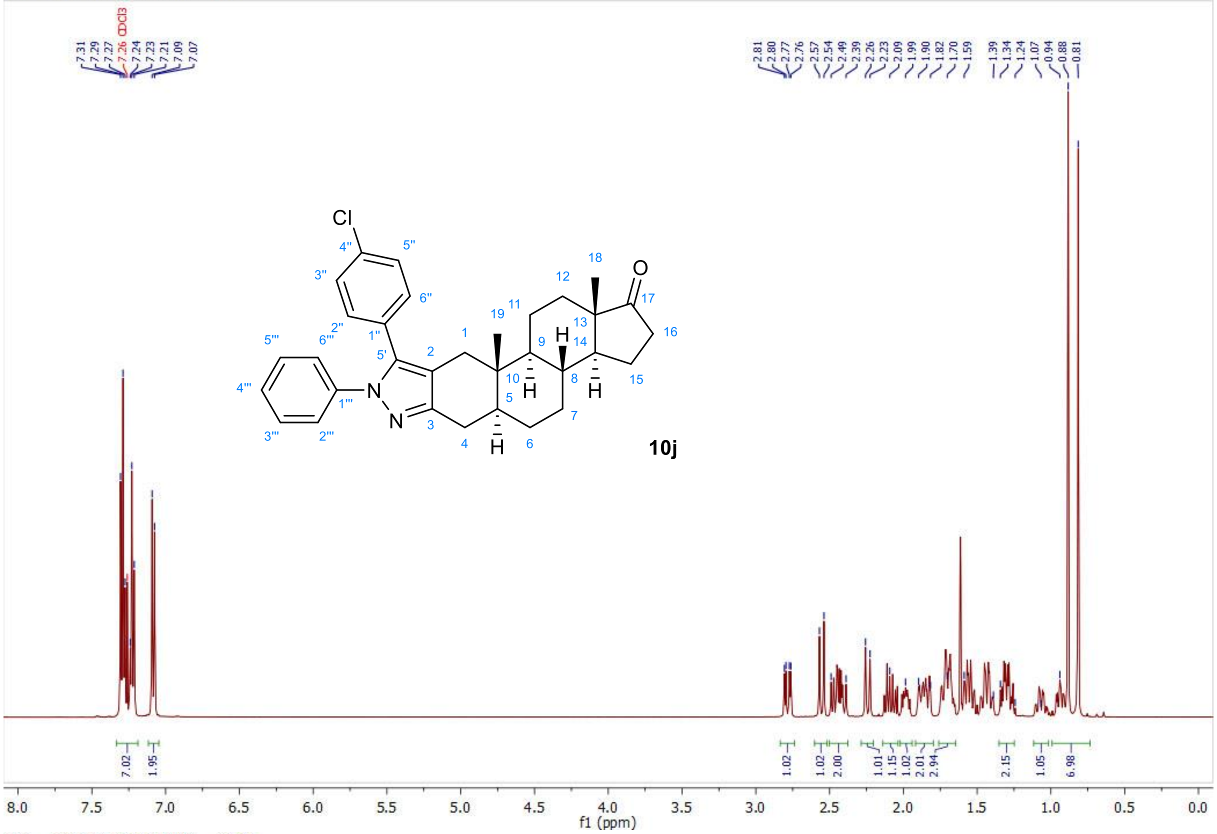


13C — 2021-04-28T22:51:36 — CDCl3

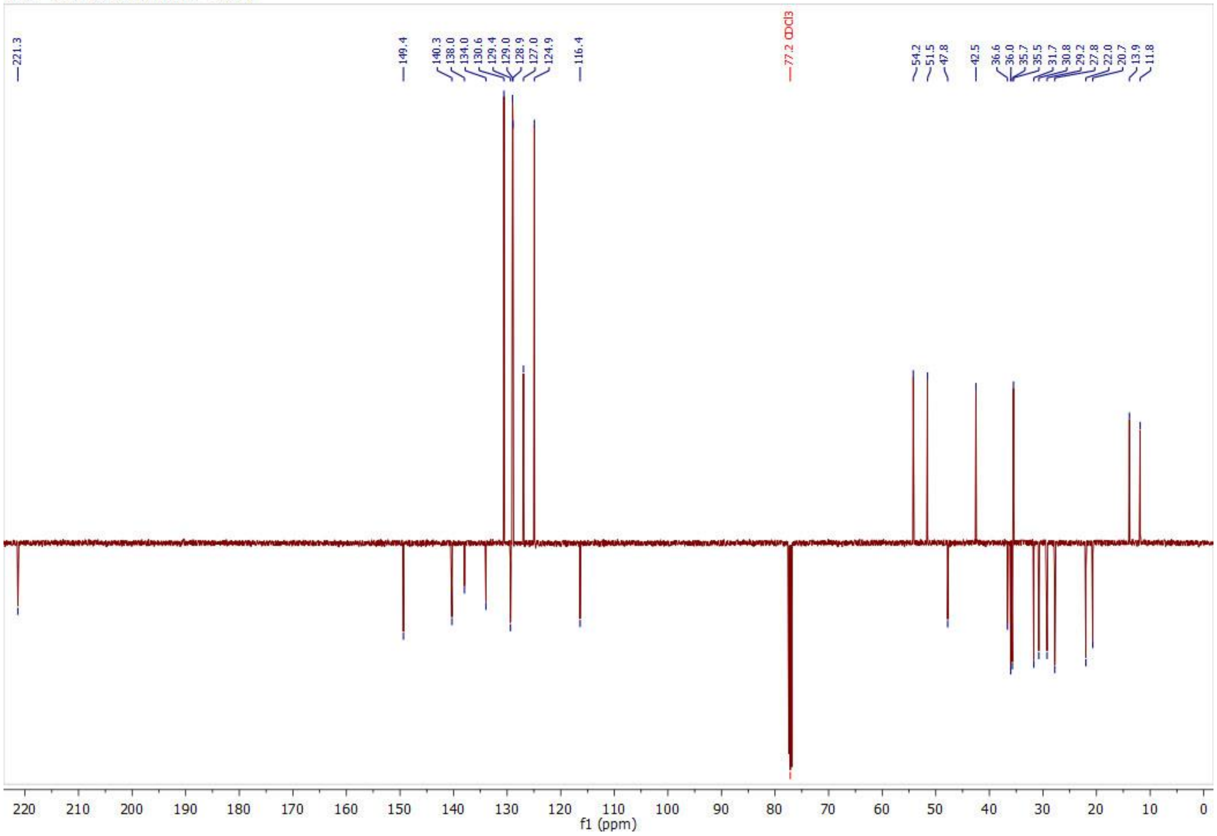

1H — 2021-05-13T21:14:47 — CDCl3

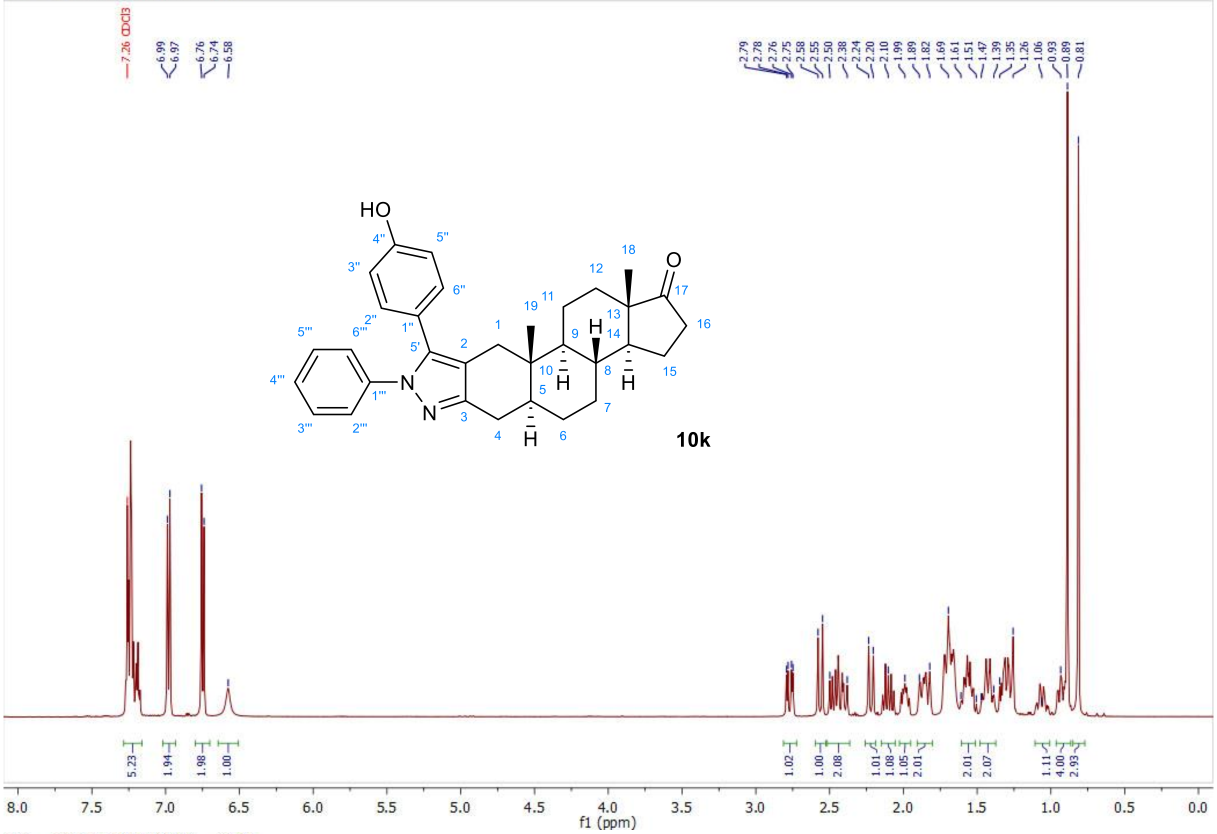


13C — 2021-05-13T21:28:26 — CDCl3

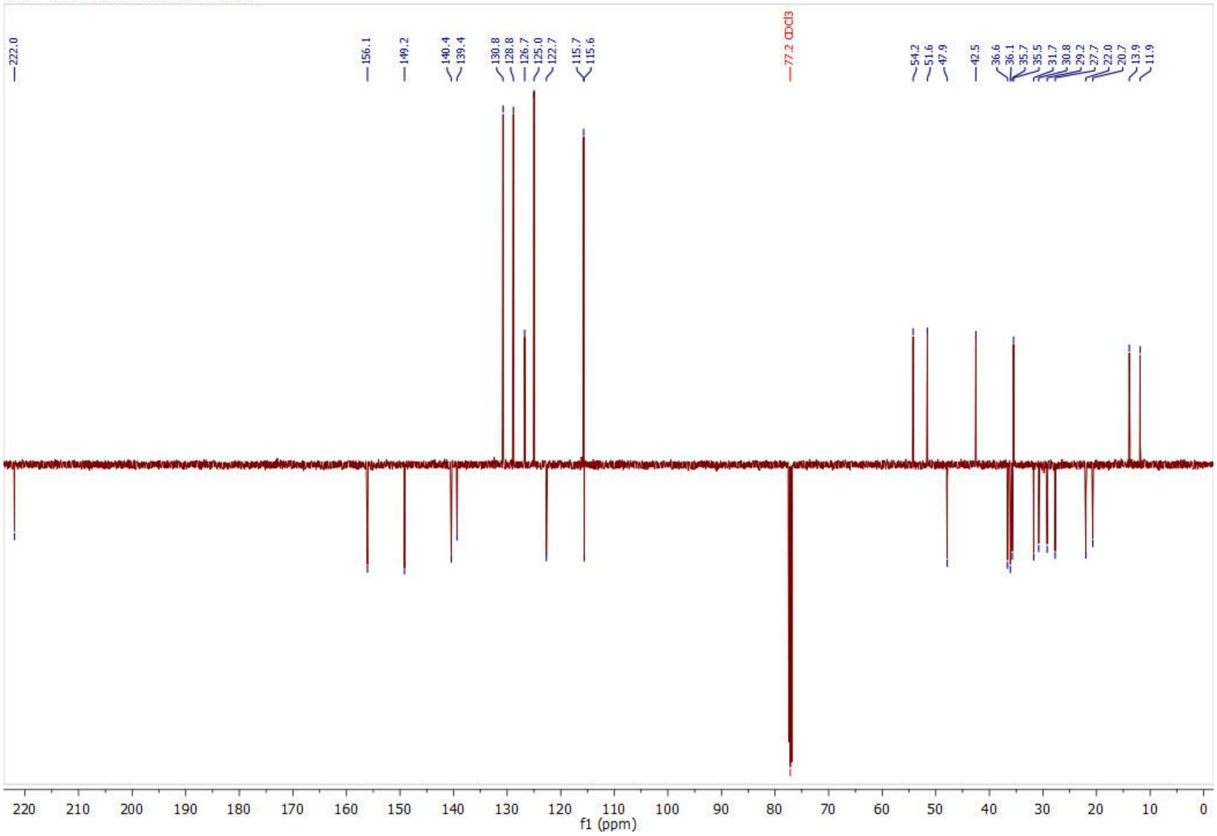

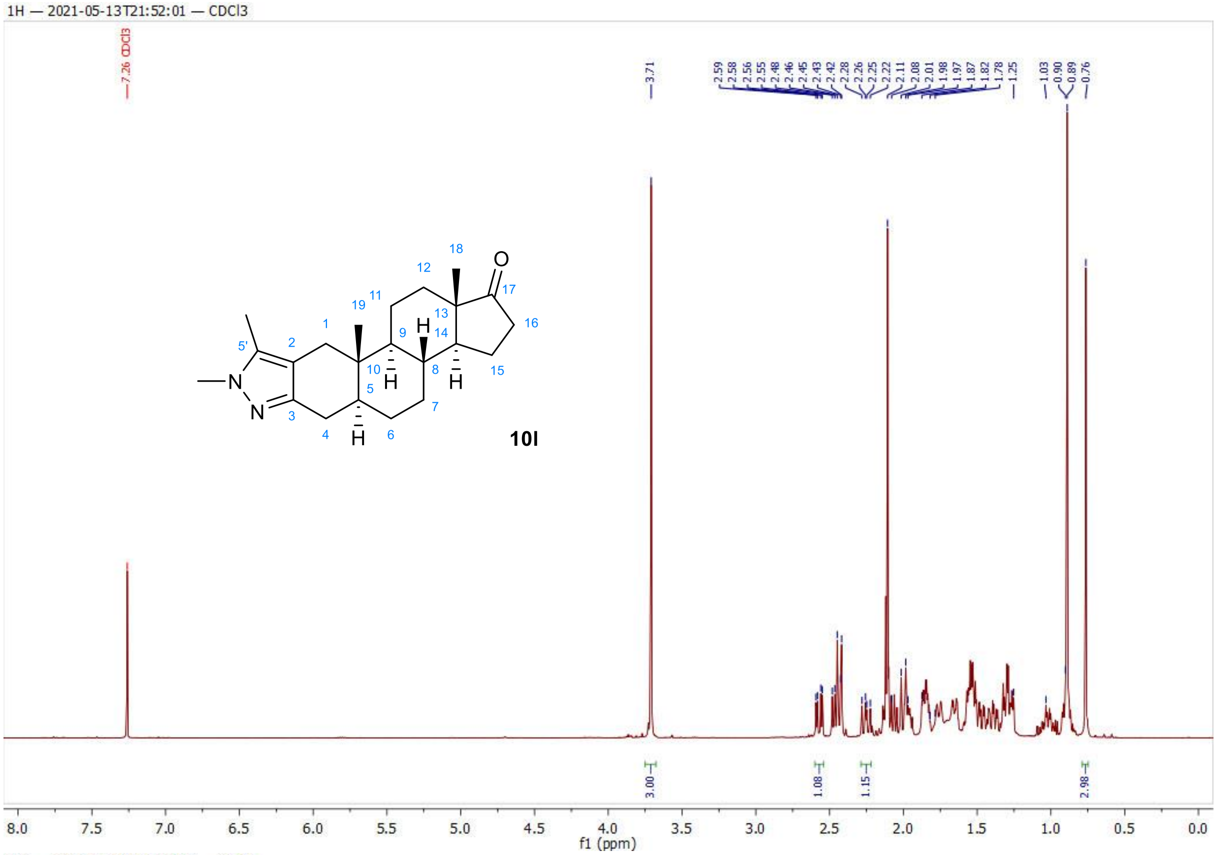
1H — 2021-05-13T21:52:01 — CDCl3
7.26 CDCl3
3.71
2.59
2.58
2.56
2.55
2.48
2.46
2.45
2.43
2.42
2.28
2.26
2.25
2.22
2.11
2.08
2.01
1.98
1.97
1.87
1.82
1.78
1.25
1.03
0.90
0.89
0.76
3.00
1.08
1.15
2.98
10l
f1 (ppm)

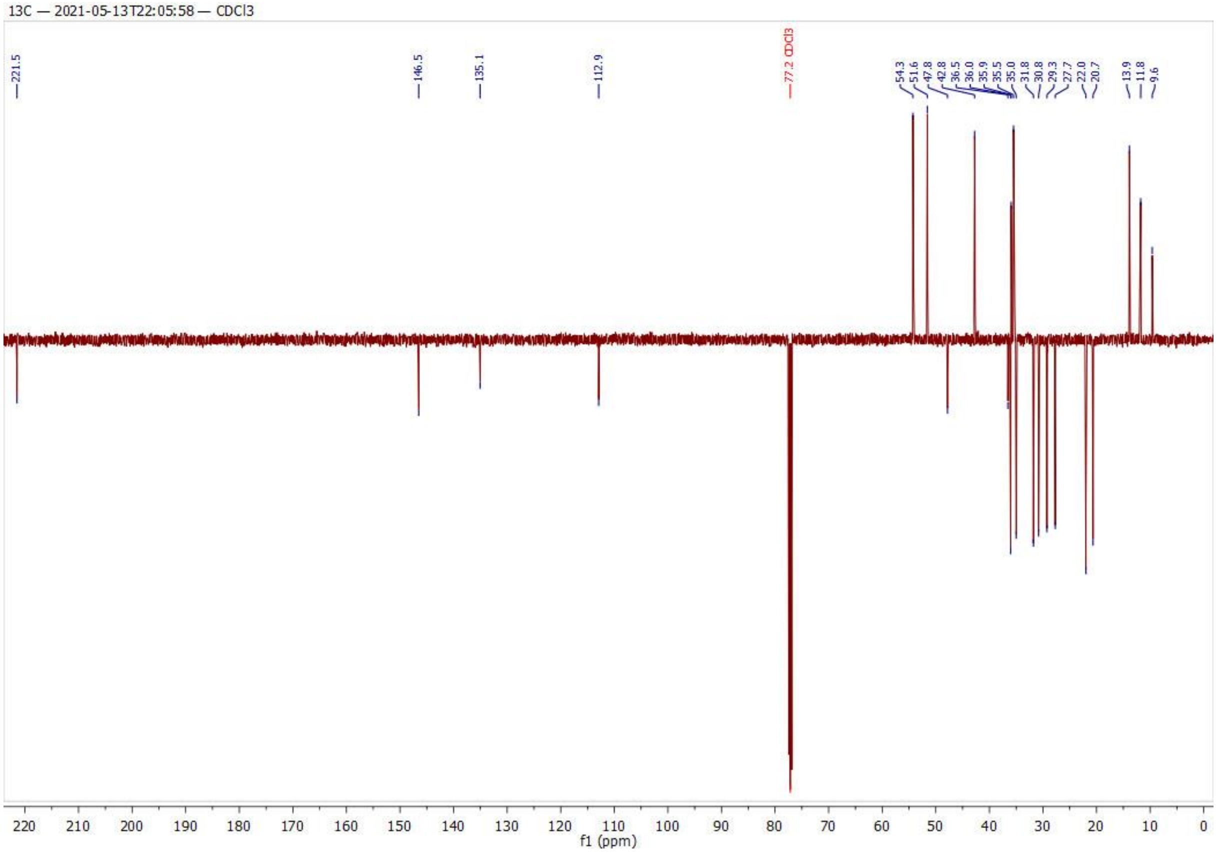
13C — 2021-05-13T22:05:58 — CDCl3
221.5
146.5
135.1
112.9
77.2 CDCl3
54.3
51.6
47.8
42.8
36.5
36.0
35.9
35.5
35.0
31.8
30.8
29.3
27.7
22.0
20.7
13.9
11.8
9.6
f1 (ppm)

**Table S1.** AR transcriptional activity in antagonist and agonist mode upon treatment with displayed concentrations of compounds.

| | AR transcriptional activity (% of control stimulated with 1 nM metribolone) ANTAGONIST | | | AR transcriptional activity (% of control stimulated with 1 nM metribolone) AGONIST | | |
|---|---|---|---|---|---|---|
| Cmp | 10 µM | 2 µM | 0.4 µM | 10 µM | 2 µM | 0.4 µM |
| **1a** | 53.7 | 90.2 | 96.7 | 73.8 | 69.3 | 63.1 |
| **1b** | 51.1 | 72.4 | 97.0 | 66.1 | 69.4 | 58.9 |
| **1c** | 48.5 | 73.4 | 92.8 | 77.4 | 83.2 | 83.9 |
| **1d** | 37.9 | 65.9 | 92.5 | 48.5 | 38.1 | 32.1 |
| **1e** | 39.2 | 64.6 | 84.2 | 43.8 | 48.3 | 42.9 |
| **1f** | 66.1 | 73.2 | 80.2 | 17.7 | 28.6 | 26.2 |
| **1g** | 69.4 | 76.9 | 78.9 | 84.1 | 72.7 | 65.4 |
| **1h** | 73.6 | 68.0 | 84.9 | 102.9 | 91.8 | 67.1 |
| **2a** | 47.5 | 75.2 | 94.5 | 72.6 | 59.2 | 56.2 |
| **2b** | 93.1 | 103.3 | 109.4 | 103.7 | 87.4 | 78.2 |
| **2c** | 67.3 | 83.8 | 96.8 | 108.0 | 84.3 | 67.9 |
| **2d** | 67.1 | 83.9 | 110.3 | 89.2 | 64.0 | 41.7 |
| **2e** | 107.0 | 120.9 | 100.7 | 87.1 | 72.2 | 46.6 |
| **2f** | 78.7 | 86.8 | 77.1 | 35.0 | 21.6 | 8.0 |
| **2g** | 67.7 | 81.6 | 86.2 | 48.5 | 57.6 | 54.0 |
| **2h** | 90.8 | 83.0 | 91.4 | 78.5 | 66.4 | 55.7 |
| **3a** | 16.6 | 46.7 | 84.5 | 16.4 | 26.7 | 26.4 |
| **3b** | 33.1 | 64.4 | 92.4 | 17.3 | 21.0 | 26.4 |
| **3c** | 34.0 | 53.1 | 80.7 | 25.2 | 29.1 | 33.8 |
| **3d** | 13.7 | 42.8 | 88.0 | 17.8 | 25.4 | 27.7 |
| **3e** | 21.7 | 53.9 | 93.2 | 21.7 | 25.4 | 26.1 |
| **3f** | 35.7 | 61.6 | 82.4 | 27.2 | 28.9 | 27.1 |
| **3g** | 18.7 | 36.8 | 72.8 | 20.7 | 23.8 | 26.1 |
| **3h** | 28.3 | 63.4 | 91.8 | 25.2 | 34.9 | 31.1 |
| **4a** | 71.2 | 82.1 | 98.3 | 31.4 | 27.9 | 23.9 |
| **4b** | 85.3 | 89.4 | 87.1 | 20.1 | 25.3 | 26.4 |
| **4c** | 78.3 | 108.6 | 119.6 | 25.7 | 31.2 | 32.7 |
| **4d** | 30.6 | 64.7 | 88.0 | 18.9 | 22.4 | 26.7 |
| **4e** | 43.3 | 97.2 | 108.2 | 17.7 | 25.8 | 26.4 |
| **4g** | 47.9 | 66.5 | 79.8 | 26.1 | 33.8 | 29.9 |
| **4h** | 54.6 | 87.6 | 98.0 | 27.3 | 33.5 | 32.7 |
| **8a** | 88.7 | 90.5 | 92.2 | 11.4 | 17.0 | 18.5 |
| **8b** | 70.4 | 99.5 | 89.1 | 41.7 | 42.3 | 25.9 |
| **8c** | 65.5 | 89.3 | 92.9 | 1.7 | 7.4 | 7.3 |
| **8d** | 63.7 | 85.0 | 92.8 | 9.0 | 16.3 | 20.1 |
| **8e** | 44.6 | 79.9 | 95.5 | 10.9 | 30.4 | 23.5 |
| **8f** | 34.5 | 80.6 | 95.5 | 6.3 | 18.0 | 15.9 |

| | | | | | | |
|---|---|---|---|---|---|---|
| **8g** | 13.0 | 67.4 | 81.5 | 0.9 | 15.2 | 14.4 |
| **8h** | 37.4 | 75.6 | 89.9 | 4.1 | 24.6 | 25.8 |
| **8i** | 139.1 | 113.4 | 95.3 | 57.0 | 43.9 | 22.6 |
| **8j** | 121.6 | 108.5 | 111.9 | 67.6 | 46.9 | 27.9 |
| **8k** | 156.8 | 106.4 | 100.6 | 67.6 | 49.7 | 25.2 |
| **8l** | 138.5 | 94.9 | 89.7 | 80.8 | 66.1 | 37.5 |
| **10a** | 100.1 | 105.5 | 105.9 | 36.3 | 21.6 | 8.1 |
| **10b** | 82.2 | 94.5 | 95.5 | 2.8 | 4.6 | 8.4 |
| **10c** | 93.5 | 98.4 | 103.7 | 29.6 | 20.3 | 13.0 |
| **10d** | 86.2 | 99.3 | 109.7 | 6.4 | 6.9 | 10.7 |
| **10e** | 40.4 | 84.1 | 92.1 | 2.6 | 4.3 | 7.7 |
| **10f** | 40.1 | 88.5 | 95.1 | 3.2 | 5.0 | 6.4 |
| **10g** | 59.6 | 99.7 | 100.9 | 14.5 | 24.7 | 22.7 |
| **10h** | 59.5 | 94.1 | 96.3 | 11.1 | 16.7 | 18.9 |
| **10i** | 136.3 | 115.8 | 97.5 | 65.4 | 42.6 | 25.1 |
| **10j** | 168.8 | 124.1 | 114.9 | 67.7 | 43.3 | 24.5 |
| **10k** | 165.0 | 131.5 | 119.0 | 64.1 | 41.0 | 23.2 |
| **10l** | 149.3 | 104.1 | 99.0 | 56.7 | 47.6 | 26.0 |

*Transcriptional activity of AR was measured with compounds alone (agonist mode) or in the presence of 1 nM R1881 (antagonist mode) in reporter cell line ARE14 for 24 h. Activity was normalised to the signal of R1881 (= 100%). Measured in duplicate and repeated twice, mean plotted in the table.*

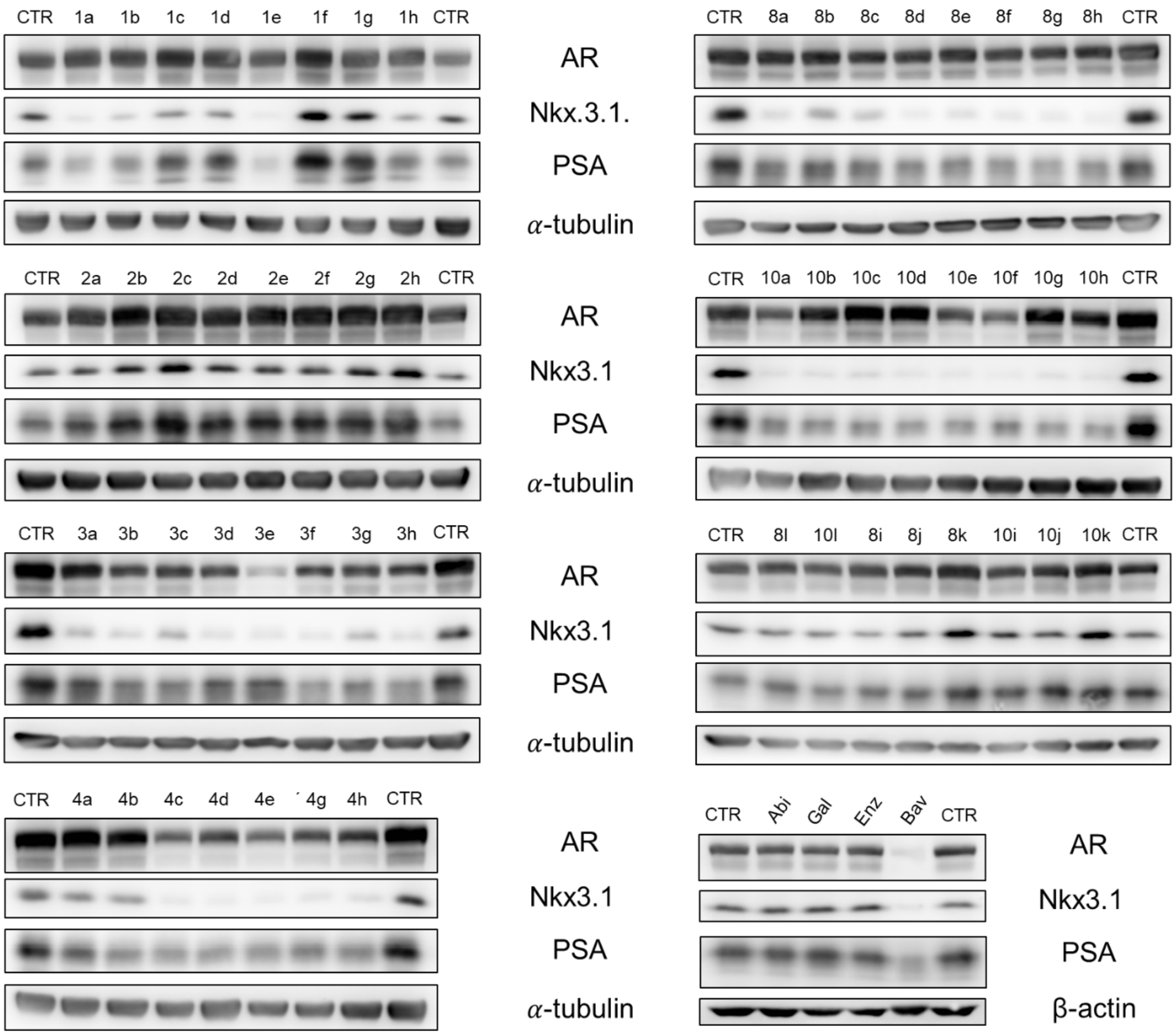


**Figure S3.** (10 µM, 24 h) in FBS containing medium and lysates were then blotted for detection of appropriate Immunoblotting analysis of important signaling proteins from the LAPC-4 cells, which were treated with studied compounds proteins. Level of α-tubulin or β-actin served as loading control. Abi, abiraterone; Gal, galeterone; Enz, enzalutamide; Bav, bavdegalutamide.

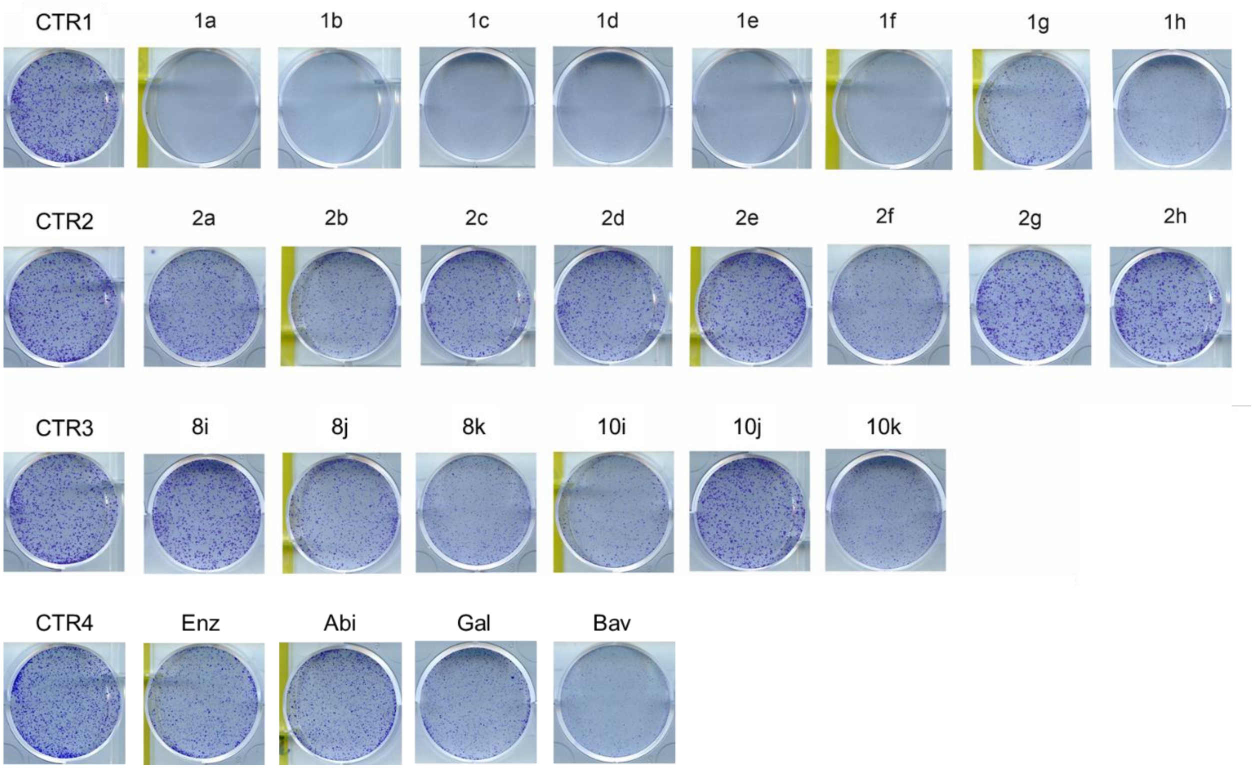


**Figure S4.** Antiproliferative activity of compounds determined by colony formation assay with LAPC-4 cell line. Cells were seeded, treated with 5 μM concentration of compounds and cultivated for 10 days in the presence of compounds. Colonies were fixed and stained as described in experimental part.

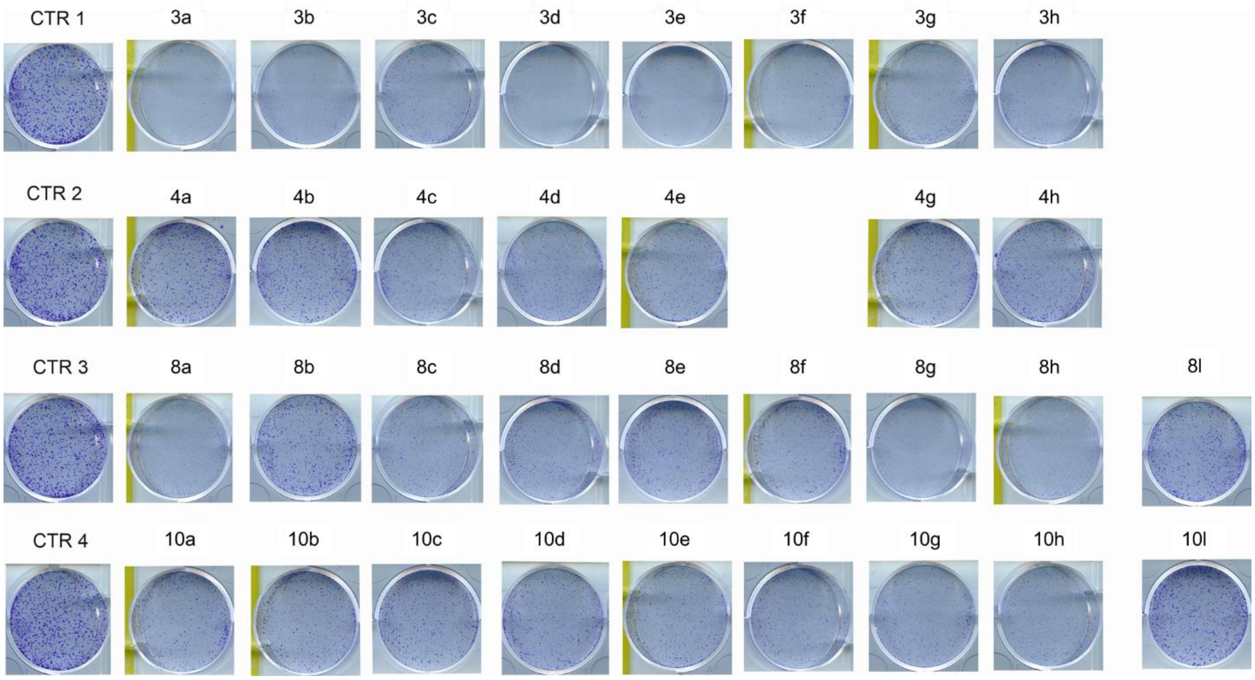


**Figure S5.** Antiproliferative activity of compounds determined by colony formation assay with LAPC-4 cell line. Cells were seeded, treated with 5 μM concentration of compounds and cultivated for 10 days in the presence of compounds. Colonies were fixed and stained as described in experimental part.

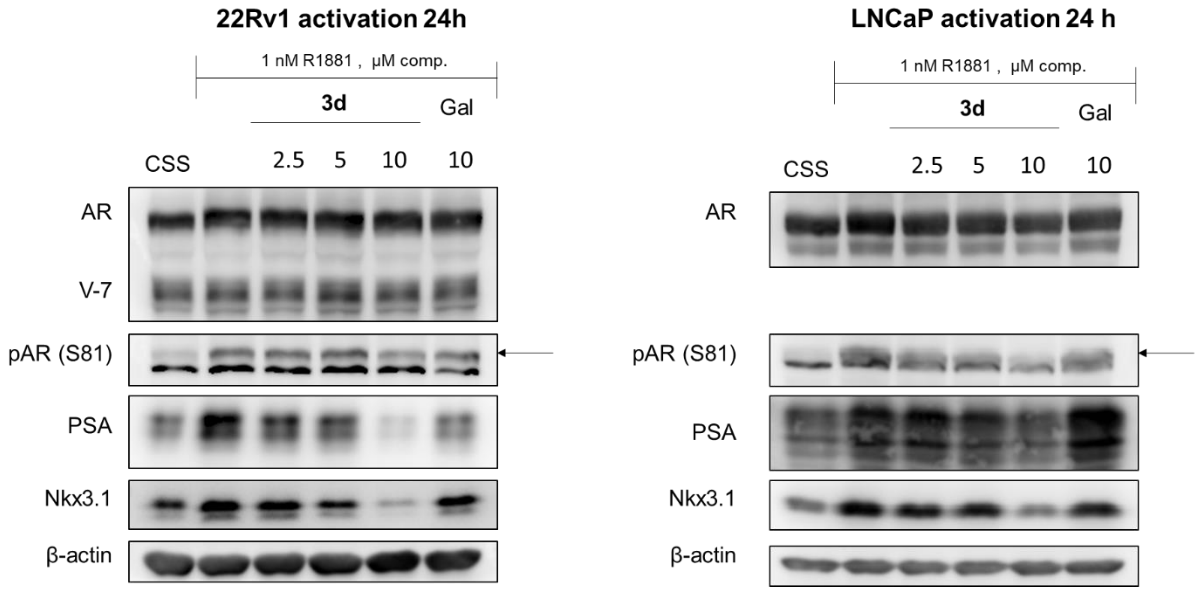


**Figure S6.** Dose dependent effect of **3d** on AR- signalling in 22Rv1, LNCaP. Level of β-actin served as loading control. Gal, galeterone.

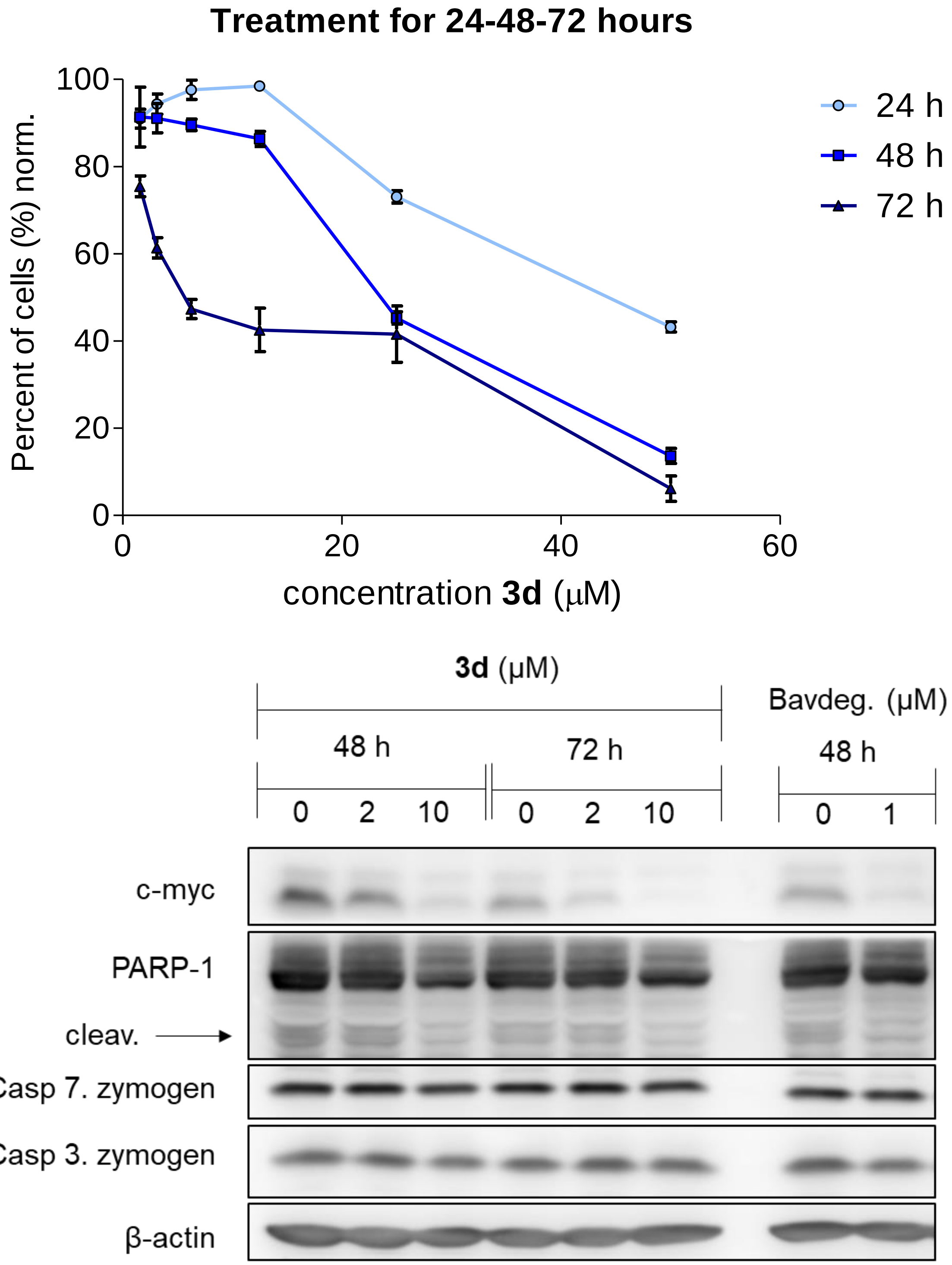


**Figure S7.** Antiproliferative activity of **3d** in LAPC-4 cell line after 24 h, 48 h and 72 h measured by resazurin-based viability assay. Protein levels of important proteins connected with proliferation and apoptosis in LAPC-4 treated by **3d** or bavdegalutamide for 48 h or 72, analogously to the treatment displayed in **Figure 5.** Level of β-actin served as loading control.

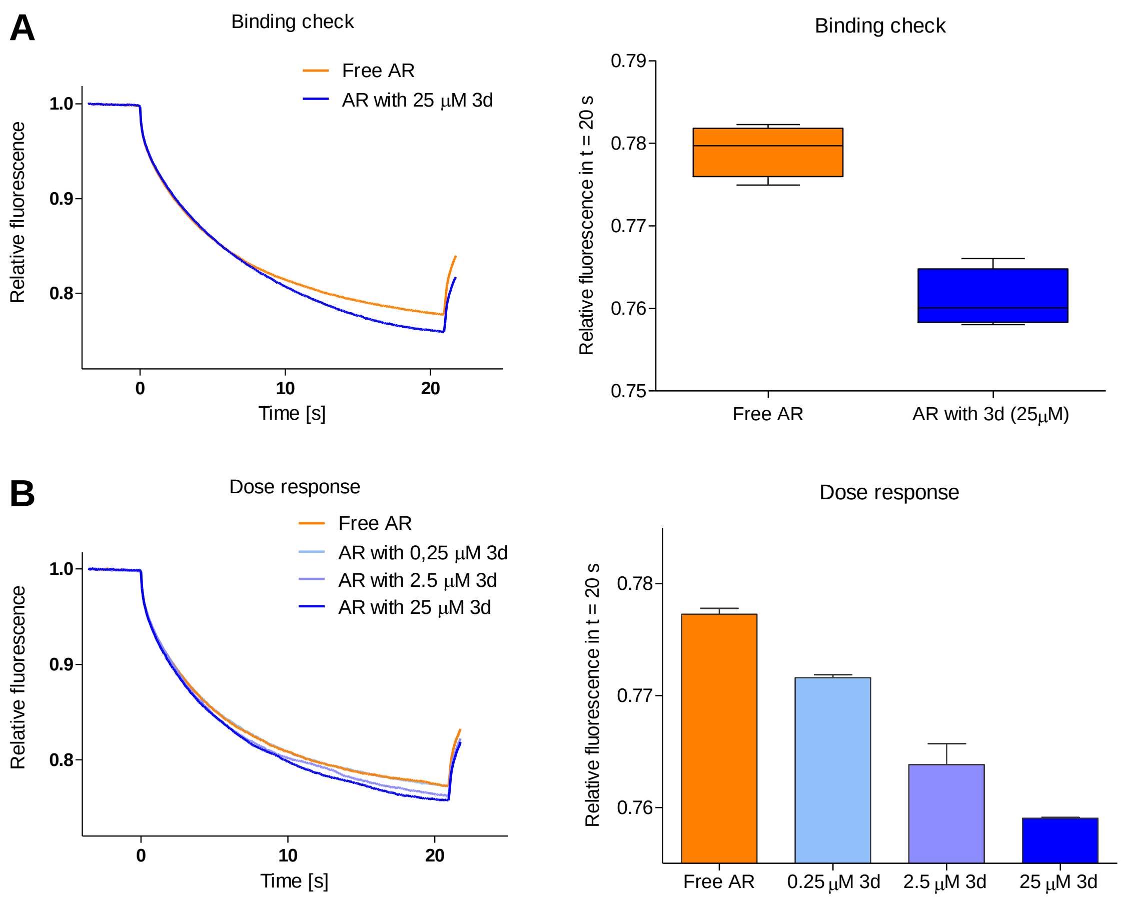


**Figure S8.** MST Measurement. Measured with recombinantly expressed and purified human AR-LBD, stained with NanoRed-NHS (Nanotemper). (A) Binding check measured with free AR and AR with 25 μM **3d**. Curves display the mean from (*n*= 4), while box plot display the mean and min. and max. value from (*n*= 4). (B) Dose response measurement was performed analogously, with free AR and 0.25 μM, 2.5 μM and 25 μM **3d**. Curves display the mean from (*n*= 2), while bar plot display the mean ± SD (*n*= 2).

**A**

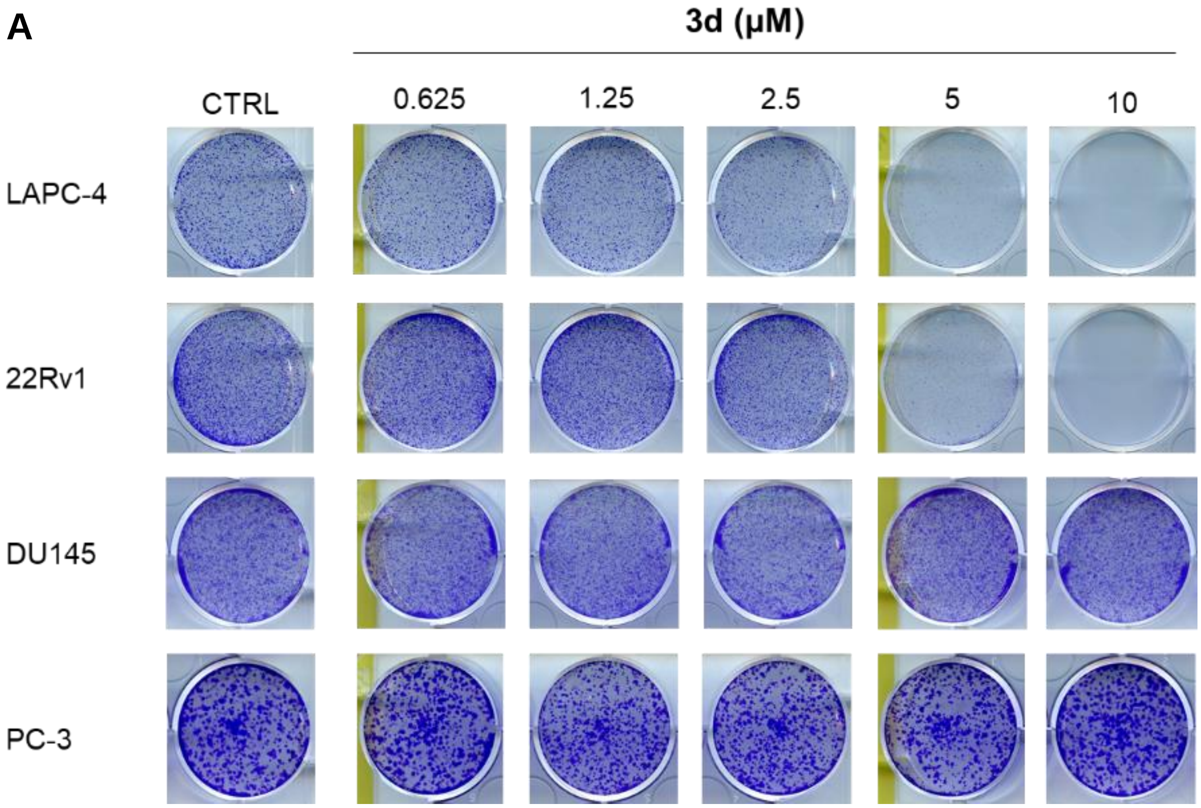


**B**

| Type | AR status* | Cell line | $GI_{50} \pm SD$ |
|---|---|---|---|
| Prostate | positive | LAPC-4 | 7.9 ± 1.6 |
| Prostate | positive | 22Rv1 | 18.3 ± 0.6 |
| Prostate | positive | LNCaP | 16.6 ± 0.8 |
| Prostate | positive | DuCaP | 22.9 ± 1.8 |
| Prostate | negative | DU145 | >50 |
| Prostate | negative | PC-3 | >50 |
| Breast | positive | T47D | 18.2 ± 0.3 |
| Breast | positive | SKBR3 | 25.5 ± 5.2 |
| Breast | positive | MCF7 | 38.0 ± 6.1 |

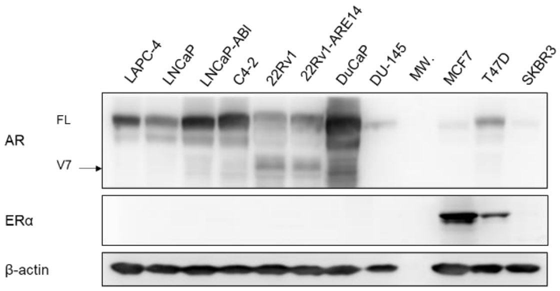


* based on literature and determined from the protein expression

**Figure S9.** Antiproliferative activity of **3d** determined by colony formation assay (A) or resazurin-based viability assay (B). In the table in panel B, protein levels of AR and ERα are shown, based on western blotting. Level of β-actin served as loading control. See experimental part for assay's detail.

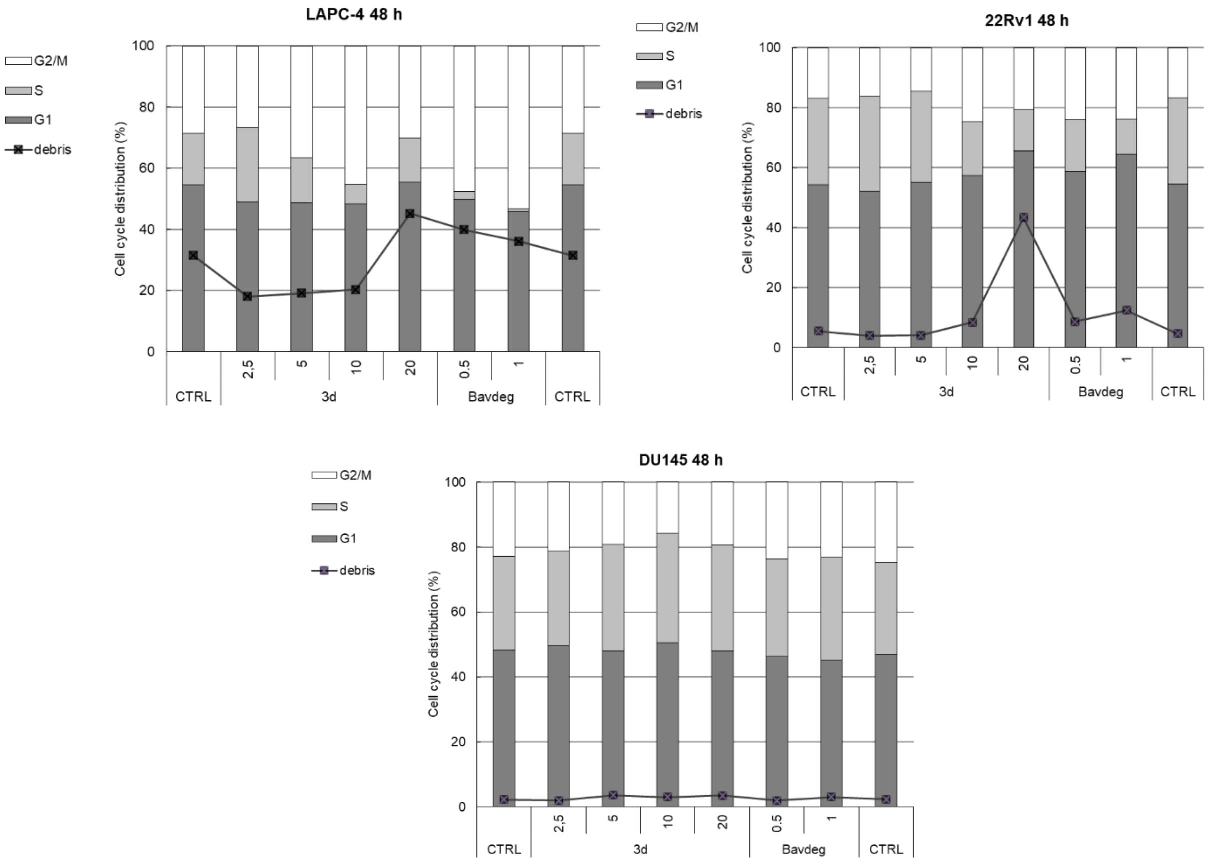


**Figure S10.** Cell cycle analysis of LAPC-4, 22Rv1 and DU145 after 48h treatment with **3d** or bavdegalutamide.

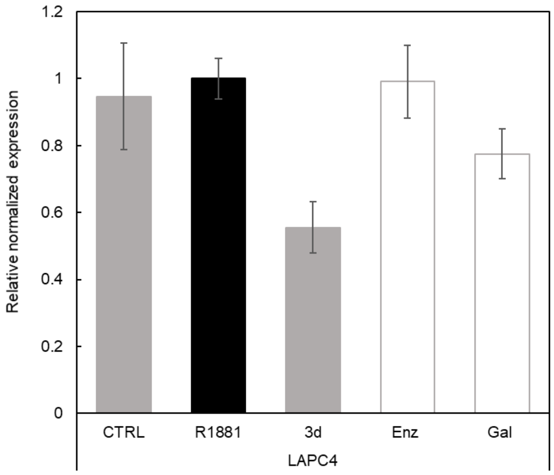


**Figure S11.** The effect of compound **3d** and standard AR antagonists on relative normalized expression of AR (full-length) in LAPC-4 cells. Cells were cultivated in CSS medium overnight, then treated with compounds in 10 μM concentration in presence of 1 nM R1881 for 24 h. Enz, enzalutamide; Gal, galeterone.

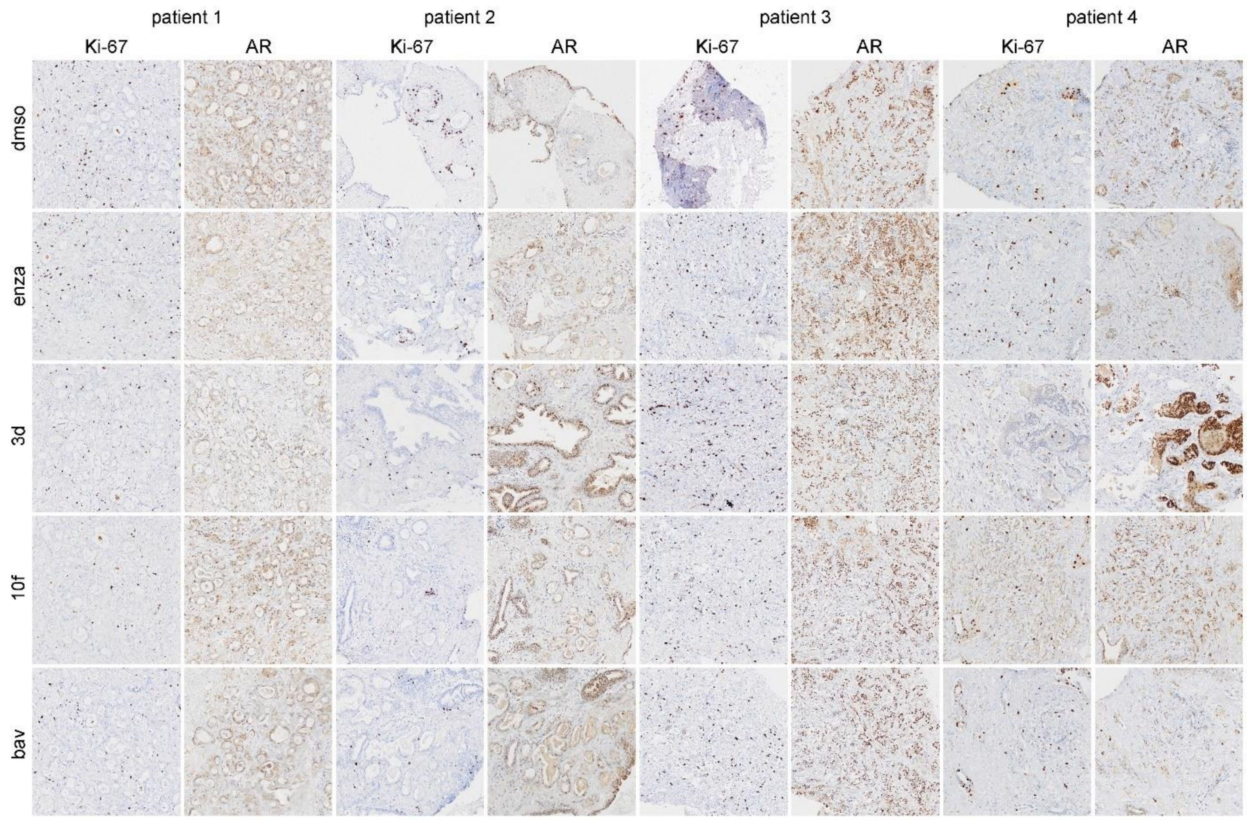


**Figure S12.** Immunohistochemistry analysis of Ki67 and AR in the *ex vivo* tissue culture experiment. The prostatectomy tissues were cut with vibratome, and slices were treated for three days with our candidates **3d** and **10f** (both 10 μM) along with enzalutamide and bavdegalutamide (both 1 μM). The tissues were then formalin-fixed, paraffin-embedded and processed for standard immunohistochemistry.
